# Repurposing UBE2W for programmable protein ubiquitylation

**DOI:** 10.1101/2025.09.22.676137

**Authors:** Paul Schnacke, Maximilian Fottner, Julian van Gerwen, Denys Kvasha, Fabian Willenborg, Pedro Beltrao, Kathrin Lang

## Abstract

Deciphering the ubiquitin code requires homogenous, site-specifically ubiquitylated proteins, yet access to such conjugates remains a major challenge. Existing approaches are often constrained by low yields, harsh reaction conditions, engineered recognition motifs or non-native linkage architectures. Here, we present UbyW (Ubiquitylation by UBE2W), a programmable platform for site-specific ubiquitylation that repurposes the E2 enzyme UBE2W to target genetically encoded isopeptidic neo-N-termini. UbyW enables efficient generation of near-native Ub-protein conjugates across diverse protein substrates, including endogenous ubiquitylation sites within folded domains, and can be implemented through a reconstituted intracellular cascade in *Escherichia coli* for streamlined high-yield production. The platform further enables installation of chemical functionalities adjacent to the isopeptidic linkage, including photocrosslinkers for capturing modification-dependent interactions. Using programmable probes targeting site-specific ubiquitylation of the small GTPase Ran, we identify USP15 as a cognate deubiquitylase and show that Ran K71 monoubiquitylation disrupts key Ran-cycle interactions.

## Introduction

Post-translational modification (PTM) of proteins with ubiquitin (Ub) regulates diverse cellular processes, including protein turnover, localization, signaling and stress responses. This functional diversity is orchestrated by a complex network, termed the “ubiquitin code”, in which modification site, Ub chain architecture and linkage topology collectively dictate distinct downstream signaling outcomes.^1,2^

Deciphering this code requires well-defined ubiquitin-protein of interest (Ub-POI) conjugates that enable direct interrogation of site-specific ubiquitylation events.^3^ However, access to such probes remains challenging. In particular, probes that preserve the native isopeptide linkage between Ub and the POI while enabling chemical functionalization at the ubiquitylation site would provide powerful tools for capturing transient and modification-dependent interactions.^4^

Over the past decades, substantial efforts have been devoted to the synthesis of ubiquitylated proteins using strategies such as native chemical ligation (NCL) and click chemistry.^5,6^ The required conjugation handles are typically introduced either by solid-phase peptide synthesis (SPPS) or recombinantly through genetic code expansion (GCE).^6,7^ Although powerful, these approaches have inherent limitations. SPPS is largely restricted to smaller, readily refoldable protein domains and NCL typically requires harsh, denaturing conditions. Click chemistry-based approaches, conversely, introduce non-native ligation scars that may perturb local structure and interaction profiles.^5^

To overcome these limitations, the field has increasingly adopted chemoenzymatic strategies that form native-like linkages under physiological conditions (Supplementary Fig. 1a). Methods such as LACE^8,9^ and SUE1^10^ utilize E2 enzymes to attach Ub to a target protein. However, these systems require installation of recognition tags of four or more amino acids into the POI, limiting access to endogenous modification sites. Sortylation, which relies on the transpeptidase Sortase 2A, addresses this constraint by selectively recognizing a non-canonical amino acid (ncAA) introduced through GCE.^11,12^ Yet, this approach requires mutations at the Ub C-terminus and is associated with comparatively low protein yields due to GCE inefficiencies. Additionally, direct incorporation of warheads or photocrosslinking handles at the linkage site has remained inaccessible with these methods.

To address these challenges, we present UbyW (Ubiquitylation by UBE2W), a chemoenzymatic platform for efficient and site-specific ubiquitylation (Supplementary Fig. 1b). UbyW exploits a previously underappreciated activity of the E2 conjugating enzyme UBE2W to selectively target neo-N-termini introduced into target proteins through incorporation of isopeptidic XisoK dipeptides via GCE. The resulting Ub-POI conjugates closely resemble their native counterparts and can be generated in high yield. UbyW operates both *in vitro* and within a reconstituted cascade in living *E. coli*, enabling straightforward production and single-step purification of complex Ub-POI conjugates. Crucially, this platform also permits direct installation of biorthogonal handles and photocrosslinkers adjacent to the isopeptidic linkage site, allowing direct interrogation of modification-dependent interactions. We demonstrate this capability by generating photocrosslinking probes for Ran K71 ubiquitylation, which enable us to show that Ub modification disrupts key Ran-cycle interactions and identify USP15 as a cognate deubiquitylase.

## Results

### Site-specific modification of XisoK-bearing target proteins by UBE2W

Our work builds on the recently developed G-XisoK toolbox, which enables efficient active uptake of G-XisoK tripeptidic non-canonical amino acids (ncAAs) into *E. coli* cells, facilitating genetic encoding of XisoKs at near wild-type expression levels using low G-XisoK concentrations (Supplementary Fig. 2).^13^ Intriguingly, XisoKs have recently been identified as naturally occurring PTMs formed through aminoacylation of the lysine N-ε by aminoacyl-tRNA synthetases (aaRS).^14^ Proteomic studies further suggest that these XisoK PTMs are ubiquitylated by the atypical E2 conjugating enzyme UBE2W.^15^ Inspired by these findings, we sought to establish a platform for site-specific, high-yield protein ubiquitylation by combining the XisoK toolbox with UBE2W.

To investigate whether UBE2W could mediate site-specific ubiquitylation of XisoK-bearing proteins, we first generated a model substrate based on SUMO2 bearing ʟ-leucine isopeptide-linked lysine (LisoK) at position 11 (SUMO2(K11LisoK)). Because UBE2W was previously reported to ubiquitylate the native N-terminus of SUMO2, we introduced an N-terminal proline residue followed by a TEV cleavage site to suppress modification at this potentially competing site. To our delight, initial incubation of SUMO2(K11LisoK) with UBE2W, Ub, and the E1 activating enzyme UBA1 *in vitro,* resulted in formation of the desired Ub-SUMO2(K11LisoK) conjugate, albeit with low efficiency (Fig. 1 a,b). We additionally observed pronounced UBE2W autoubiquitylation, a known characteristic of this enzyme.^16^

**Figure 1.**
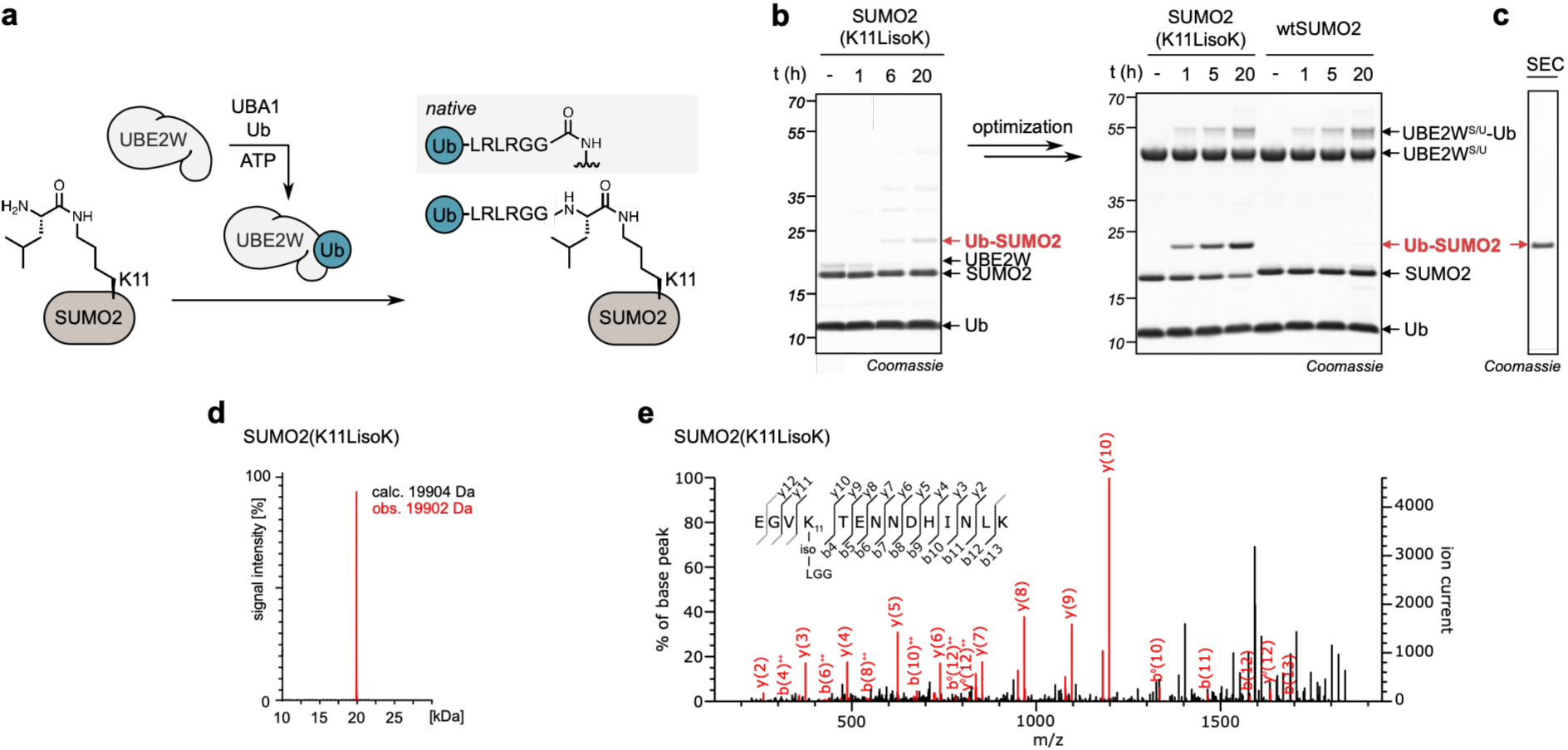
Repurposing UBE2W for programmable, high-yield protein ubiquitylation. **a,** SUMO2 bearing the non-canonical amino acid (ncAA) LisoK at K11 is site-specifically modified by UBE2W, yielding a near-native Ub-SUMO2 conjugate. **b,** Left: SDS-PAGE analysis of initial in vitro ubiquitylation of SUMO2(K11LisoK) with UBE2W, UBA1, and Ub (20 μM SUMO2(K11LisoK) with 5 μM UBE2W, 50 μM Ub, and 100 nM of the E1 activating), Right: SDS-PAGE analysis of optimized modification of SUMO2(K11LisoK) using UBE2W^S/U^ (40 μM SUMO2(K11LisoK), 40 μM UBE2W^S/U^, 100 μM Ub, 100 nM UBA1). **c,** SDS-PAGE analysis of purified Ub-SUMO2(K11LisoK). **d**, LC-MS analysis confirms successful conjugation. **e,** MS-MS analysis of Ub-SUMO2(K11LisoK) confirms site-specific attachment of Ub to LisoK. Full gels are shown in Supplementary Fig. 4. Consistent results were obtained over three distinct replicate experiments.

To enhance product yield, we systematically optimized the reaction conditions. By adjusting the reaction buffer and stoichiometry of the reaction components, and by generating a Strep-SUMO2-Ub-UBE2W fusion construct, termed UBE2W^S/U^, we reduced UBE2W autoubiquitylation and increased conversion more than five-fold. Optimized reaction conditions comprising 40 μM SUMO2(K11LisoK), 40 μM UBE2W^S/U^, 100 μM Ub, 100 nM UBA1 resulted in over 50% product formation within 20 hours at 37°C and pH 7.1. Importantly, no ubiquitylation was observed for wild-type SUMO2 as a substrate, confirming the reaction’s dependence on the neo-N-terminus within LisoK (Fig. 1b, Supplementary Figs 3,4). The resulting Ub-SUMO2(K11LisoK) conjugate was readily purified to homogeneity by Ni-NTA affinity chromatography, TEV protease cleavage and size-exclusion chromatography (SEC) (Fig. 1c, Supplementary Fig. 4). The identity and site-specificity of the modification were confirmed by intact protein mass spectrometry (ESI-LC-MS) and tandem mass spectrometry (MS/MS), which identified a peptide fragment corresponding to Ub linked specifically to the LisoK residue (Fig. 1d, e, Supplementary Fig. 4). We term this platform UbyW (Ubiquitylation by UBE2W).

### Exploring UBE2W’s donor substrates and XisoK permissiveness

Next, we investigated the permissiveness of UbyW towards different Ub donor substrates. Linear (M1) di-, tri-, and tetraUb chains were conjugated to SUMO2(K11LisoK) with efficiencies comparable to monoUb, and conjugation yield was independent of Ub chain length (Fig. 2a, Supplementary Fig. 5a). We further tested UBE2W-mediated attachment of a K63-linked diUb to LisoK at position 11 within SUMO2. This conjugate also formed successfully, albeit with slightly reduced efficiency compared to the linear chains (Fig. 2b, Supplementary Fig. 5b). Together, these results demonstrate that UBE2W accepts a broad range of Ub donor substrates, from monoUb to polyUb chains of defined length and linkage type associated with distinct signaling pathways.^17^

**Figure 2.**
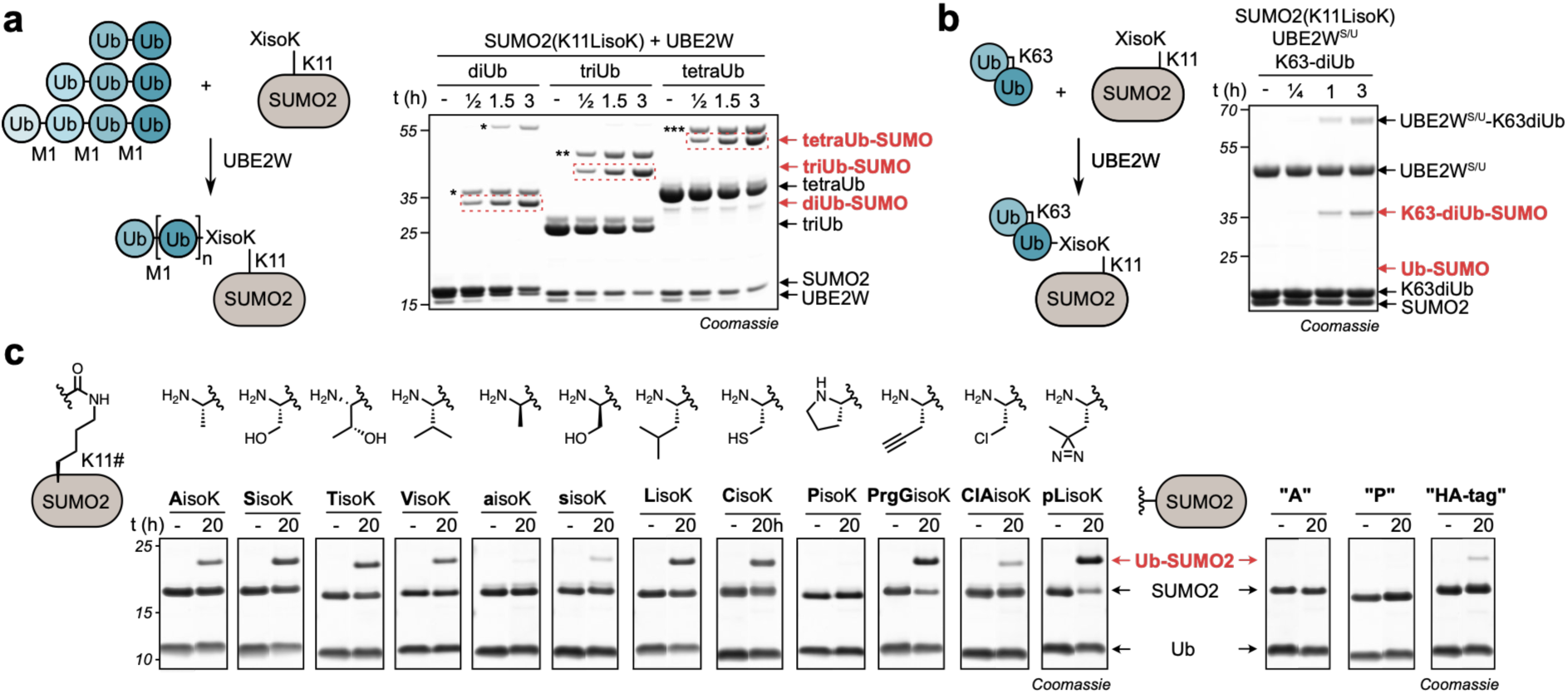
UBE2W-mediated polyUb chain attachment and XisoK permissiveness. **a,** SDS-PAGE analysis of the attachment of linear (M1) Ub polymers onto SUMO2(K11LisoK). Red dashed boxes highlight the formation of the respective polyUb-SUMO2 conjugates. Asterisks (*, **, ***) denote the corresponding autoubiquitylated UBE2W species. **b,** SDS-PAGE analysis of the attachment of K63-linked diUb onto SUMO2(K11LisoK). Consistent results were obtained over three distinct replicate experiments. **c,** SDS-PAGE analysis of UBE2W-mediated modification of SUMO2-K11 bearing different XisoKs at K11 (left) and different amino acid at the unstructured N-termini (right). Full gels and reaction conditions are shown in Supplementary Fig. 8. Consistent results were obtained over three distinct replicate experiments.

Building on this, we then investigated how the identity of the X residue within the XisoK-modified acceptor protein influences UBE2W reactivity. A key advantage of the XisoK toolbox is its compatibility with a broad range of amino acid side chains.^13^ We therefore generated a panel of SUMO2(K11XisoK) variants bearing small (A, S), aliphatic (V, L), nucleophilic (C), and conformationally restricted (P) ʟ-amino acids, as well as ᴅ-amino acids (a, s) (Fig. 2c and Supplementary Figs 6-8). Incubation of these SUMO2 XisoK variants under optimized conditions revealed a strong preference for primary ʟ-amino acids, with high conversion observed for A, S, T, V, L, and C (Fig. 2c, Supplementary Fig. 8). In contrast, proline was a poor substrate, and ᴅ-amino acids were modified only to a limited extent, indicating a strict stereochemical requirement for the reaction. For comparison, we also tested SUMO2 constructs bearing different N-termini under identical reaction conditions, including mature SUMO2 with its native cleaved N-terminal alanine (denoted ‘A’), an HA-tagged variant (denoted ‘HA’, with a starting methionine), and an N-terminal proline construct (MP-TEV-SUMO2, denoted ‘P’). Although ‘A’ represents the authentic wild-type sequence, we employ the ‘P’ construct as the standard control (referred to as wtSUMO2 in all other assays) to suppress background N-terminal ubiquitylation. SUMO2 variants bearing either the native N-terminal alanine or an HA-tag have previously been characterized as UBE2W substrates,^16,18^ but both were substantially poorer substrates in our assay than the ʟ-XisoK variants (Fig. 2c, Supplementary Fig. 8). Together, these findings suggest that UBE2W may preferentially recognizes and ubiquitylates isopeptidically modified lysines over native N-termini.

Finally, we investigated whether UbyW is compatible with direct installation of reactive handles at the ubiquitylation site. To this end, we incorporated XisoK scaffolds bearing bioorthogonal handles (PrgG), chemical crosslinkers (ClA) or photocrosslinkers (pL) as the X residue. Notably, all three variants were efficiently modified, demonstrating that UbyW can generate Ub-POI conjugates bearing defined chemical handles adjacent to the isopeptidic linkage (Fig. 2c, Supplementary Fig. 8). These conjugates are therefore ideally suited for downstream conjugation, trapping and crosslinking applications.

### Ubiquitylation of structured protein domains at endogenous modification sites

With the broad scope of compatible XisoK acceptors established, we next investigated the permissiveness of UBE2W towards structurally and biologically diverse target proteins. We found that both Histone H3 and the microtubule-associated protein Tau can be ubiquitylated by UBE2W at endogenous modification sites (H3(K27) and Tau(244-372, K353), Fig. 3a, b, Supplementary Fig. 9). We also successfully ubiquitylated α-synuclein at K10 and K21 (Fig. 3c, Supplementary Fig. 9). Together, these results demonstrate that UbyW is compatible with diverse protein sequence contexts beyond SUMO2(K11).

**Figure 3.**
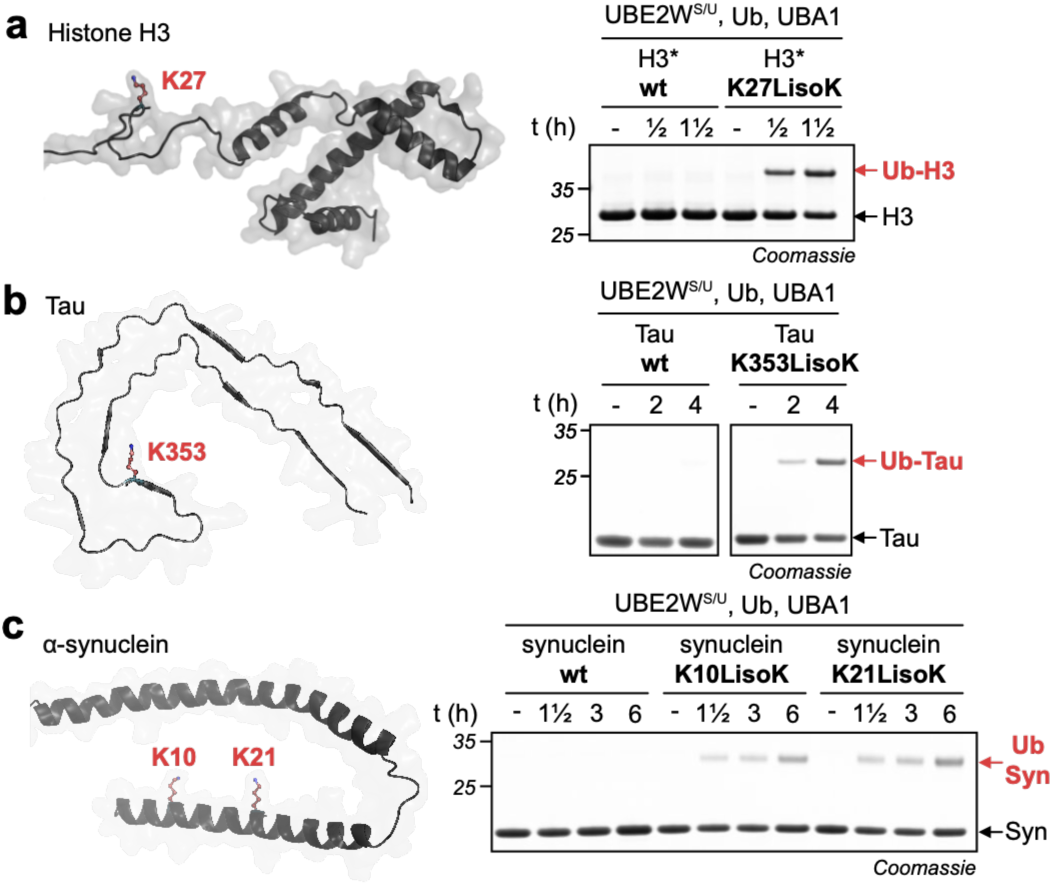
UbyW enables site-specific ubiquitylation across diverse protein folds. **a,** Left: Structure of Histone H3 with K27 highlighted in red (Protein Data Bank, 1kx5)^32^. Right: SDS-PAGE analysis of *in vitro* modification of H3*(K27LisoK). The asterisk (*) indicates an N-terminal ubiquitin fusion to enhance H3 solubility. **b,** Left: Structure of tau with K353 highlighted in red (Protein Data Bank, 6hrf)^33^. Right: SDS-PAGE analysis of *in vitro* modification of tau(244-372, K353LisoK). wt-tau(244-372) is not modified. **c,** Left: Structure of α-synuclein with K10 and K21 highlighted in red (Protein Data Bank, 1xq8)^34^. Right: SDS-PAGE analysis of *in vitro* modification of α-synuclein(K10LisoK), and α-synuclein(K21LisoK). Full gels reaction conditions are shown in Supplementary Fig. 9. Consistent results were obtained over three distinct replicate experiments.

However, modification of more complex proteins, such as homotrimeric PCNA, proved substantially more challenging and resulted in low *in vitro* conjugation yields (Supplementary Fig. 10). Beyond poor efficiency, the *in vitro* workflow is experimentally demanding, requiring independent expression and purification of all reaction components, including UBA1, UBE2W^S/U^, Ub, and the XisoK-bearing target protein, followed by an additional purification step to isolate the final conjugate. Prolonged *in vitro* incubation may further increase the risk of protein degradation.

### An intracellular UbyW cascade enables high-yield production of Ub-POI conjugates

To address the limitations of *in vitro* UbyW, we next asked whether the entire ubiquitylation cascade, together with the required GCE machinery, could be reconstituted in living *E. coli* (Fig. 4a, Supplementary Fig. 11). To this end, we co-expressed SUMO2 bearing an amber codon at position K11 with the corresponding aaRS/tRNA pair (PylRS/PylT). Upon supplementation of the culture medium with G-XisoK, the tripeptide is actively taken up by *E. coli* and proteolytically processed, enabling XisoK incorporation into the target protein. In parallel, we co-expressed the remaining UbyW components, allowing UBA1 to charge UBE2W with Ub for subsequent ubiquitylation of the XisoK-containing SUMO2 substrate.

**Figure 4.**
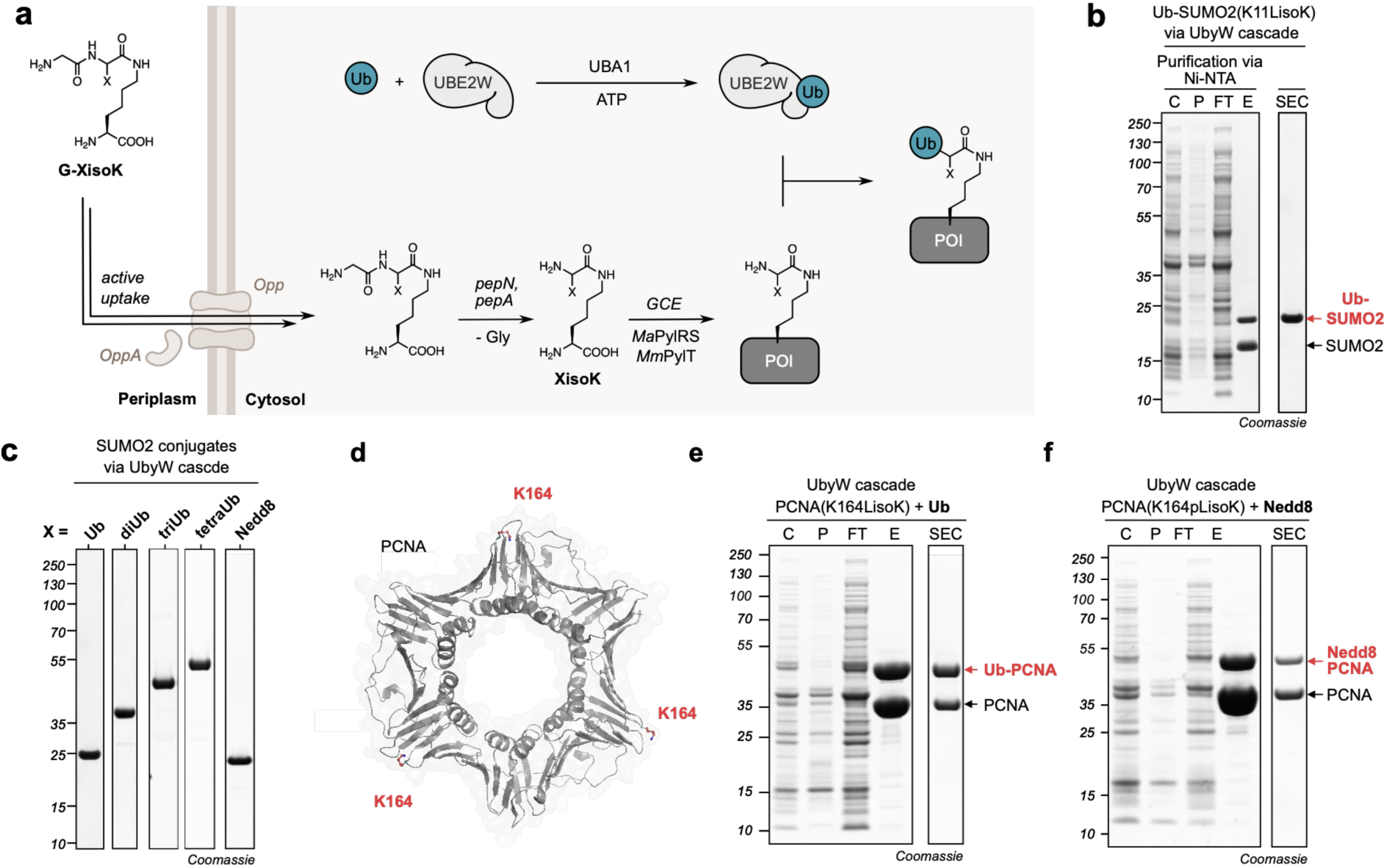
*In cellulo* UbyW enables site-specific modification of complex proteins in *E. coli*. **a,** Schematic of the UbyW cascade. Active uptake of G-XisoK into *E. coli* via Opp and subsequent proteolytic cleavage by pepN and pepA enable high-yield amber suppression. Co-expressed UBE2W is charged with Ub by UBA1 and modifies the XisoK-bearing target protein directly in *E. coli.* **b,** SDS-PAGE analysis of Ni-NTA purification and size exclusion chromatography of the Ub-SUMO2(K11LisoK) conjugate generated with the reconstituted cascade using *H. sapiens* UBE2W. C, P, FT, and E indicate culture, pellet, flowthrough, and elution fractions, respectively. **c** SDS-PAGE analysis of purified SUMO2 conjugates, including M1-linked polyUb chains and Nedd8, generated via *in cellulo* UbyW using *H. sapiens* UBE2W. **d,** Structure of the PCNA homo-trimer with the K164 modification site highlighted in red (Protein Data Bank, 1axc).^35^ **e,** SDS-PAGE analysis of the purification of the Ub-PCNA(K164LisoK) conjugate generated via the UbyW cascade. **f,** SDS-PAGE analysis of the purification of the Nedd8-PCNA(K164pLisoK) conjugate generated via the UbyW cascade. Full gels reaction conditions are shown in Supplementary Figs. 14-16. Consistent results were obtained over three distinct replicate experiments.

Initial experiments yielded detectable but low amounts of the ubiquitylated product (Supplementary Fig. 11a). We therefore optimized the *in cellulo* setup by consolidating the genetic components onto fewer plasmids and testing different promoters to balance expression levels of the cascade components. This resulted in a more than ten-fold improvement in isolated conjugate yield (Supplementary Fig. 11 b-d). Using this optimized UbyW system, we isolated 3.1 mg/L purified Ub-SUMO2(K11LisoK) conjugate directly from cultures supplemented with as little as 0.5 mM of G-LisoK (Fig. 4b) after a single affinity purification and size-exclusion chromatography step. The identity of the conjugate was confirmed by mass spectrometry (Supplementary Fig. 11e, 13). We termed this intracellular ubiquitylation platform the ‘UbyW cascade’.

This cascade proved highly versatile and enabled direct production of SUMO2 conjugates bearing linear M1-linked di-, tri-, and tetraUb chains in *E. coli*, reaching isolated yields up to 4.8 mg/L (Fig. Supplementary Fig. 11f-h, 13). Importantly, the system also tolerated modifications to the Ub C-terminus. We successfully conjugated a Ub variant carrying an L73P mutation to SUMO2(K11LisoK), generating conjugates resistant to deubiquitylase (DUB) cleavage both *in vitro* and in HEK293T cell lysate. These DUB-stable Ub-POI conjugates are therefore useful for mammalian lysate-based assays and cellular delivery experiments. (Supplementary Fig. 12a, c, d, 13).^19^

Notably, the UbyW cascade was also compatible with other Ub-like modifiers (Ubls).^20^ We demonstrated efficient conjugation of Nedd8, a Ubl whose recombinant production and conjugation are often low-yielding and experimentally demanding (Fig. 4c, Supplementary Fig. 12b, 13). These results highlight the advantage of the *in cellulo* cascade for direct generation of challenging Ubl-POI conjugates.

To investigate whether additional protein acceptor scaffolds are permissive to the cascade, we next applied *in cellulo* UbyW to SUMO1 and the essential NF-κB regulator NEMO (Supplementary Fig. 14). We achieved site-specific ubiquitylation of SUMO1 at K7 and NEMO at K302 and K309. Notably, both NEMO lysine residues are positioned within the α-helical UBAN domain, a structural motif proposed to regulate NEMO activity in the NF-κB pathway.^21^ This demonstrates that UbyW can modify sites embedded within folded secondary structures without requiring installation of extended recognition sequences around the target residue, distinguishing it from methods that depend on engineered pepetide motifs and are therefore limited in their ability to target endogenous modification sites.^9,10^

Given the high degree of conservation of UBE2W across different species, we investigated whether homologs from other organisms could also function in the intracellular E. coli cascade and potentially further improve conjugate formation.^22^ We tested a panel of UBE2W homologs and found that ubiquitylation activity was broadly conserved, although isolated product yields varied between species (Supplementary Fig. 15a-c). Notably, the *D. rerio* and *C. elegans* homologs outperformed human UBE2W both *in vitro* and *in cellulo*, increasing the yield of isolated Ub-POI conjugates (Supplementary Fig. 15d). For subsequent applications, we therefore employed the highly active *D. rerio* homolog as an optimized variant of the UbyW cascade.

Having established and optimized the UbyW cascade, we revisited position K164 of PCNA, a central Ubl modification site within the homo-trimeric DNA sliding clamp that coordinates replication-associated DNA damage responses and had previously resisted efficient modification *in vitro* (Supplementary Fig. 10, 16). The optimized cascade enabled production of Ub-PCNA(K164LisoK) in high yield (35 mg/L, Fig. 4e). Moreover, the system also supported Nedd8ylation of PCNA at the same site, providing access to a native but still poorly characterized modification implicated in DNA repair regulation (Fig. 4f). Both conjugates were confirmed by LC-MS. Together, these results establish the UbyW cascade as a streamlined intracellular platform for producing site-specifically modified Ub- and Ubl-POI conjugates combining flexibility in the Ub/Ubl donor architecture with compatibility across structurally complex protein acceptors.

### Functional Profiling of Ran K71 Ubiquitylation using UbyW-derived Probes

To demonstrate the utility of UbyW for uncovering regulatory mechanisms, we next sought to investigate a site-specific ubiquitylation event of high biological interest. Although proteomic studies have identified thousands of ubiquitylation sites, the functional relevance of most individual modifications remains unknown. To prioritize candidate sites, we utilized a recently developed machine learning-based positional importance score for ubiquitylation sites.^23^ Derived from harmonized, pan-species ubiquitin proteomics data across various cellular perturbations, this metric predicts functionally important, site-specific ubiquitylation events and enriches for non-degradative signaling functions.^23^

This analysis identified lysine 71 (K71) of the small GTPase Ran as a prime candidate, ranking within the top percentile of all evaluated human ubiquitylation sites and scoring highest among all sites identified in Ran itself (Fig. 5a). Furthermore, these harmonized ubiquitin proteomics data indicate that K71 ubiquitylation is highly dynamic, being upregulated in response to cellular stressors and downregulated upon proteasome inhibition (Fig. 5b). This response pattern suggests a targeted regulatory signaling function rather than a general degradation-associated modification.

**Figure 5.**
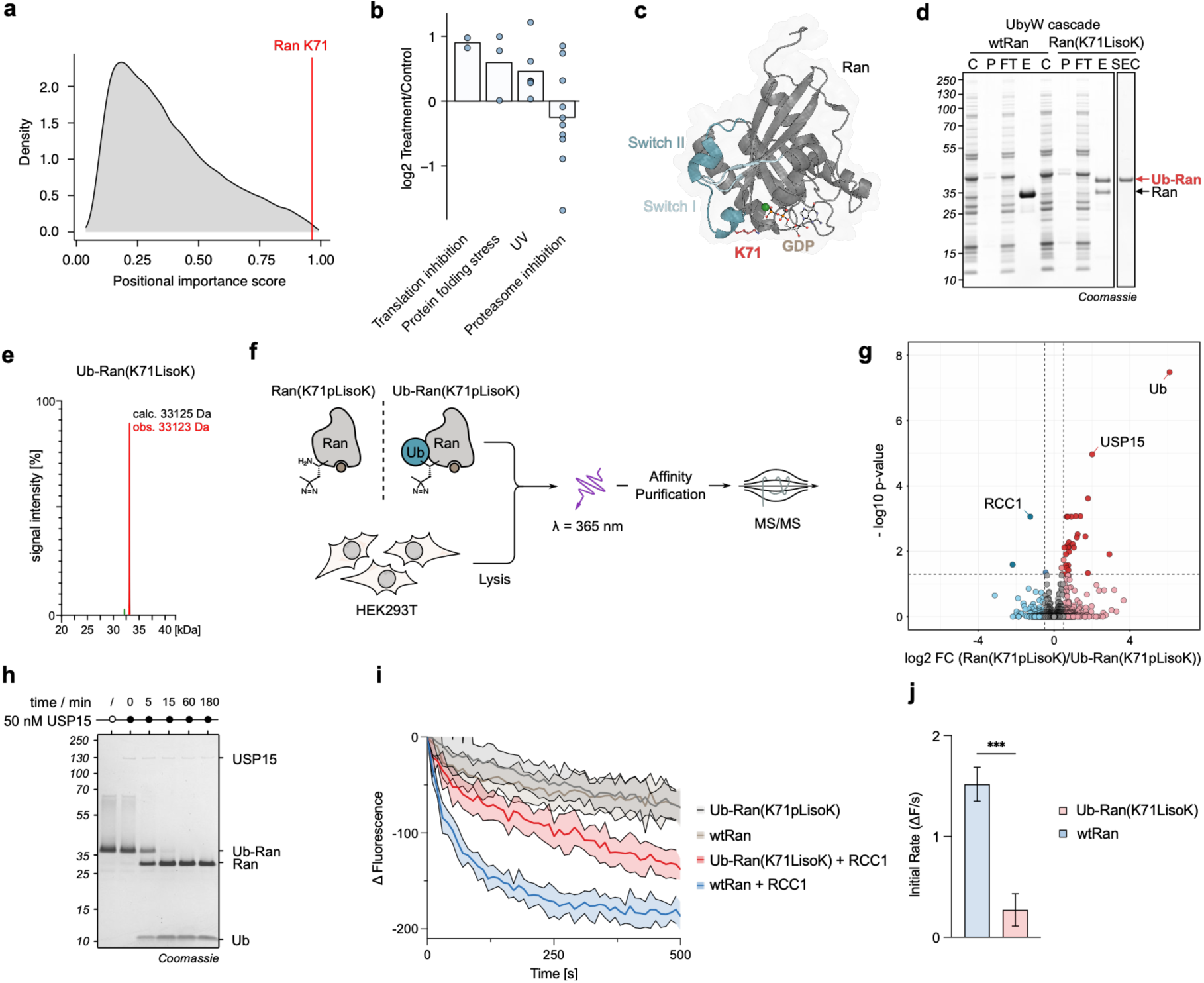
Generation and functional profiling of site-specifically ubiquitylated Ran(K71). **a,** Density plot showing the overall distribution of the ubiquitylation site positional importance score. The relative position of Ran(K71) is highlighted. **b,** Log_2_ fold changes of Ran(K71) ubiquitylation across distinct cellular stress conditions derived from ubiquitin proteomics data. **c,** Structure of the small GTPase Ran bound to GDP (Protein Data Bank, 3gj0).^24^ The K71 modification site is highlighted in red. Co-crystallized GDP and Mg^2+^ (green) are shown alongside the Switch I (light blue) and Switch II (teal) domains. **d,** SDS-PAGE analysis of the purification of the Ub-Ran(K71LisoK) conjugate generated in *E. coli.* **e,** LC-MS analysis confirming the successful generation of the Ub-Ran(K71LisoK) conjugate. **f,** Schematic workflow of the crosslinking AP-MS approach. Ran(K71pLisoK), Ub-Ran(K71pLisoK), or no-probe controls were incubated in HEK293T cell lysate, followed by UV irradiation to crosslink interacting proteins, affinity purification, and identification via LC-MS/MS. Consistent results were obtained over three distinct replicate experiments. **g,** Volcano plot comparing protein enrichment between the Ran(K71pLisoK) and Ub-Ran(K71pLisoK) probes. **h,** *In vitro* deubiquitylation assay. Ub-Ran(K71pLisoK) (10 µM) was incubated with the AP-MS hit USP15 (50 nM) at 37°C, and cleavage was monitored over time via SDS-PAGE. Full gels are shown in Supplementary Fig. 19. **i,** RCC1-mediated nucleotide exchange assay. Wild-type Ran (wtRan) and Ub-Ran(K71LisoK) were loaded with Mant-GDP. Nucleotide exchange in the presence of RCC1 was monitored continuously via changes in Mant-GDP fluorescence. **j,** Initial nucleotide exchange rates (v_0_) calculated from the linear phase (0-50 s) of the reactions shown in (i). Error bars represent the standard error of the regression slope (SEM, n = 5 independent replicates). Statistical significance was determined using an unpaired Welch’s t-test (*** P = 0.0007 vs. wtRan). The Ub-Ran exchange rate is not significantly different from zero (ns, P = 0.1033, one-sample t-test).

Structurally, K71 is positioned within the dynamic Switch II domain of Ran (Fig. 5c).^24^ Ran regulates nucleocytoplasmic transport by cycling between GDP- and GTP-bound states, with the Switch II domain forming a critical interface for nucleotide-dependent interactors, including the transport factor NTF2, the exchange factor RCC1.^25,26^ Although K71 acetylation has previously been studied, the functional consequences of Ran K71 ubiquitylation remain unknown. Together with its predicted functional importance and dynamic regulation, this made Ran K71 an attractive target for functional profiling using the UbyW platform.

To investigate the functional consequences of K71 ubiquitylation, we first attempted to generate the modified protein *in vitro*. However, Ran exhibited poor stability under these conditions, and prolonged incubation of Ran(K71LisoK) resulted in severe degradation (Supplementary Fig. 17a). We therefore turned to the UbyW cascade, which yielded intact, Ub-Ran(K71LisoK) conjugate with >50% conversion and no detectable degradation (Fig. 5d, Supplementary Fig. 17b, c). LC-MS analysis confirmed formation of the defined conjugate (Fig. 5e, Supplementary Fig. 17d). This result highlights the practical value of the intracellular UbyW cascade for target proteins that are unstable during *in vitro* modification and therefore difficult to access using standard *in vitro* chemoenzymatic methods.

To map the K71 ubiquitin-dependent interactome and identify candidate regulators of this modification, including potential DUBs, we next leveraged the acceptor-site permissiveness of UbyW to generate matched non-ubiquitylated and ubiquitylated Ran photocrosslinking probes. Specifically, the X residue within XisoK was replaced with diazirine-containing photoleucine (pL), enabling UV-induced covalent trapping of interacting proteins (Supplementary Fig. 17e-g). The resulting probes (Ran(K71pLisoK) and Ub-Ran(K71pLisoK)) were applied in a label-free quantitative affinity purification mass spectrometry (LFQ-AP-MS) workflow in HEK293T cell lysates (Fig. 5f). Following UV irradiation, probe-bound proteins were affinity-purified and analyzed by MS/MS. Across the dataset, an average of 7,800 proteins was identified with a median coefficient of variation (CV) of 18.0% (Supplementary Fig. S18).

To evaluate probe performance, we first examined their global interaction profiles. Both probes enriched a robust network of known Ran interactors relative to the control, with a particular enrichment for Ran•GDP-specific binders (Supplementary Fig. S19a), confirming capture of the Ran interactome. However, we noted the absence of the canonical Ran•GDP binder NTF2. Because K71 acetylation has previously been shown to disrupt NTF2 binding,^27^ we reasoned that modification of the K71 side chain itself might preclude this interaction. Indeed, *in vitro* pulldowns confirmed that only wild-type Ran efficiently captures NTF2, whereas both Ran(K71LisoK) and Ub-Ran(K71LisoK) failed to bind NTF2 (Supplementary Fig. S19c), indicating that even minimal modification at K71 disrupts the Ran-NTF2 interface.

With the broader Ran interactome validated, we next compared Ran(K71pLisoK) and Ub-Ran(K71pLisoK) to uncover distinct ubiquitin-dependent interaction changes (Fig. 5g, Supplementary Fig. 19b). Alongside the expected enrichment of ubiquitin itself, USP15 emerged as one of the most strongly enriched proteins in the Ub-Ran(K71pLisoK) samples, identifying it as a candidate regulator of this PTM (Supplementary Fig. S19d). In contrast, the Ran guanine nucleotide exchange factor (GEF) RCC1 was among the most strongly depleted proteins upon K71 ubiquitylation.

Having disentangled the general effects of K71 side-chain modification (loss of NTF2) from the specific consequences of the ubiquitin appendage (USP15 recruitment and RCC1 depletion), we next investigated the functional roles of the identified hits. USP15 is a versatile cysteine protease that shuttles between the cytoplasm and the nucleus and is implicated in diverse signaling pathways.^28^ To determine whether USP15 directly processes the K71-linked conjugate, we performed an *in vitro* deubiquitylation assay. Co-incubation of purified Ub-Ran(K71LisoK) with recombinant USP15 resulted in rapid cleavage of the Ub moiety, with an estimated conjugate half-life of 5 minutes under the tested reaction conditions (Fig. 5h). These findings establish USP15 as an eraser of Ran K71 ubiquitylation, providing a mechanism for dynamic reversal of this modification.

In contrast to USP15 recruitment, the strong depletion of RCC1 suggests that K71 ubiquitylation impairs the interaction between Ran and its primary GEF. RCC1 catalyzes GDP to GTP exchange on Ran within the nucleus and thereby maintains directionality of nuclear transport.^29^ To investigate the functional consequences of K71 ubiquitylation on this process, we performed real-time nucleotide exchange assays using Ran probes loaded with the fluorescent nucleotide analog Mant-GDP. While wtRan exhibited robust RCC1-stimulated exchange, ubiquitylation at K71 markedly reduced the exchange rate. Notably, nucleotide exchange of Ub-Ran in the presence of RCC1 was statistically indistinguishable from spontaneous baseline exchange observed in the absence of RCC1 (Fig. 5i, Supplementary Figs 19e, f). Together, these findings demonstrate that K71-ubiquitylated Ran is an incompetent substrate for RCC1, indicating that this modification blocks RCC1-mediated nucleotide exchange and may thereby disrupt the Ran cycle.

## Discussion

In this study, we introduce UbyW (Ubiquitylation by UBE2W), a chemoenzymatic platform for site-specific protein ubiquitylation. UbyW enables modification of diverse protein substrates across endogenous ubiquitylation sites and is compatible with both unstructured regions and folded domains. To overcome the limitations of conventional *in vitro* approaches, we further developed an intracellular UbyW cascade in *E. coli*, enabling high-yield generation of challenging conjugates such as PCNA and Ran through straightforward co-transformation and standard purification workflows, without requiring specialized expertise in chemical synthesis or protein semi-synthesis.

The resulting conjugates closely mimic their native counterparts. In contrast to existing E2-based approaches, such as LACE or SUE1,^9,10^ UbyW does not require installation of extended recognition motifs within the target protein sequence. Instead, the conjugates differ only by a single ‘X’ residue derived from the XisoK handle, which introduces a minimal spacer between the Ub C-terminus and the target lysine in the POI, while preserving the unperturbed Ub core and its native interaction surfaces. Importantly, the ‘X’ residue provides a particularly advantageous position for installing chemical handles directly adjacent to the isopeptidic linkage, including photocrosslinkers for covalent capture of modification-dependent interactors.^30^

Mechanistically, UbyW exploits a previously underappreciated activity of UBE2W towards isopeptidic neo-N-termini. Our data reveal a pronounced preference for XisoK over native protein N-termini, alongside broad tolerance for diverse L-amino acid chains but poor acceptance of D-amino acids. These observations point to a highly stereoselective recognition mechanism that remains poorly understood. Unlike canonical E2 enzymes, UBE2W lacks the conserved gatekeeper aspartate residue and contains an HPH rather than HPN motif (Supplementary Fig. 20), features that may contribute to its unusual α-over ε-amine substrate selectivity.^31^ However, these structural differences alone do not explain the striking preference for isopeptidic neo-N-termini, which may additionally involve the enzyme’s flexible C-terminus, previously implicated in substrate recognition.^18^ Notably, UBE2W displays minimal dependence on local sequence context (Supplementary Fig. 20), enabling broad applicability of UbyW across diverse protein substrates. More broadly, the evolutionary conservation of this E3-independent activity raises the possibility that UBE2W contributes to the regulation and turnover of naturally occurring XisoK PTMs *in vivo*.^15,22^

Owing to its modularity and chemical programmability, UbyW enables functional interrogation of poorly characterized ubiquitylation sites. Using site-specifically modified photocrosslinking Ran probes, we show that K71 ubiquitylation disrupts interactions of Ran with both NTF2 and RCC1, which are essential for the cyto-nuclear transport cycle. At the same time, we identify USP15 as a potent eraser of this modification site, supporting a model in which Ran K71 ubiquitylation acts as a dynamic and reversible signaling event rather than a mark for proteasomal degradation. Consistent with this interpretation, ubiquitin proteomics data indicate stress-dependent regulation of K71 ubiquitylation.^23^ Future work will define the cellular and subcellular contexts and states in which this modification operates and investigate its interplay with K71 acetylation and SUMOylation.

The Ran case study illustrates how UbyW can connect predictive bioinformatic prioritization of ubiquitylation sites with direct biochemical validation. More generally, combining such predictions with the UbyW platform provides a scalable strategy for systematic functional interrogation of the Ub code. By overcoming bottlenecks of traditional synthetic and chemoenzymatic ubiquitylation methods and enabling streamlined access to native-like Ub-protein conjugates, including chemically functionalized probes, UbyW transforms previously inaccessible ubiquitylation sites into tractable targets for biochemical, structural and mechanistic investigation.

## Notes

### Competing Interest Statement

K.L, M.F. and Tarun Iype are named as inventors on a pending patent application EP24205590.3. All other authors do not declare any competing interests.

### Summary of Updates

This revised version of the manuscript contains photocrosslinking and protoemics data on the use of our Ub-probes

### Funding

- European Research Council: ERC under the European Union’s Horizon 2020 research and innovation program, grant agreement no. 101003289–Ubl-tool to K.L

