## Supplementary Information for "Repurposing UBE2W for programmable protein ubiquitylation"

*3.8 Expression and purification of H6-tagged wt proteins (SUMO2, PCNA, tau, synuclein, Ub-H3, and Ran)*.. 49

### Supplementary Figures

#### Supplementary Figure 1

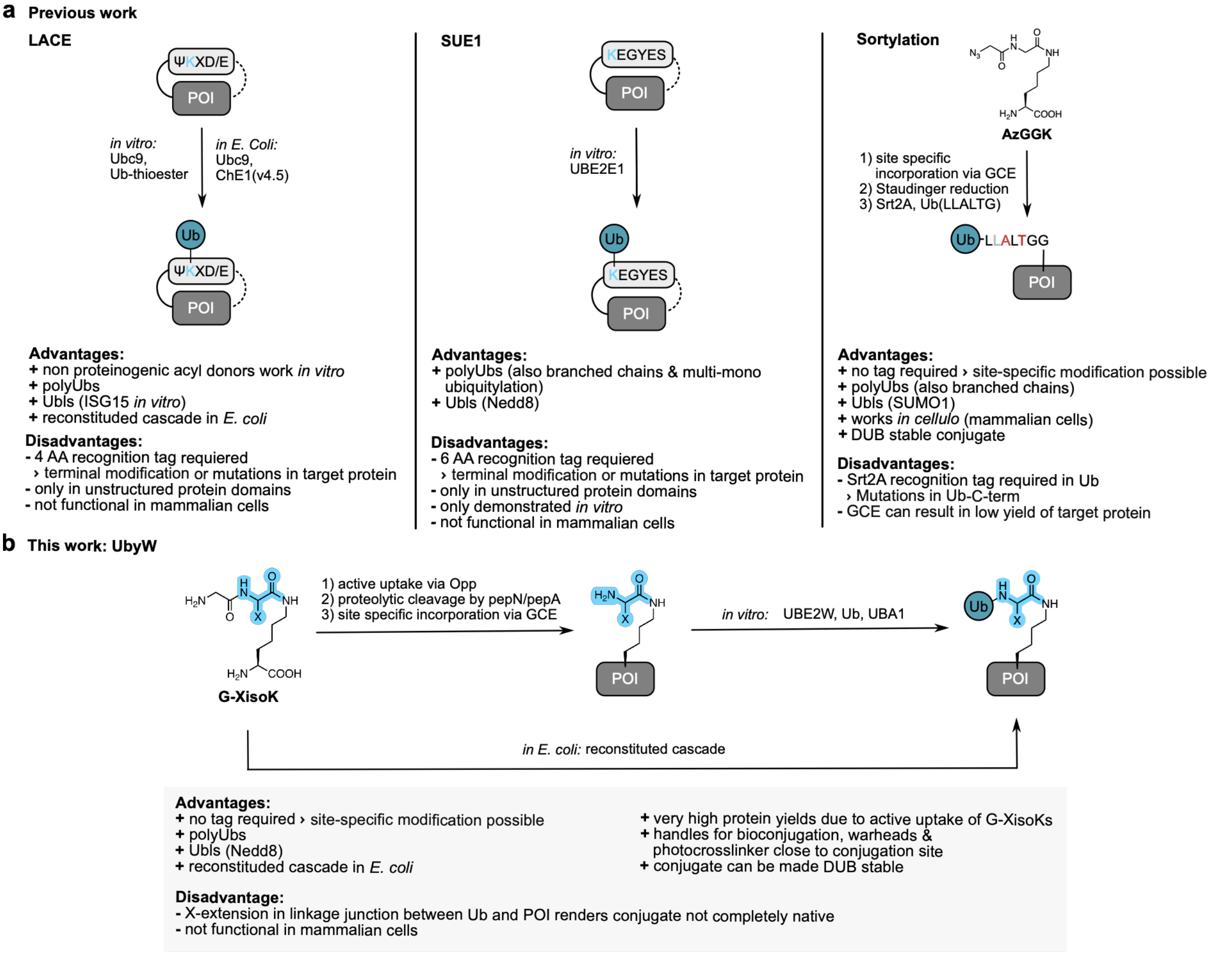

**Supplementary Figure S1. Chemoenzymatic methods for site specific ubiquitylation. a,** Previous methods for ubiquitylation: Left: Lysine Acylation using Conjugating Enzymes (LACE), Middle: Sequence-dependent Ubiquitination using UBE2E1 (SUE1), Right: Sortylation: site-specific, Srt2A catalyzed ubiquitylation. **b,** Ubiquitylation by UBE2W (UbyW) described in this manuscript.

**Supplementary Figure 2**

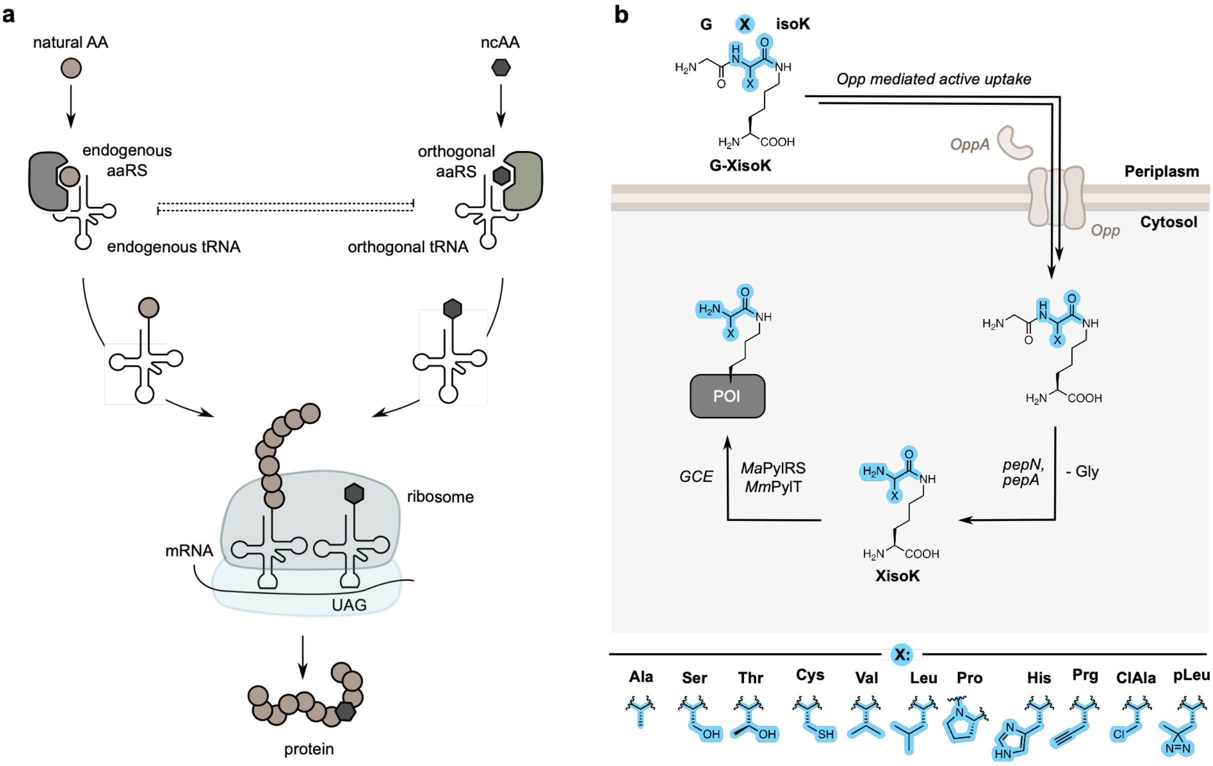

**Supplementary Figure S2. Working principle of genetic code expansion and the XisoK toolbox. a,** Genetic code expansion (GCE) by amber suppression is based on the employment of aminoacyl tRNA Synthetases (aaRS) and their cognate tRNAs that are orthogonal to endogenous aaRS. The orthogonal aaRS can charge its tRNA with a non-canonical amino acid (ncAA). The ncAA is co-translationally incorporated into a protein in response to the amber stop codon (UAG).[1] **b,** XisoK toolbox allows for expression of amber suppressed protein in wt-like yield. G-XisoK tripeptides are actively transported into *E. coli* by ABC transporter Opp. Subsequent proteolytic processing by pepN/pepA yields XisoK that is incorporated into proteins via GCE. Below: Structures of different XisoK amino acids, that are actively taken up and incorporated by leveraging Opp.[2]

**Supplementary Figure 3**

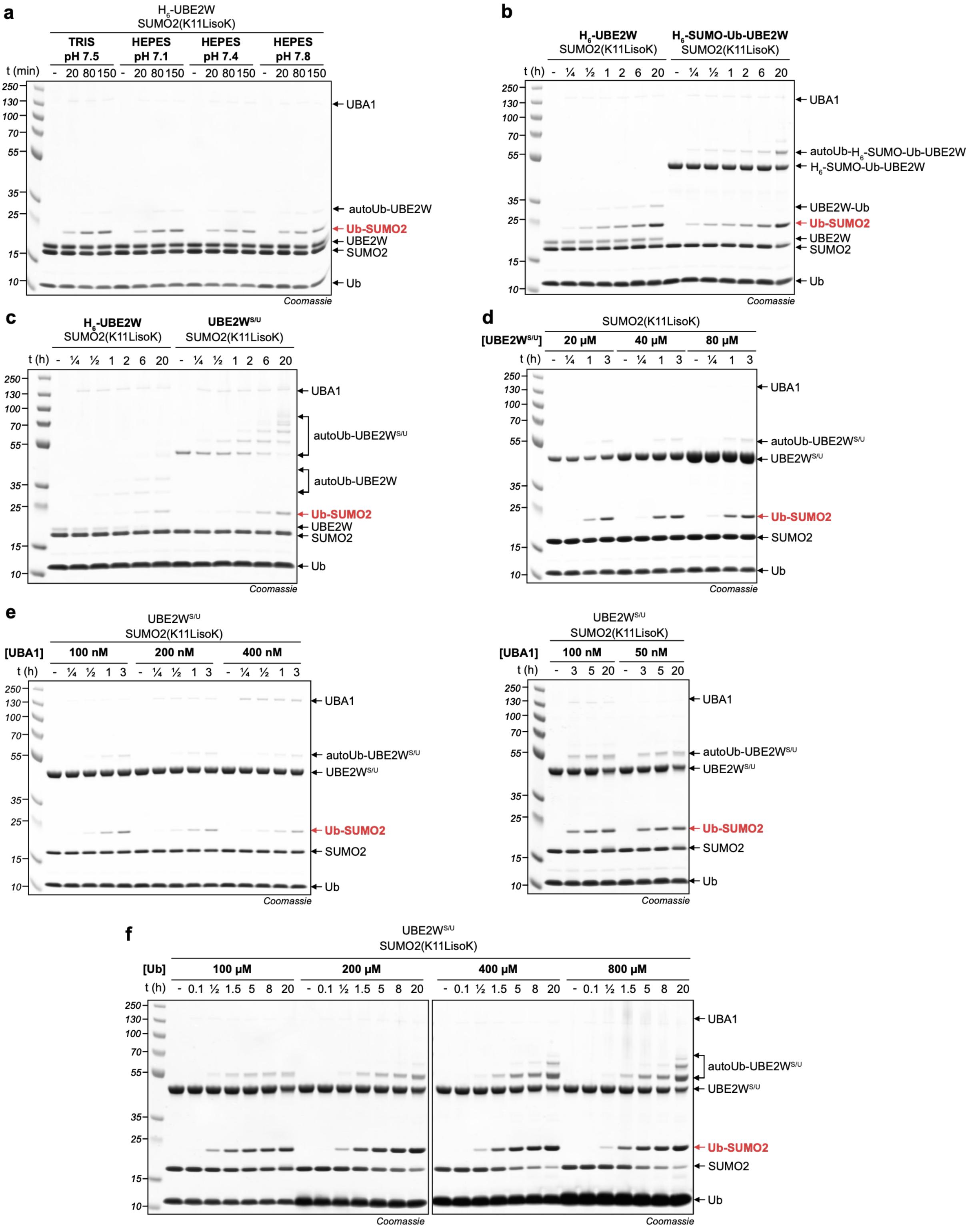

**Supplementary Figure S3. Optimization of in vitro reaction conditions and constructs for the modification of SUMO2(K11LisoK) with Ub.** Reaction progression is analyzed by SDS-PAGE**. a,** Variation of reaction buffer pH. Reaction conditions: 40 µM SUMO2(K11LisoK), 40 µM UBE2W, 60 µM Ub, 100 nM UBA1, 1 mM DTT, 10 mM ATP, 10 mM $MgCl_2$, 37°C. **b**, Comparison of $H_6$-UBE2W and $H_6$-SUMO-Ub-UBE2W. Reaction conditions: 40 µM SUMO2(K11LisoK), 20 µM UBE2W,

100 µM Ub, 100 nM UBA1, 50 mM TRIS pH 7.5, 1 mM DTT, 10 mM ATP, 10 mM $MgCl_2$, 37°C. **c**, Comparison $H_6$-UBE2W and Strep-SUMO-Ub-UBE2W ($UBE2W^{S/U}$). Reaction conditions: 20 µM SUMO2(K11LisoK), 5 µM UBE2W, 50 µM Ub, 100 nM UBA1, 50 mM TRIS pH 7.5, 1 mM DTT, 10 mM ATP, 10 mM $MgCl_2$, 37°C. **d**, Titration of Strep-SUMO-Ub-UBE2W ($UBE2W^{S/U}$). Reaction conditions: 80 µM SUMO2(K11LisoK), 100 µM Ub, 50 nM UBA1, 50 mM HEPES pH 7.1, 1 mM DTT, 100 mM NaCl, 10 mM ATP, 10 mM $MgCl_2$, 37°C. **e**, Titration of UBA1. Reaction conditions: 40 µM SUMO2(K11LisoK), 40 µM UBE2W, 100 µM Ub, 50 mM HEPES pH 7.1, 1 mM DTT, 100 mM NaCl, 10 mM ATP, 10 mM MgCl, 37°C. **f**, Titration of Ub. Reaction conditions: 40 µM SUMO2(K11LisoK), 40 µM UBE2W, 100 nM UBA1, 50 mM HEPES pH 7.1, 1 mM DTT, 100 mM NaCl, 10 mM ATP, 10 mM $MgCl_2$, 37°C.

**Supplementary Figure 4**

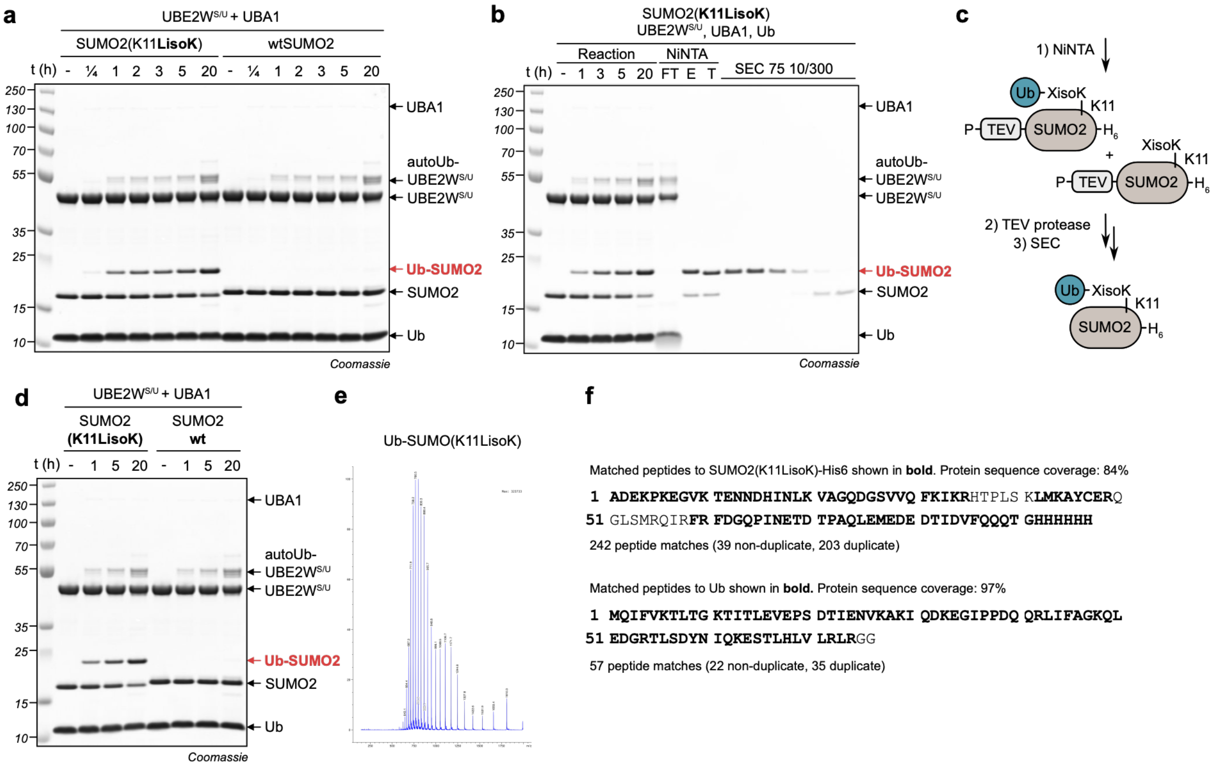

**Supplementary Figure S4. In vitro generation, purification, and mass spectrometry validation of Ub-SUMO2(K11LisoK).** Full SDS-PAGE gels of **a,** Modification of SUMO2(K11LisoK) and wtSUMO2 by UBE2W$^{S/U}$ *in vitro*. Reaction conditions: 40 µM SUMO2(K11LisoK), 20 µM UBE2W$^{S/U}$, 150 µM Ub, 50 nM UBA1, 50 mM HEPES pH 7.1, 1 mM DTT, 100 mM NaCl, 10 mM ATP, 10 mM $MgCl_2$, 37°C. **b,** Modification, purification by Ni-NTA and size exclusion chromatography of Ub-SUMO2(K11LisoK). Reaction conditions: 40 µM SUMO2(K11LisoK), 20 µM UBE2W$^{S/U}$, 150 µM Ub, 50 nM UBA1, 50 mM HEPES pH 7.1, 1 mM DTT, 100 mM NaCl, 10 mM ATP, 10 mM $MgCl_2$, 37°C. **c,** Purification scheme for Ub-SUMO2(K11LisoK). Initial Ni-NTA affords both conjugated and unconjugated protein, subsequent TEV proteolysis removes N-terminal blocking proline residue, and at last SEC yields pure conjugate. **d,** Modification, purification by Ni-NTA and size exclusion chromatography of SUMO2(K11LisoK) modified by UBE2W$^{S/U}$ with Ub. Reaction conditions: 40 µM SUMO2(K11LisoK), 20 µM UBE2W$^{S/U}$, 150 µM Ub, 50 nM UBA1, 50 mM HEPES pH 7.1, 1 mM DTT, 100 mM NaCl, 10 mM ATP, 10 mM $MgCl_2$, 37°C. **e,** ESI-MS m/z spectrum of Ub-SUMO2(K11LisoK) in native charge state distribution (non-deconvoluted), deconvoluted MS shown in Fig. 1c. **f,** Peptide coverage map of tryptic digest and MSMS of Ub-SUMO2(K11LisoK)-$H_6$, generated using Mascot.[3]

**Supplementary Figure 5**

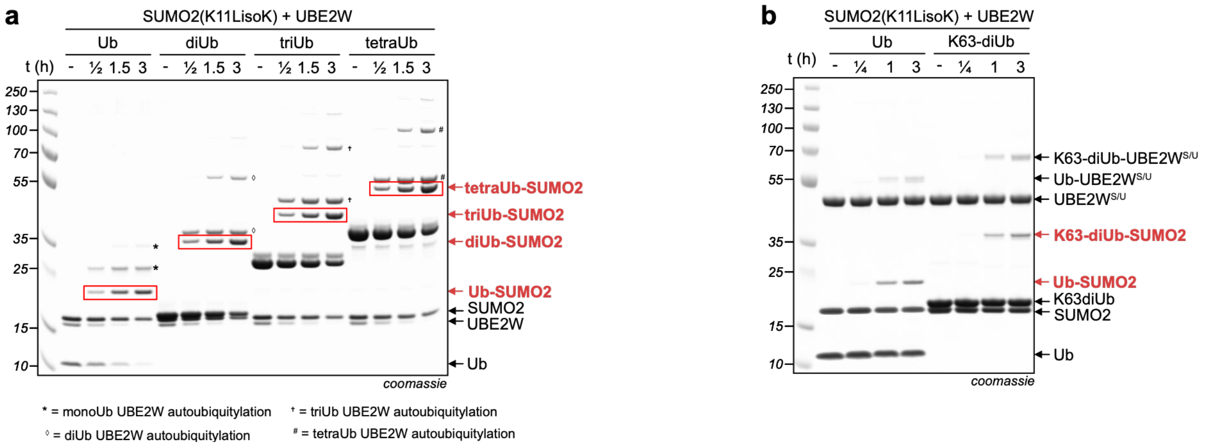

**Supplementary Figure S5. Full SDS-PAGE gels of modification of SUMO2(K11LisoK) with polyUb chains in vitro. a**, SDS-PAGE analysis of linear (M1) polyUb chain attachment to SUMO2(K11LisoK). Symbols denote the corresponding autoubiquitylated UBE2W species: asterisk (*), monoUb; diamond (◊), diUb; dagger (†), triUb; and hash (#), tetraUb. Reaction conditions: 40 μM SUMO2(K11LisoK), 20 μM UBE2W, 60 μM Ub/diUb(M1)/triUb(M1)/tetraUb(M1), 100 nM UBA1, 50 mM HEPES pH 7.1, 1 mM DTT, 100 mM NaCl, 10 mM ATP, 10 mM MgCl, 37°C. **b**, Modification with Ub, and K63-diUb. Reaction conditions: 40 μM SUMO2(K11LisoK), 40 μM UBE2W$^{S/U}$, 100 μM Ub/diUb(K63), 50 nM UBA1, 50 mM HEPES pH 7.1, 1 mM DTT, 100 mM NaCl, 10 mM ATP, 10 mM MgCl, 37°C.

**Supplementary Figure 6**

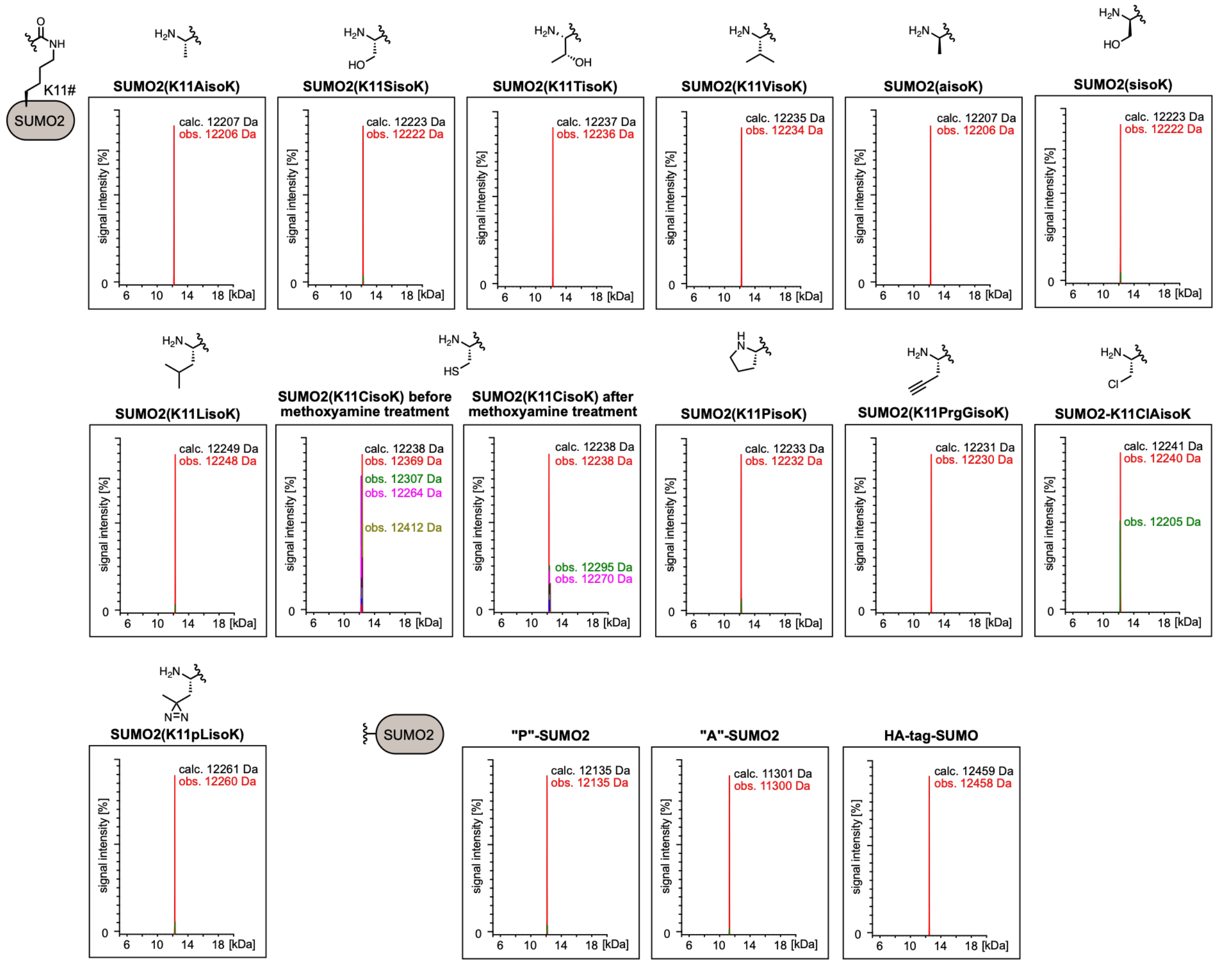

**Supplementary Figure S6. LC-MS analysis of SUMO2 bearing different XisoKs at K11.** Representative deconvoluted electrospray ionization mass spectra showing observed and calculated masses. Observed masses confirm incorporation of the corresponding XisoK derivatives. Non-deconvoluted m/z spectra are shown in Supplementary Figure 7. SUMO2(K11CisoK) was treated with methoxyamine to remove metabolic adducts. Peak denoted in green for SUMO2(K11ClAisoK) corresponds to the elimination of HCl.

### Supplementary Figure 7

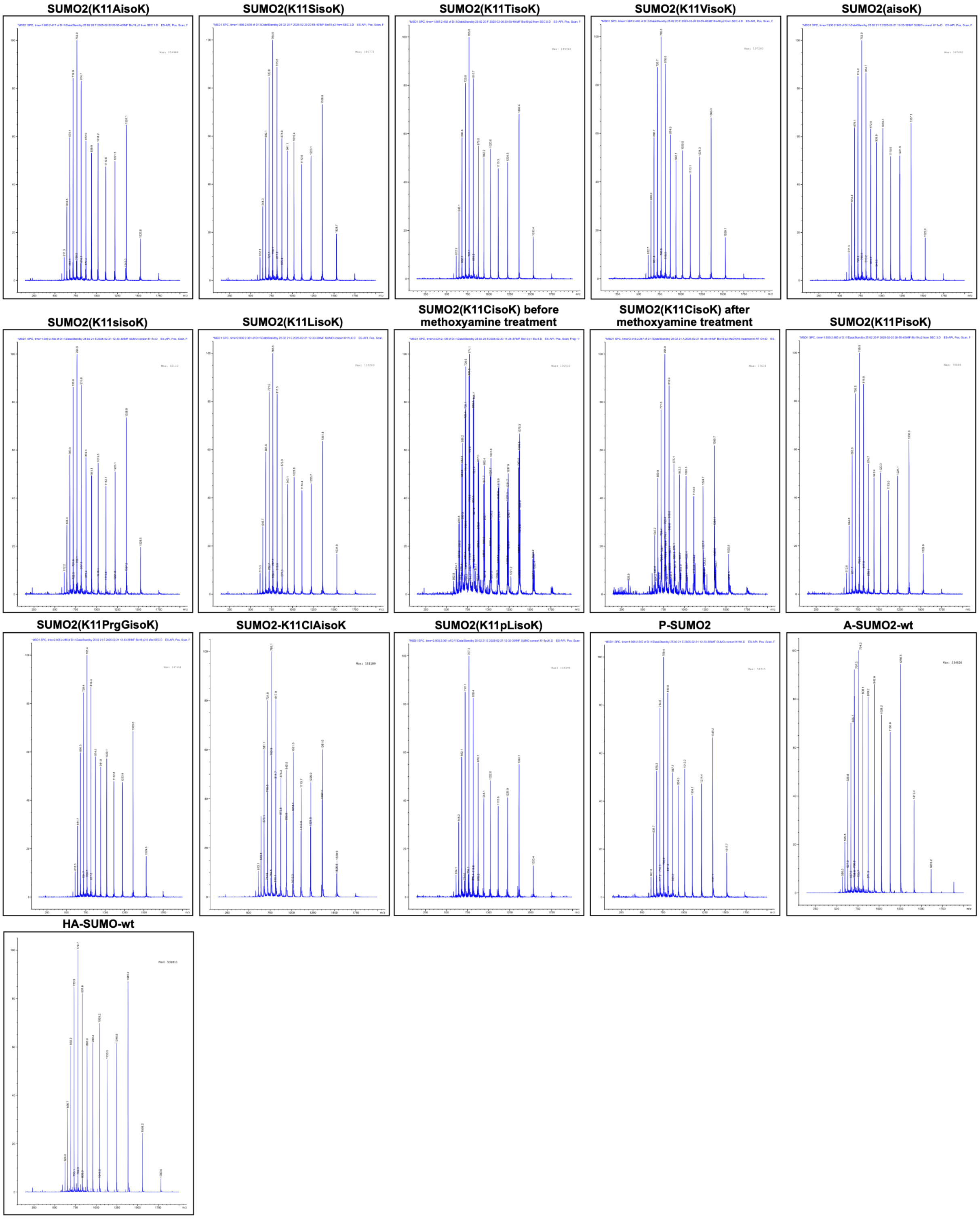

**Supplementary Figure S7. Raw ESI-MS spectra of SUMO2 bearing diverse XisoKs at K11.** ESI-MS m/z spectrum of SUMO2 bearing different XisoKs at K11 in native charge state distribution (non-deconvoluted). Deconvoluted masses are shown in Supplementary Figure 6.

**Supplementary Figure 8**

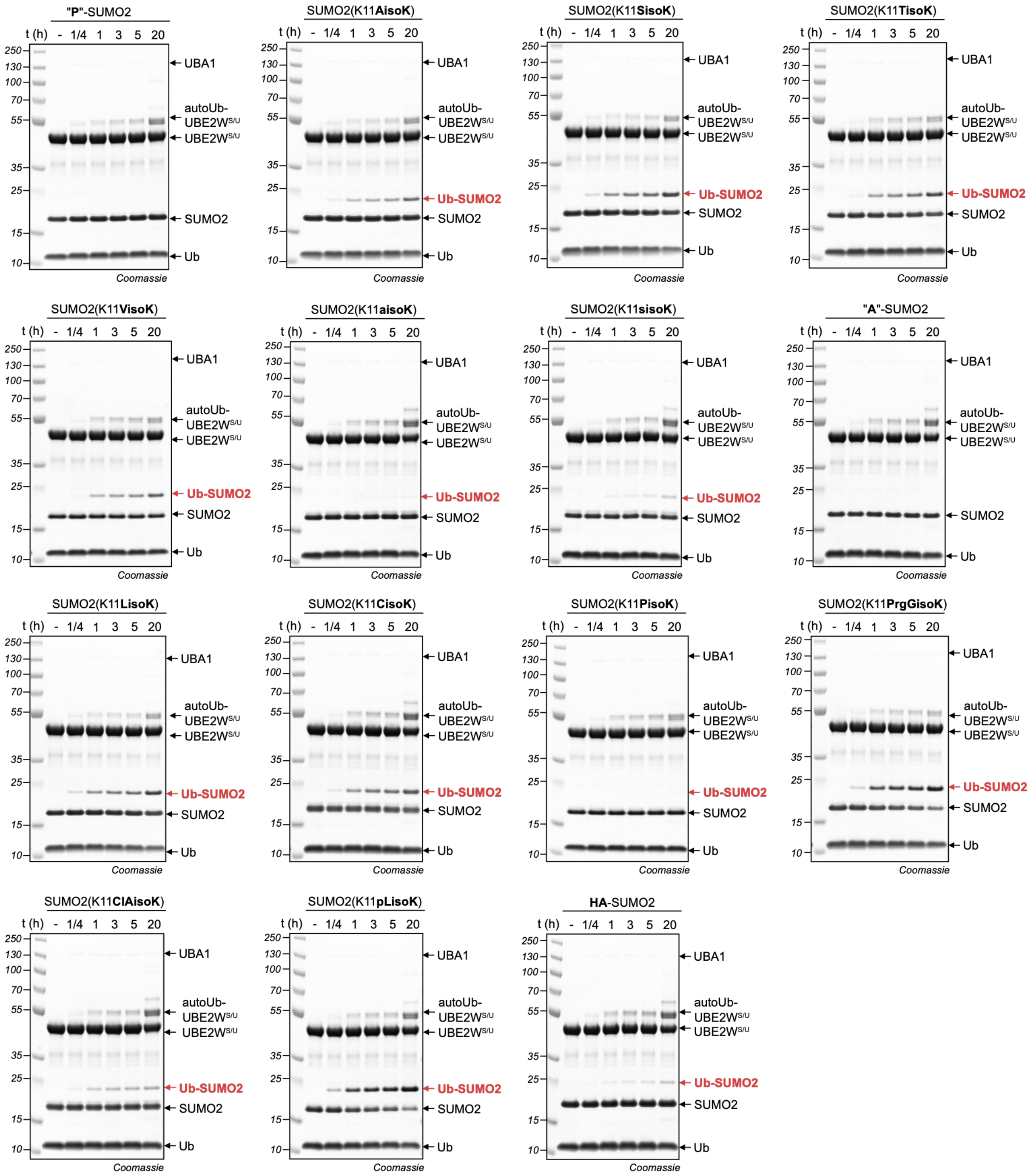

**Supplementary Figure S8. Full SDS-PAGE gels evaluating the XisoK substrate scope.** Full SDS-PAGE gels of reaction progression *in vitro* of SUMO2 bearing different XisoKs at K11 being modified with Ub. Reaction conditions: 40 μM SUMO2(K11LisoK), 40 μM UBE2W$^{S/U}$, 100 μM Ub, 50 nM UBA1, 50 mM HEPES pH 7.1, 1 mM DTT, 100 mM NaCl, 10 mM ATP, 10 mM $MgCl_2$, 37°C.

**Supplementary Figure 9**

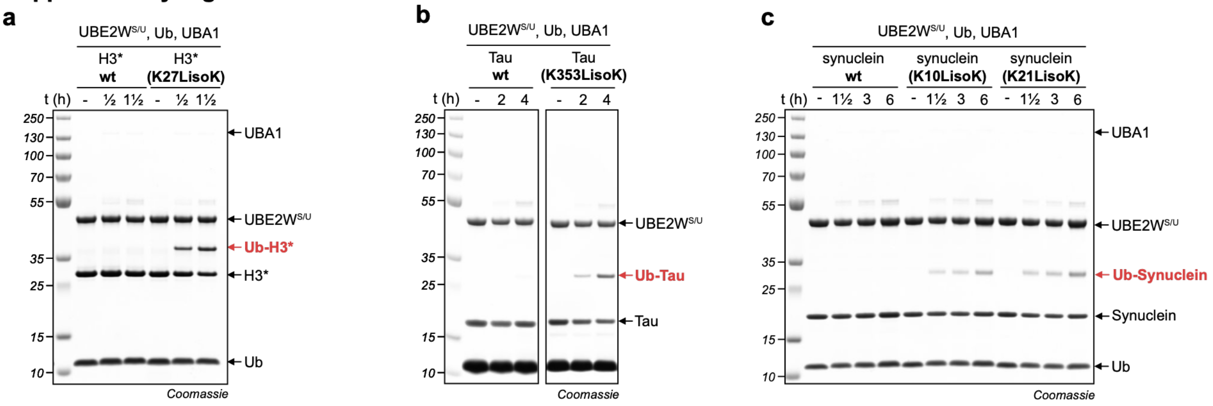

**Supplementary Figure S9. Full SDS-PAGE gels of UBE2WS/U-catalyzed in vitro ubiquitylation across diverse protein folds.** Full SDS-PAGE gels of UBE2W$^{S/U}$ catalyzed *in vitro* ubiquitylation of **a,** wtH3*, and H3*(K27LisoK). The asterisk (*) indicates an N-terminal ubiquitin fusion to enhance protein solubility. Reaction conditions: 40 µM wtH3*/H3*(K27LisoK), 40 µM UBE2W$^{S/U}$, 120 µM Ub, 50 nM UBA1, 50 mM HEPES pH 7.1, 1 mM DTT, 100 mM NaCl, 10 mM ATP, 10 mM $MgCl_2$, 37°C. **b,** wt-tau(244-372), tau(244-372, K353LisoK). Reaction conditions: 40 µM tau-constructs, 40 µM UBE2W$^{S/U}$, 400 µM Ub, 50 nM UBA1, 50 mM HEPES pH 7.1, 1 mM DTT, 100 mM NaCl, 10 mM ATP, 10 mM $MgCl_2$, 37°C. **c,** wt-α-synuclein, α-synuclein(K10LisoK), and α-synuclein(K21LisoK). Reaction conditions: 40 µM α-synuclein-constructs, 40 µM UBE2W$^{S/U}$, 120 µM Ub, 50 nM UBA1, 50 mM HEPES pH 7.1, 1 mM DTT, 100 mM NaCl, 10 mM ATP, 10 mM $MgCl_2$, 37°C.

**Supplementary Figure 10**

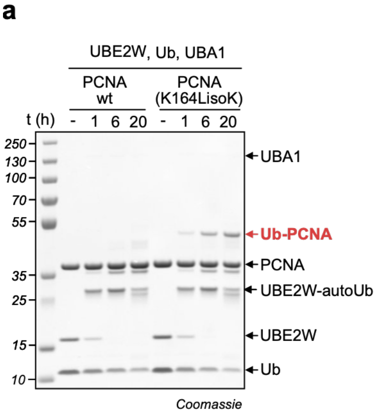

**Supplementary Figure S10. In vitro modification of PCNA by UbyW. a,** SDS-PAGE analysis of *in vitro* modification of wtPCNA and PCNA(K164LisoK) by UBE2W. Reaction conditions: 20 µM PCNA(K164LisoK), 20 µM UBE2W, 60 µM Ub, 100 nM UBA1, 50 mM HEPES pH 7.1, 1 mM DTT, 100 mM NaCl, 10 mM ATP, 10 mM $MgCl_2$, 37°C.

### Supplementary Figure 11

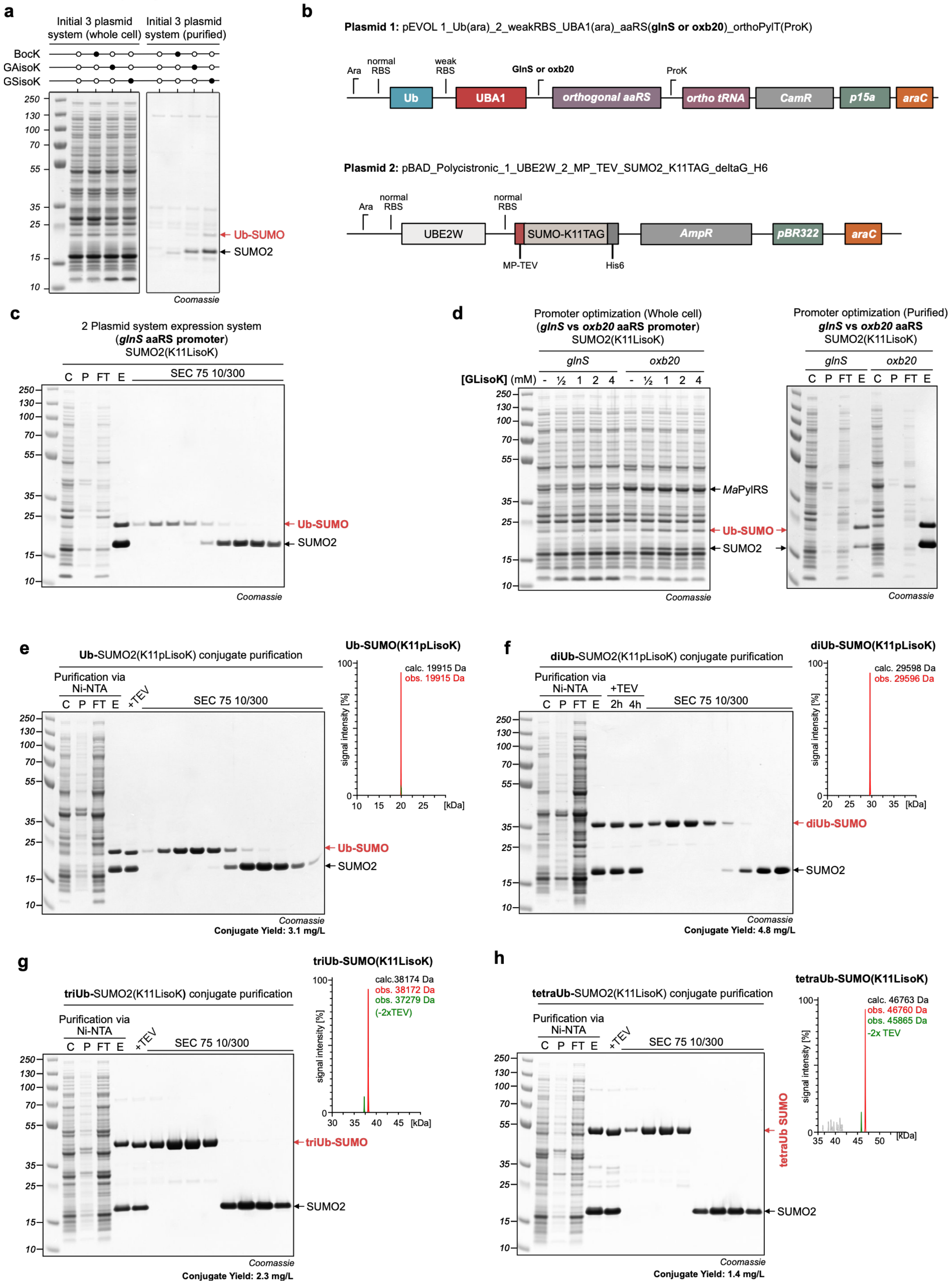

**Supplementary Figure S11. Optimization of the E. coli ubiquitylation cascade and generation of linear polyUb chains. b**, Maps of dual-plasmid system for expressing ubiquitylation and GCE machinery. **c**, SDS-PAGE analysis of Ub-SUMO2(K11LisoK) purification by Ni-NTA and size exclusion chromatography

(Superdex 75 10/300) generated using the Ub cascade in *E. coli*. **d**, Comparison of *glnS* and *oxb20* promoter driving expression of the aminoacyl-tRNA synthetase (aaRS). Whole-cell lysate SDS-PAGE analysis of reaction progression (left) and Ni-NTA purification (right). **e-h,** SDS-PAGE analysis of modified SUMO2 purification by Ni-NTA, subsequent TEV protease cleavage and size exclusion chromatography (Superdex 75 10/300) (left). LC-MS analysis of purified conjugate: Representative deconvoluted electrospray ionization mass spectra showing observed and calculated mass (right). Non-deconvoluted m/z spectra are shown in Supplementary Figure SI 13. Modification with Ub (**d**), diUb (**e**) triUb, (**f**) tetraUb, (**g**). C: culture, P: pellet, FT: Flow through Ni-NTA, E: Elution Ni-NTA. Reaction conditions: *E. coli* K12 cultured in auto-induction media (AI, 14 aa, 10 mM NAM, 37°C, over-night, 2 mM GLisoK).

**Supplementary Figure 12**

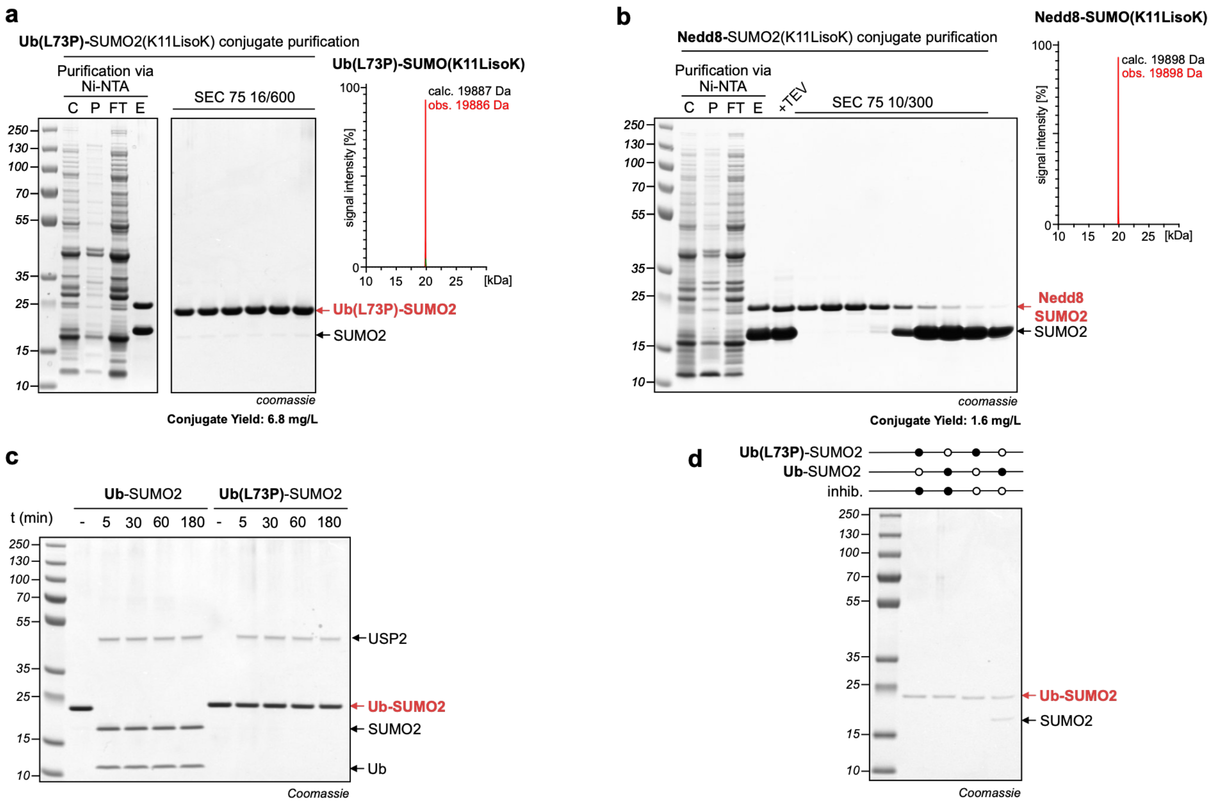

**Supplementary Figure S12. Conjugation of Ub(L73P) and Nedd8. a,** SDS-PAGE analysis of Ub-SUMO2(K11LisoK) purification by Ni-NTA, subsequent TEV protease cleavage and size exclusion chromatography (Superdex 75 10/300) (left). LC-MS analysis of purified conjugate: Representative deconvoluted electrospray ionization mass spectra showing observed and calculated mass (right). Non-deconvoluted m/z spectra are shown in Supplementary Figure SI 13. Modification with Ub(L73P) (**a**), and Nedd8 (**b**). C: culture, P: insoluble after sonication, FT: Flow through Ni-NTA, E: Elution Ni-NTA. Reaction conditions: *E. coli* K12 cultured in auto-induction media (AI, 14 aa, 10 mM NAM, 37°C, over-night, 2 mM GLK). **c,** SDS-PAGE analysis of incubation of Ub-SUMO2(K11LisoK) and Ub(L73P)-SUMO2(K11LisoK) conjugate with DUB USP2 *in vitro.* **d,** SDS-PAGE analysis of Ni-NTA purification of Ub-SUMO2(K11LisoK) and Ub(L73P)-SUMO2(K11LisoK) conjugate after incubation in HEK293T cell lysate in presence or absence of DUB inactivating inhibitors.

### Supplementary Figure 13

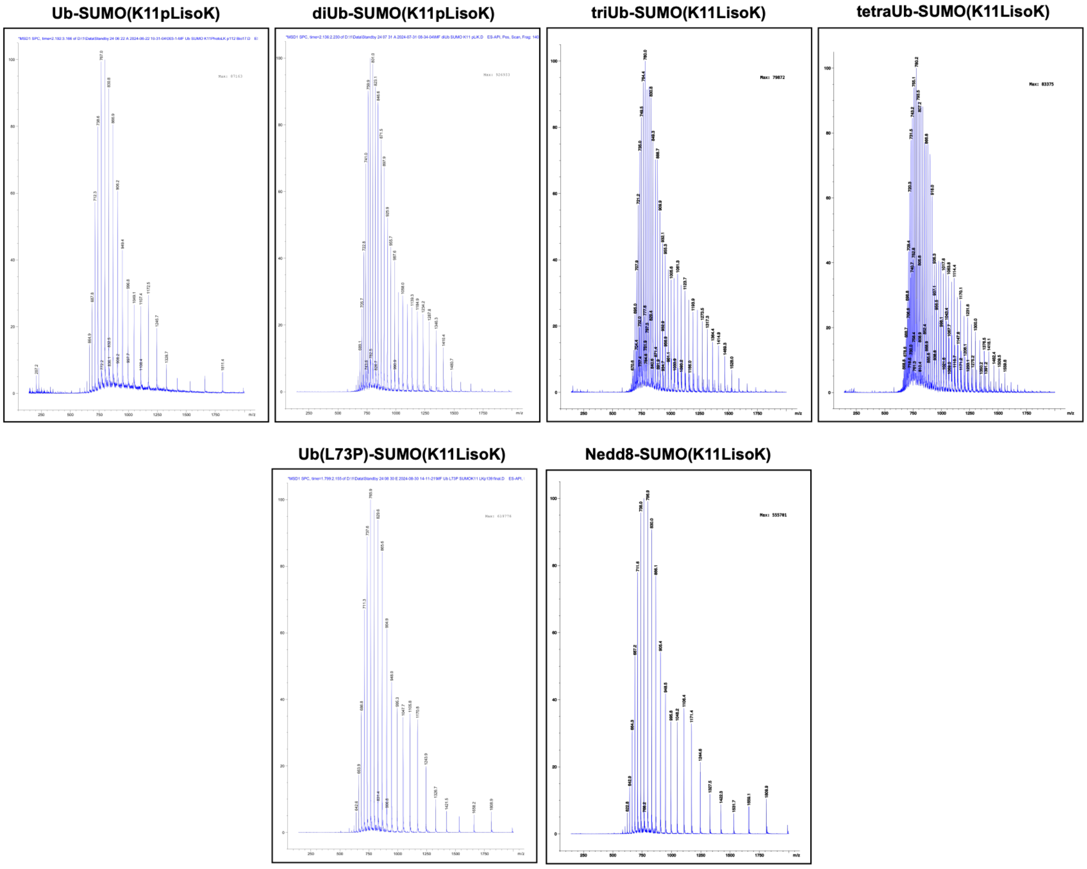

**Supplementary Figure S13. Raw ESI-MS spectra of SUMO2(K11pLisoK) modified with polyUb chains, Ub(L73P), and Nedd8.** ESI-MS m/z spectrum of SUMO2 modified with Ub, diUb, triUb, tetraUb, Ub(L73P) and Nedd8 in native charge state distribution (non-deconvoluted).

### Supplementary Figure 14

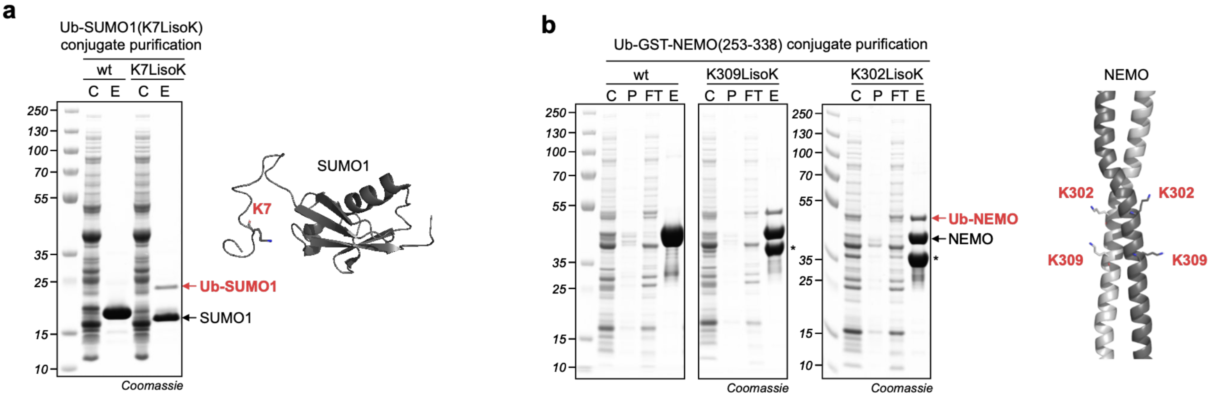

**Supplementary Figure S14. Modification of SUMO1 and NEMO using the cellular ubiquitylation system. a,** Left: Structure of SUMO1 with K7 highlighted in red (Protein Data Bank, 2kqs)[4]. Right: SDS-PAGE analysis of UBE2W-mediated modification of SUMO1(K7LisoK) in *E. coli.* **b,** Left: Structure of NEMO(253-338) homo-dimer with K302 and K309 highlighted in red (Protein Data Bank, 7tv4)[5]. Right: SDS-PAGE analysis of UBE2W mediated modification of GST-NEMO(253-338, K309LisoK) and GST-NEMO(253-338, K302LisoK) in *E. coli.* Reaction conditions: *E. coli* K12 cultured in auto-induction media (AI, 14 aa, 10 mM NAM, 37°C, over-night, 2 mM GLisoK). The asterisk (*) indicates the GST-NEMO truncation due to competition between RF1 and suppressor PylT during GCE.

### Supplementary Figure 15

**a**

| *Origin* | *Sequence ID* | *Foldseek score* |
|---|---|---|
| *Homo sapiens* | 100 | 690 |
| *Danio rerio* | 93.3 | 662 |
| *Dinothrombium tinctorium* | 79.2 | - |
| *Drosophila melanogaster* | 71.6 | 565 |
| *Caenorhabditis elegans* | 59.7 | 523 |
| *Cyclocybe aegerita* | 48.2 | 370 |
| *Zea Mays* | 49.6 | 439 |
| *Plasmodium falciparum* | 39 | 382 |

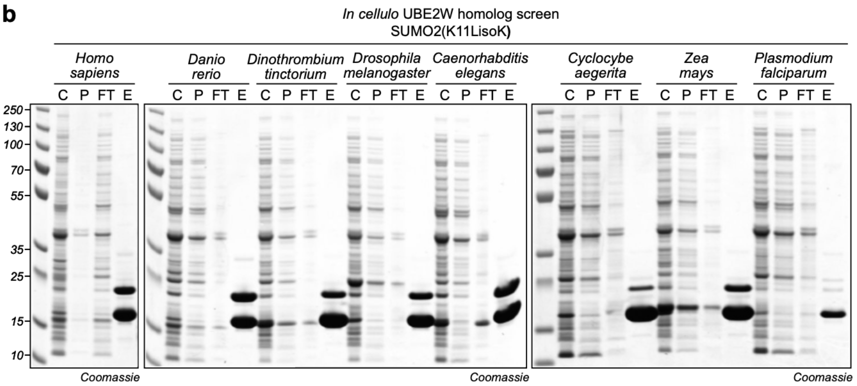

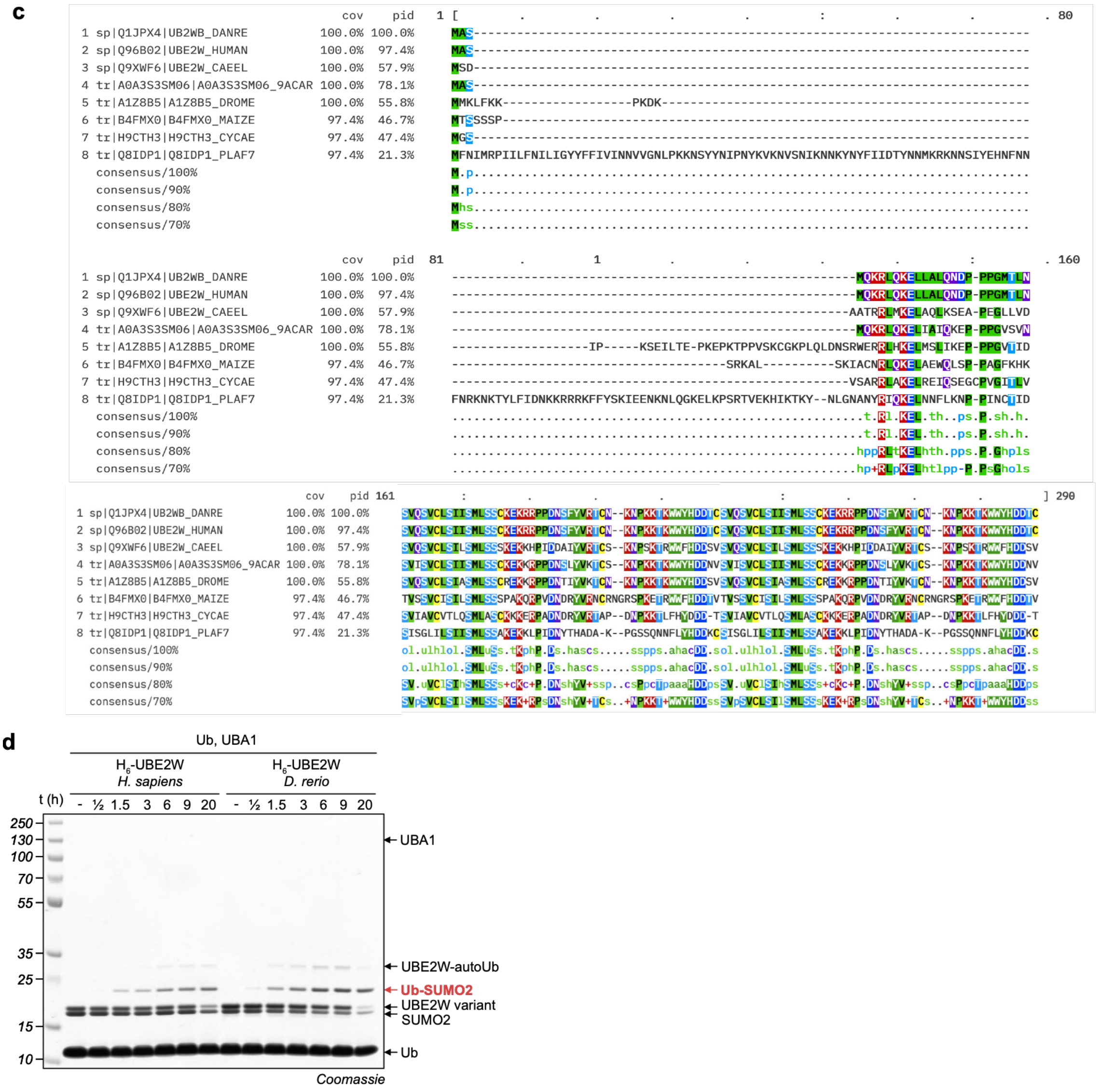

**Supplementary Figure S15.** ***In cellulo*** **UBE2W homolog screen and conjugate purification**. **a,** Comparison of sequence identity (sequence ID) and Foldseq score of selected UBE2W homologs.[6] **b**, SDS-PAGE analysis of Ni-NTA purified SUMO2(K11LisoK) modified with Ub by the different UBE2W homologs in the *E. coli* cascade. Reaction conditions: *E. coli* K12 cultured in auto-induction media (AI, 14 aa, 10 mM NAM, 37°C, over-night, 2 mM GLisoK). **c,** MSA of UBE2W homologs, generated using the EMBL-EBI Job Dispatcher sequence analysis tools framework in

2024. **d,** SDS-PAGE analysis of reaction progression *in vitro* of SUMO2(K11LisoK) modification with Ub, catalyzed by *Hs*UBE2W and *Dr*UBE2W. Reaction conditions: 40 SUMO(K11LisoK), 40 μM *Hs*UBE2W$^{S/U}$/ *Dr*UBE2W$^{S/U}$, 400 μM Ub, 50 nM UBA1, 50 mM HEPES pH 7.1, 1 mM DTT, 100 mM NaCl, 10 mM ATP, 10 mM $MgCl_2$, 37°C.

**Supplementary Figure 16**

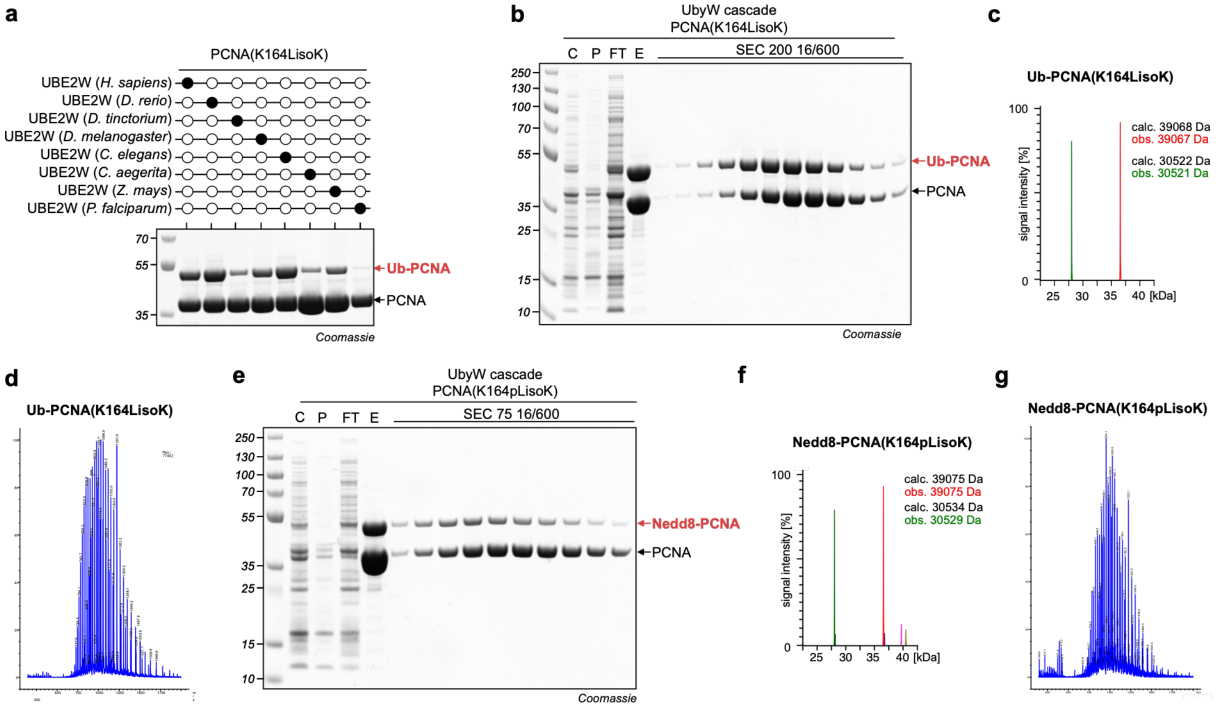

**Supplementary Figure S16. Modification of PCNA via the UbyW cascade. a,** SDS-PAGE analysis of Ni-NTA purified modified and unmodified PCNA(K164LisoK) generated by a selection of different UBE2W homologs in *E. coli*. **b,** SDS-PAGE analysis of Ni-NTA purification and size exclusion chromatography (Superdex 200 16/600) of Ub-PCNA(K164LisoK) generated via the UbyW cascade. **c,** Deconvoluted ESI-MS of Ub-PCNA(K164LisoK) showing observed and calculated masses. **d,** ESI-MS m/z spectrum of Ub-PCNA(K164LisoK). **e,** SDS-PAGE analysis of Ni-NTA purification and size exclusion chromatography (Superdex 75 16/600) of Nedd8-PCNA(K164LisoK) generated via the UbyW cascade. **f,** Deconvoluted ESI-MS of Nedd8-PCNA(K164LisoK) showing observed and calculated masses. **g,** ESI-MS m/z spectrum of Nedd8-PCNA(K164pLisoK). Reaction conditions: *E. coli* K12 cultured in auto-induction media (AI, 14 aa, 10 mM NAM, 37°C, overnight, 2 mM GLisoK or GpLisoK).

**Supplementary Figure 17**

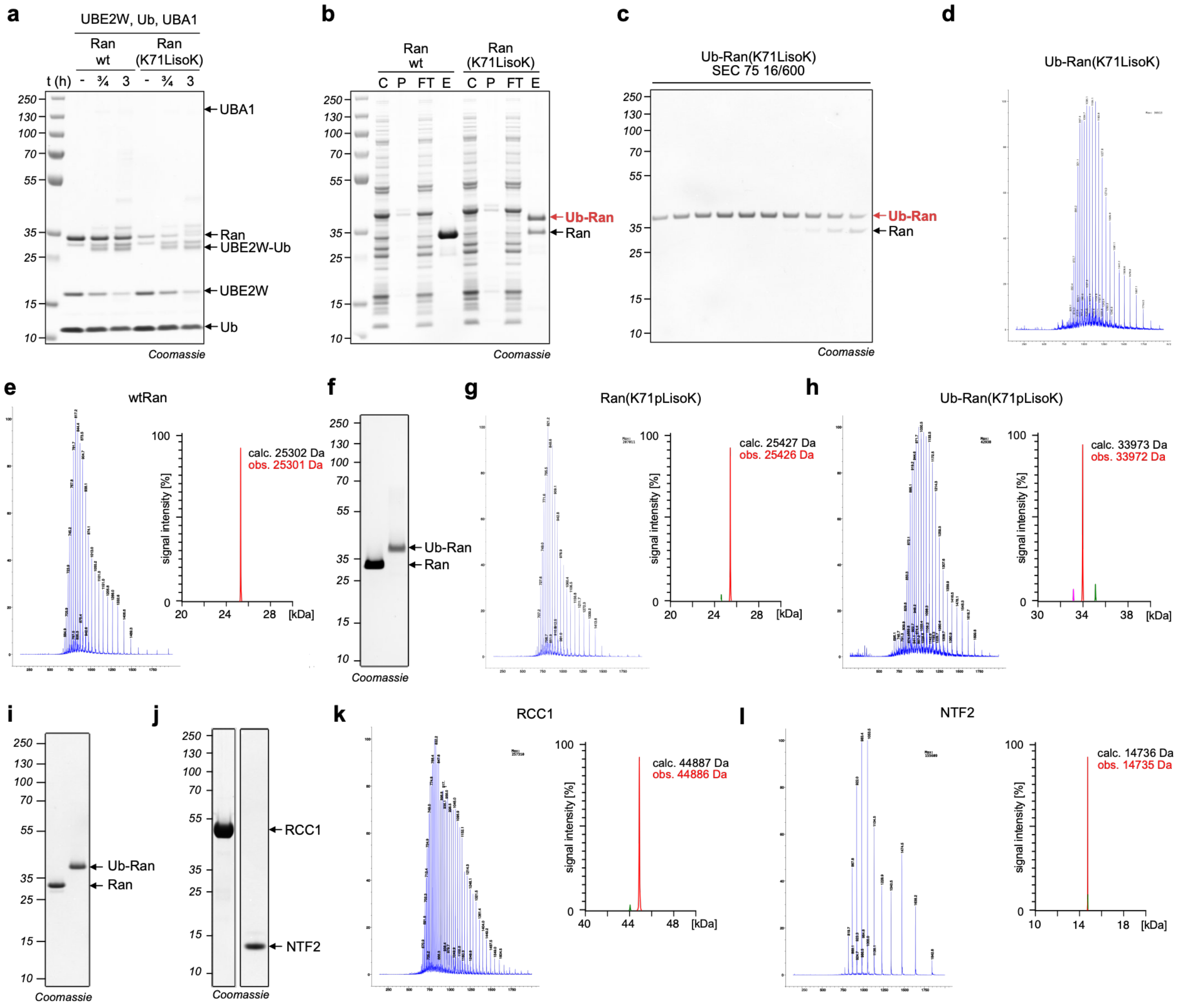

**Supplementary Figure S17. Modification of Ran via the UbyW cascade. a,** SDS-PAGE analysis of *in vitro* modification of wtRan and Ran(K71LisoK) by UBE2W. Reaction conditions: 20 µM Ran construct, 20 µM UBE2W, 60 µM Ub, 100 nM UBA1, 50 mM HEPES pH 7.1, 1 mM DTT, 100 mM NaCl, 10 mM ATP, 10 mM MgCl, 37°C. **b,** SDS-PAGE analysis of Ni-NTA purification of wtRan and modified Ran(K71LisoK) using UBE2W in the reconstituted Ub cascade. Reaction conditions: *E. coli* K12 cultured in auto-induction media (AI, 14 aa, 10 mM NAM, 37°C, over-night, 2 mM GLK). **b,** SDS-PAGE analysis of the size exclusion chromatography (Superdex 75 16/600) of Ub-Ran(K71LisoK). **d,** ESI-MS m/z spectrum of Ub-Ran(K71LisoK) in native charge state distribution (non-deconvoluted), a representative deconvoluted spectrum is shown in Figure 5e. **e,** Left: ESI-MS m/z spectrum of wtRan, Right: Deconvoluted ESI-MS of wtRan showing observed and calculated masses. **f,** SDS-PAGE analysis of purified Ran(K71pLisoK) and Ub-Ran(K71pLisoK) used for LFQ-AP-MS experiments. **g,** Left, ESI-MS m/z spectrum of Ran(K71pLisoK), Right: Deconvoluted ESI-MS of Ran(K71pLisoK) showing observed and calculated masses. **h,** Left: ESI-MS m/z spectrum of Ub-Ran(K71pLisoK), Right: Deconvoluted ESI-MS of Ub-Ran(K71pLisoK) showing observed and calculated masses. **i,** SDS-PAGE analysis of purified and Mant-GDP charged wtRan and Ub-Ran(K71LisoK) used for the nucleotide exchange assays

(Fig. 5i, j, Supplementary Fig. S19e, f) **j,** SDS-PAGE analysis of purified RCC1 and NTF2 used for nucleotide exchange assays (Fig. 5i and 5j, Supplementary Fig. S19e, f) or *in vitro* co-affintiy purification assays (Supplementary Fig. S19c) respectively **k,** Left: ESI-MS m/z spectrum of RCC1, Right: Deconvoluted ESI-MS of RCC1 showing observed and calculated masses. **l,** Left: ESI-MS m/z spectrum of NTF2, Right: Deconvoluted ESI-MS of NTF2 showing observed and calculated masses.

**Supplementary Figure 18**

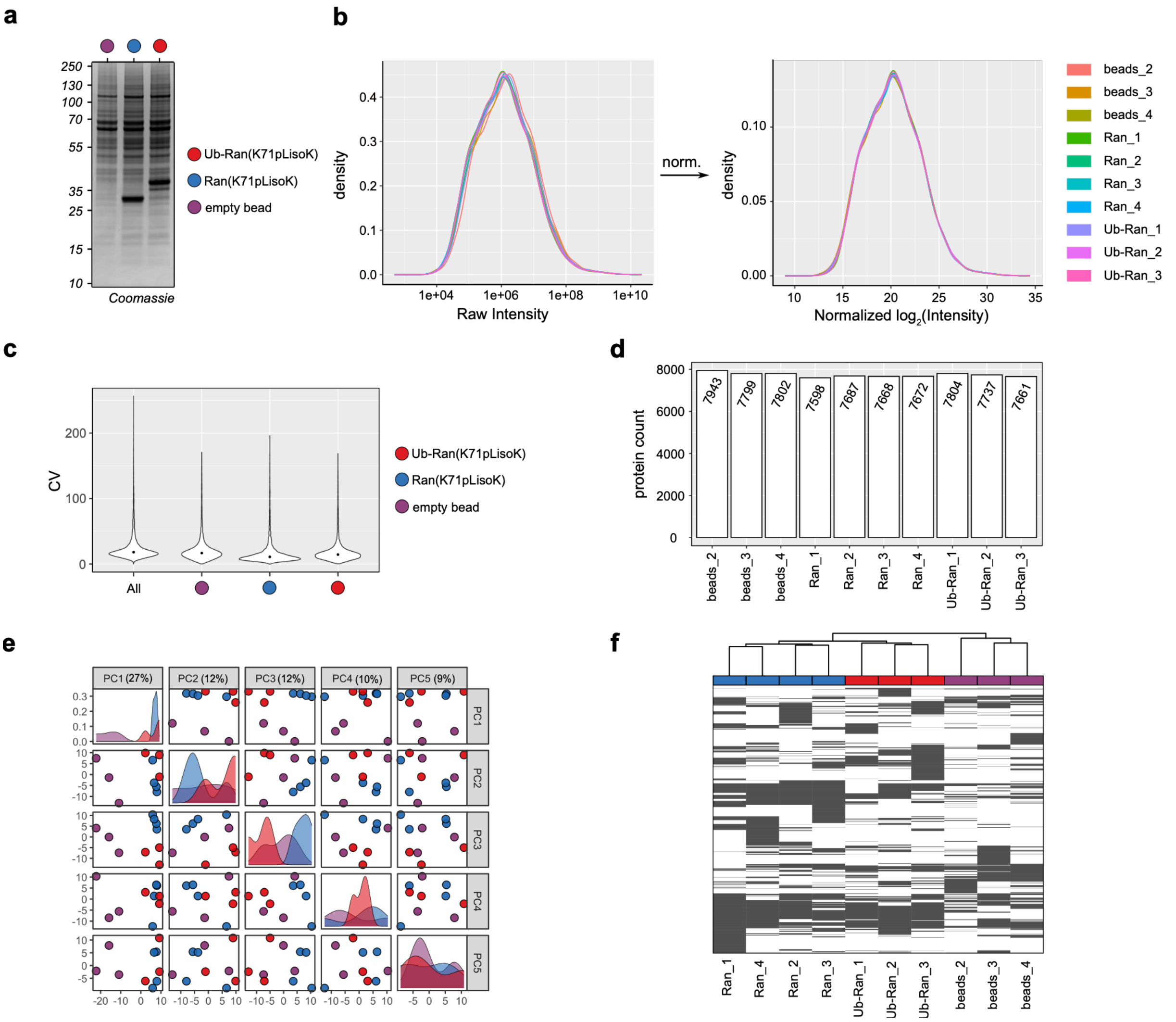

**Supplementary Figure S18. Quality control of crosslinking LFQ-AP-MS proteomics for Ran(K71) probes. a,** SDS-PAGE analysis of representative input samples **b,** Kernel density estimates of protein log2-intensities before (left, raw) and after (right) robust scale (robscale) normalization using prolfqua. Consistent distribution profiles across runs confirms successful normalization.[7] **c,** Coefficient of variation (CV) distributions for all quantified proteins (left) and separated by experimental condition (right). Median CVs: 18.0% overall, 16.7% for beads, 10.9% for Ran(K71pLisoK), and 14.4% for Ub-Ran(K71pLisoK). **d,** Total number of identified proteins per LC-MS/MS run. **e,** Principal component analysis (PCA) of normalized protein intensities. **f,** Missingness map illustrating the distribution of missing values across all samples and identified proteins. **g,** In all applicable panels, violet denotes the bead control (no probe), blue denotes Ran(K71pLisoK), and red denotes Ub-Ran(K71pLisoK).

**Supplementary Figure 19**

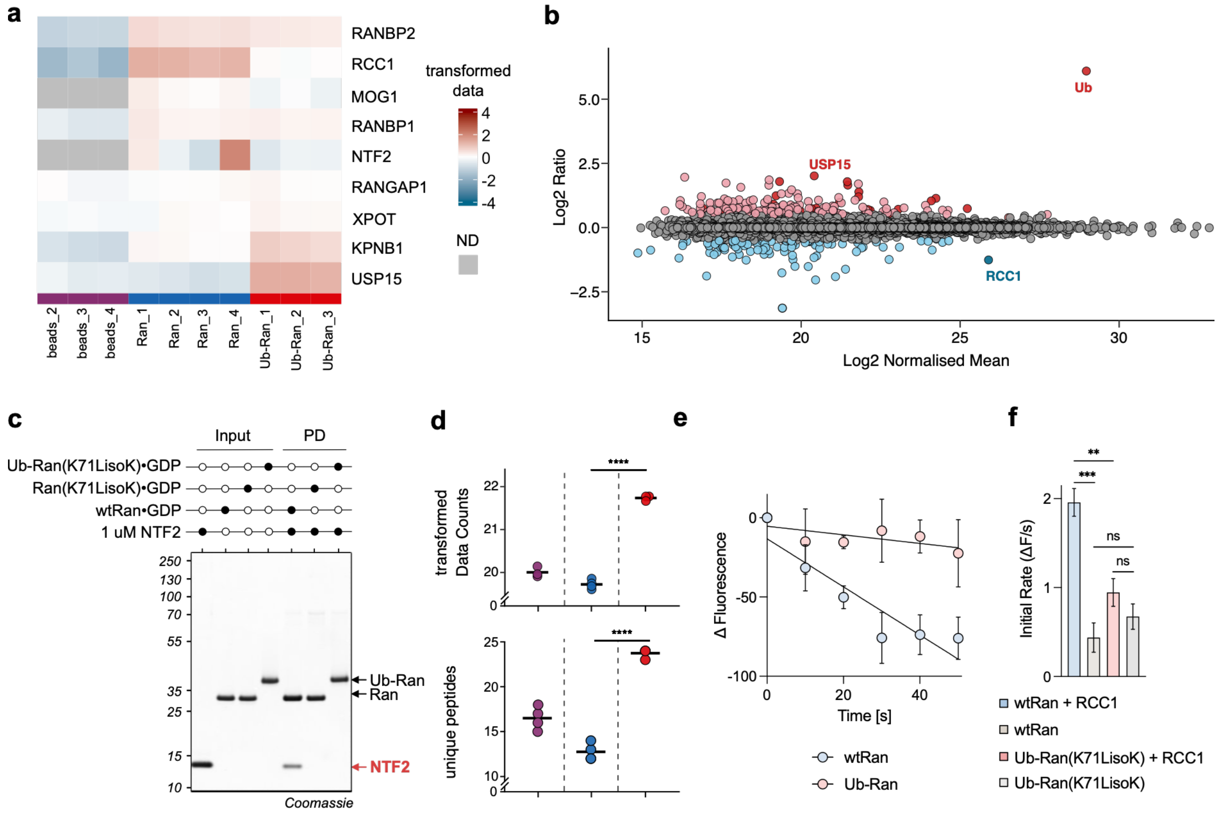

**Supplementary Figure S19. AP-MS target validation and biochemical characterization of Ran(K71) conjugates. a,** Heatmap of normalized protein intensities for known Ran interactors and USP15 across AP-MS conditions. Grey cells indicate proteins not detected (ND) in the respective condition. Color intensity reflects normalized protein abundance, with warmer colors indicating higher abundance. **b,** MA plot comparing protein enrichment between the Ran(K71pLisoK) and Ub-Ran(K71pLisoK) samples. **c,** *In vitro* co-affinity purification of NTF2 (1 µM) using wild-type Ran (wtRan), Ran(K71LisoK), and Ub-Ran(K71LisoK). Bait beads were charged with 4 µg of the respective Ran species, which were selectively loaded with GDP prior to incubation. *Left:* input samples; *Right:* Pulldown (PD) fractions. Only wtRan successfully captures NTF2. **d,** Validation of the USP15 AP-MS hit. *Top:* Transformed intensity counts across conditions. Statistical significance was determined using a linear model followed by an empirical Bayes moderated t-test. *Bottom:* Number of unique USP15 peptides detected per run. Statistical significance between Ran(K71pLisoK) and Ub-Ran(K71pLisoK) was determined using an ordinary one-way ANOVA followed by Tukey's multiple comparisons test (**** $P < 0.0001$). Both metrics demonstrate significant enrichment in the ubiquitylated bait samples. violet denotes the bead control (no probe), blue denotes Ran(K71pLisoK), and red denotes Ub-Ran(K71pLisoK). **e,** Initial linear phase (0–50 s) of the Ran nucleotide exchange reaction monitored via mant-GDP fluorescence. Data is background-corrected using matched samples lacking RCC1. Points represent mean ΔFluorescence ± SD (n = 5 independent replicates). Solid lines represent simple linear regression fits used to calculate the initial exchange velocity ($v_0$). The corresponding bar graph comparing these rates is shown in Figure 5. **f,** Absolute initial nucleotide exchange rates ($v_0$) for wtRan and Ub-Ran(K71LisoK) in the presence and absence of the RCC1 exchange factor. Bar heights correspond to the absolute slope. Error bars represent the standard error of the regression slope (SEM). Statistical significance for pre-planned comparisons was determined using unpaired Welch's t-tests to account for unequal variances between conditions (*** $P < 0.001$; ** $P < 0.01$; ns, not significant).

**Supplementary Figure 20**

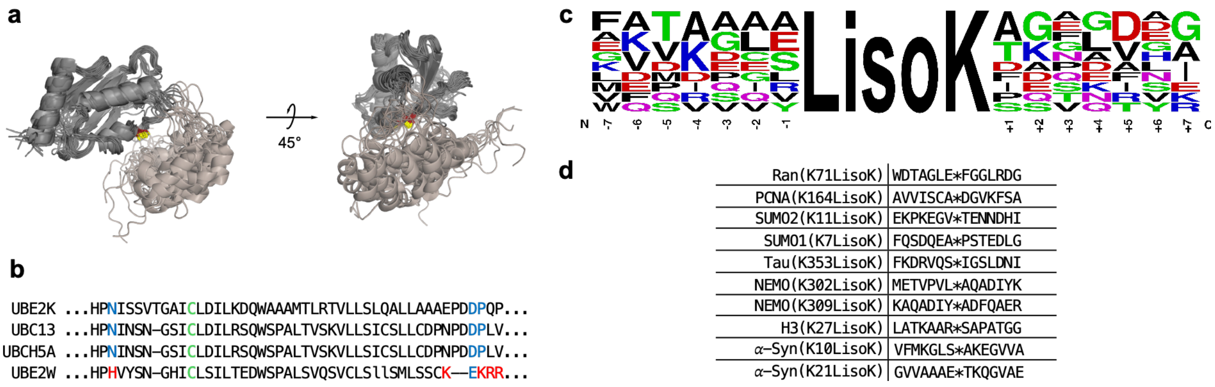

**Supplementary Figure S20. Structural features and target sequence independence of UBE2W. a,** Alignment of the 20 top scoring models of UBE2W shown in two orientations. The C-terminal tail of Ube2W is much more conformationally flexible in comparison to the catalytic core. **b,** Sequence alignment of the active site of selected E2s. The active site Cys is colored green, the highly conserved Asp of the HPN motif as well as gatekeeper acidic patch found in most E2s is colored blue. Red indicates differences in UBE2W. **c,** Sequence logo of the amino acid context of target proteins endowed with LisoK and modified by UBE2W in UbyW. Generated using WebLogo.[8] **d,** Table of individual surrounding sequence to LisoK (✻) in the target proteins, used to generate the sequence logo in Supplementary Fig. 18c.

### Materials and Methods

#### 1. General Methods: Plasmids and Reagents

Codon optimized genes encoding all UBE2W variants and homologues, SUMO1/2, linear di/tri/tetra Ubiquitin, Ub, PCNA, α-synuclein, tau (244-372), GST-NEMO (253-338), Ub-H3, UBA1, RCC1, NTF2, UBE2N, UBE2V1 and Ran were purchased as DNA Strings (Twist Bioscience) and cloned into the pBAD, pEVOL, or pET28 vectors via restriction cloning (see Supplementary Table S1). Amino acid sequences of all proteins are listed in Supplementary Table S2.

All solvents and chemical reagents were purchased from Sigma Aldrich, Senn, cslabs, Carbolution, Acros Organics, or Fisher Scientific. Fmoc building blocks of Propaglyglycine (PrgG) and Chloroalanine (ClA) were purchased from Iris Biotech GmbH. All Reagents were used without further purification unless stated otherwise.

All GXisoK non-canonical amino acids such as: GLisoK, GAisoK, GSisoK, GTisoK, GVisoK, GaisoK, GsisoK, GLisoK, GCisoK, GPisoK, GPrgGisoK, GClAisoK were prepared as previously described.[2]

H6-UBE1 was purchased from Boston Biochem (Cat. No. E-304-050). USP15 was purchased from Sino Biological (Cat. No. U515-380G). Mant-GDP was purchased from Sigma Aldrich (69244).

The sequence coding for UBA1 was a gift from Titia Sixma, Addgene #63571, UniProt P22314-2).[9]

Ni-NTA agarose was purchased from Cytivia (Cat. No. 17-5318-02). Magentic Ni-NTA beads were purchased from Cube Biotech (Cat. No. 31201).

Bolt 4-12 % Bis-Tris gradient gels (Invitrogen) gels were run (at 165 V for 45 min) using a Bolt™ Mini Gel Tank system (Invitrogen) using a MES running buffer (50 mM, pH 6, 50 mM Tris, 0.1% (w/v) SDS, 1 mM EDTA. Gels were stained with Quick Coomassie Stain (Generon). PageRuler Prestained Plus Protein Ladder 10-250 kDa (ThermoFisher) was used as the protein marker.

Protein and DNA concentrations were measured on a NanoPhotometer® NP60 (Implen).

### 1.1 Plasmids

#### Supplementary Table S1. Plasmids employed in this study and their respective description.

| Plasmid | Description |
|---|---|
| pBAD_RSF1031K_UBE2W_(*H.sapiens*) (OP1154) | Tagless human UBE2W under an arabinose promotor (kanamycin resistance). |
| pBAD_RSF1031K_MP_His_TEV_UBE2W _(*H.sapiens*) (OP1299) | Human UBE2W with an N-terminal proline followed by a H6-tag and a TEV site under an arabinose promotor (kanamycin resistance). |
| pBAD_MP_H8_TEV_SUMO2_Ub_UBE2W _(*H.sapiens*) (OP1610) | Human UBE2W N-terminally fused to SUMO-Ub with an N-terminal proline followed by a H8-tag and a TEV site under an arabinose promotor (ampicillin resistance). |
| pBAD_MP_Strep_TEV_SUMO2v_Ub_UB E2W_(*H.sapiens*) (OP1614) | Human UBE2W N-terminally fused to SUMO-Ub with an N-terminal proline followed by a StrepII-tag and a TEV site under an arabinose promotor (ampicillin resistance). |
| pBAD_RSF1031K_MP_His_TEV_UBE2W _(D. rerio) (OP1620) | *D. rerio* UBE2W with an N-terminal proline followed by a H6-tag and a TEV site under an arabinose promotor (kanamycin resistance). |
| pBAD_MP_TEV_SUMO2_wt_deltaG_H6 (OP1181) | SUMO2-wt-H6 with an N-terminal proline followed by a TEV site and a G93 deletion with a C-terminal H6-tag under an arabinose promotor (ampicillin resistance). |
| pBAD_MP_TEV_SUMO2_K11TAG_ deltaG_H6 (OP1182) | SUMO2-K11TAG-H6 with an N-terminal proline followed by a TEV site and a G93 deletion with a C-terminal H6-tag under an arabinose promotor (ampicillin resistance). |
| pBAD_MP_TEV_PCNA_wt_His6 (OP1198) | Human PCNA-wt-H6 with an N-terminal proline followed by a TEV site under an arabinose promotor with a C-terminal H6-tag (ampicillin resistance). |
| pBAD_MP_TEV_PCNA_K164TAG_His6 (OP1197) | Human PCNA-K164TAG-H6 with an N-terminal proline followed by a TEV site under an arabinose promotor with a C-terminal H6-tag (ampicillin resistance). |
| pBAD_MP_Ub_TEV_H3_wt_H6 (OP1716) | Human Histone H3-wt-H6 N-terminally fused to Ub with an N-terminal proline and a TEV site between Ub and H3 under an arabinose promotor with a C-terminal H6-tag (ampicillin resistance). |
| pBAD_MP_Ub_H3_K27TAG_H6 (OP1715) | Human Histone H3-K27TAG-H6 N-terminally fused to Ub with an N-terminal proline and a TEV site between Ub and H3 under an arabinose promotor with a C-terminal H6-tag (ampicillin resistance). |
| pBAD_MP_H6_TEV_tau(244-372)_wt (OP1465) | Human tau(244-372)-wt with an N-terminal proline and a H6-tag site followed by a TEV site under an arabinose (ampicillin resistance). |
| pBAD_MP_H6_TEV_tau(244-372)_K353TAG (OP1470) | Human tau(244-372)-K353TAG with an N-terminal proline and a H6-tag site followed by a TEV site under an arabinose (ampicillin resistance). |
| pBAD_MP_H6_TEV_a-synuclein_wt (OP1460) | Human alpha-synuclein-wt with an N-terminal proline followed by a H6-tag and a TEV site under an arabinose promotor (ampicillin resistance). |
| pBAD_MP_H6_TEV_a-synuclein_K10TAG (OP1461) | Human alpha-synuclein-K10TAG with an N-terminal proline followed by a H6-tag and a TEV site under an arabinose promotor (ampicillin resistance). |
| pBAD_MP_H6_TEV_a-synuclein_K21TAG (OP1463) | Human alpha-synuclein-K21TAG with an N-terminal proline followed by a H6-tag and a TEV site under an arabinose promotor (ampicillin resistance). |
| pBAD_MP_H6_TEV_Ran_wt (OP1274) | Human Ran-wt with an N-terminal proline followed by a H6-tag and a TEV site under an arabinose promotor (ampicillin resistance). |
| pBAD_MP_H6_TEV_Ran_K71TAG (OP1275) | Human Ran-K71TAG with an N-terminal proline followed by a H6-tag and a TEV site under an arabinose promotor (ampicillin resistance). |

| | |
|---|---|
| pET17b_Ubiquitin_wt (UbC1) | Human Ubiquitin wt under a T7 promotor (ampicillin resistance). |
| pBAD_MP_H6_TEV_diUb_(G76V) (OP1282) | Linear diubiquitin with an N-terminal proline followed by a H6-tag and TEV site and a G76V mutation at the distal ubiquitin under an arabinose promotor with a (ampicillin resistance). |
| pBAD_MP_H6_TEV_triUb_(2xG76V) (OP1281) | Linear triubiquitin with an N-terminal proline followed by a H6-tag and TEV site and a G76V mutation at the two distal ubiquitins under an arabinose promotor (ampicillin resistance). |
| pBAD_MP_H6_TEV_tetraUb_(3xG76V) (OP1283) | Linear tetraubiquitin with an N-terminal proline followed by a H6-tag and TEV site and a G76V mutation at the three distal ubiquitins under an arabinose promotor (ampicillin resistance). |
| pGEX6P1-UBE2N (OP11) | Human GST-UBE2N under a T7 promotor (ampicillin resistance). |
| pGEX6P1-UBE2V1 (OP12) | Human GST-UBE2V1 under a T7 promotor (ampicillin resistance). |
| pEVOL_Mb_wt_aaRS _PylT | *M. barkeri* wt PylRS under a constitutive glnS promotor and an arabinose inducible promoter with a *M. alvus* PylT copy under a constitutive promotor (chloramphenicol resistance). |
| pEVOL_Ma_aaRS_H227I_Y288P_PylT | *M. alvus* H227I Y288P PylRS under a constitutive glnS promotor and an arabinose inducible promoter with a *M. alvus* PylT copy under a constitutive promotor (chloramphenicol resistance). |
| pEVOL_1_normalRBS_Ub_wt_2_weakRBS_UBA1(polycistronic_ara)_Ma_wt_IP_(glnS) (OP1289) | Human Ub-wt under an arabinose inducible promotor with a normal RBS followed by human UBA1 with a weak RBS followed by *M. alvus* H227I Y288P PylRS under a constitutive glnS promotor and a *M. alvus* PylT copy under a constitutive promotor (chloramphenicol resistance). |
| pEVOL_1_normalRBS_Ub_wt_2_weakRBS_UBA1(polycistronic_ara)_Ma_wt_IP (oxb20)_Ma_tRNA (OP1325) | Human Ub-wt under an arabinose inducible promotor with a normal RBS followed by human UBA1 with a weak RBS followed by *M. alvus* H227I Y288P PylRS under a strong constitutive oxb20 promotor and a *M. alvus* PylT copy under a constitutive promotor (chloramphenicol resistance). |
| pEVOL_1_normalRBS_Ub(L73P)_wt_2_weakRBS_UBA1(polycistronic_ara)_Ma_wt IP_(oxb20)_Ma_tRNA (OP1395) | Human Ub(L73P) under an arabinose inducible promotor with a normal RBS followed by human UBA1 with a weak RBS followed by *M. alvus* H227I Y288P PylRS under a strong constitutive oxb20 promotor and a *M. alvus* PylT copy under a constitutive promotor (chloramphenicol resistance). |
| pEVOL_1_normalRBS_MP_TEV_diUb_2_weakRBS_UBA1(polycistronic_ara)_Ma_wt_IP_(oxb20)_Ma_tRNA (OP1363) | Human Ub-wt with a N-terminal proline followed by TEV site and a G76V mutation at the distal Ub under an arabinose inducible promotor with a normal RBS followed by human UBA1 with a weak RBS followed by *M. alvus* H227I Y288P PylRS under a strong constitutive oxb20 promotor and a *M. alvus* PylT copy under a constitutive promotor (chloramphenicol resistance). |
| pEVOL_1_normalRBS_Nedd8_2_weakRBS_UBA1(polycistronic_ara)_Ma_wt_IP_(oxb20)_Ma_tRNA (OP1567) | Human Nedd8-wt under an arabinose inducible promotor with a normal RBS followed by human UBA1 with a weak RBS followed by *M. alvus* H227I Y288P PylRS under a strong constitutive oxb20 promotor and a *M. alvus* PylT copy under a constitutive promotor (chloramphenicol resistance). |
| pBAD_Polycistronic_1_UBE2W_(*H.sapiens*)_2_MP_TEV_SUMO2_K11TAG_deltaG_H6 (OP1287) | *H. sapiens* UBE2W followed by SUMO2-K11TAG with an N-terminal proline followed by a TEV site and a G93 deletion with a C-terminal H6-tag under an arabinose promotor (ampicillin resistance). |
| pBAD_Polycistronic_1_UBE2W_(*D.rerio*)_2_MP_TEV_SUMO2_K11TAG_deltaG_H6 (OP1478) | *D. rerio* UBE2W followed by SUMO2-K11TAG with an N-terminal proline followed by a TEV site and a G93 deletion with a C-terminal H6-tag under an arabinose promotor (ampicillin resistance). |
| pBAD_Polycistronic_1_UBE2W_(*D.tinctorium*)_2_MP_TEV_SUMO2_K11TAG_deltaG_H6 (OP1479) | *D. tinctorium* UBE2W followed by SUMO2-K11TAG with an N-terminal proline followed by a TEV site and a G93 deletion with a C-terminal H6-tag under an arabinose promotor (ampicillin resistance). |
| pBAD_Polycistronic_1_UBE2W_(*D.melan* | *D. melanogaster* UBE2W followed by SUMO2-K11TAG with an N-terminal proline |

| | |
|---|---|
| *ogaster*)_2_MP_TEV_SUMO2_K11TAG_deltaG_H6 (OP1480) | followed by a TEV site and a G93 deletion with a C-terminal H6-tag under an arabinose promotor (ampicillin resistance). |
| pBAD_Polycistronic_1_UBE2W_(*C.elegans*)_2_MP_TEV_SUMO2_K11TAG_deltaG_H6 (OP1481) | *C. elegans* UBE2W followed by SUMO2-K11TAG with an N-terminal proline followed by a TEV site and a G93 deletion with a C-terminal H6-tag under an arabinose promotor (ampicillin resistance). |
| pBAD_Polycistronic_1_UBE2W_(*C.aegerita*)_2_MP_TEV_SUMO2_K11TAG_deltaG_H6 (OP1482) | *C. aegerita* UBE2W followed by SUMO2-K11TAG with an N-terminal proline followed by a TEV site and a G93 deletion with a C-terminal H6-tag under an arabinose promotor (ampicillin resistance). |
| pBAD_Polycistronic_1_UBE2W_(*Z.mays*)_2_MP_TEV_SUMO2_K11TAG_deltaG_H6 (OP1483) | *Z. mays* UBE2W followed by SUMO2-K11TAG with an N-terminal proline followed by a TEV site and a G93 deletion with a C-terminal H6-tag under an arabinose promotor (ampicillin resistance). |
| pBAD_Polycistronic_1_UBE2W_(*P.falciparum*)_2_MP_TEV_SUMO2_K11TAG_deltaG_H6 (OP1483) | *P. falciparum* UBE2W followed by SUMO2-K11TAG with an N-terminal proline followed by a TEV site and a G93 deletion with a C-terminal H6-tag under an arabinose promotor (ampicillin resistance). |
| pBAD_Polycistronic_1_UBE2W_(*H.sapiens*)_2_MP_TEV_PCNA_K164TAG_H6 (OP1301) | *H. sapiens* UBE2W followed by PCNA-K164TAG with an N-terminal proline followed by a TEV site with a C-terminal H6-tag under an arabinose promotor (ampicillin resistance). |
| pBAD_Polycistronic_1_UBE2W_(*D.rerio*)_2_MP_TEV_PCNA_K164TAG_H6 (OP1582) | *D. rerio* UBE2W followed by PCNA-K164TAG with an N-terminal proline followed by a TEV site with a C-terminal H6-tag under an arabinose promotor (ampicillin resistance). |
| pBAD_Polycistronic_1_UBE2W_(*D.tinctorium*)_2_MP_TEV_PCNA_K164TAG_H6 (OP1583) | *D. tinctorium* UBE2W followed by PCNA-K164TAG with an N-terminal proline followed by a TEV site with a C-terminal H6-tag under an arabinose promotor (ampicillin resistance). |
| pBAD_Polycistronic_1_UBE2W_(*D.melanogaster*)_2_MP_TEV_PCNA_K164TAG_H6 (OP1584) | *D. melanogaster* UBE2W followed by PCNA-K164TAG with an N-terminal proline followed by a TEV site with a C-terminal H6-tag under an arabinose promotor (ampicillin resistance). |
| pBAD_Polycistronic_1_UBE2W_(*C.elegans*)_2_MP_TEV_PCNA_K164TAG_H6 (OP1585) | *C. elegans* UBE2W followed by PCNA-K164TAG with an N-terminal proline followed by a TEV site with a C-terminal H6-tag under an arabinose promotor (ampicillin resistance). |
| pBAD_Polycistronic_1_UBE2W_(*C.aegerita*)_2_MP_TEV_PCNA_K164TAG_H6 (OP1586) | *C. aegerita* UBE2W followed by PCNA-K164TAG with an N-terminal proline followed by a TEV site with a C-terminal H6-tag under an arabinose promotor (ampicillin resistance). |
| pBAD_Polycistronic_1_UBE2W_(*Z.mays*)_2_MP_TEV_PCNA_K164TAG_H6 (OP1587) | *Z. mays* UBE2W followed by PCNA-K164TAG with an N-terminal proline followed by a TEV site with a C-terminal H6-tag under an arabinose promotor (ampicillin resistance). |
| pBAD_Polycistronic_1_UBE2W_(*P.falciparum*)_2_MP_TEV_PCNA_K164TAG_H6 (OP1588) | *P. falciparum* UBE2W followed by PCNA-K164TAG with an N-terminal proline followed by a TEV site with a C-terminal H6-tag under an arabinose promotor (ampicillin resistance). |
| pBAD_Polycistronic_1_UBE2W_(*H.sapiens*)_2_MP_TEV_SUMO1_wt_deltaG_H6 (OP1488) | *H. sapiens* UBE2W followed by SUMO1-wt with an N-terminal proline followed by a TEV site and a G97 deletion with a C-terminal H6-tag under an arabinose promotor (ampicillin resistance). |
| pBAD_Polycistronic_1_UBE2W_(*H.sapiens*)_2_MP_TEV_SUMO1_K7TAG_deltaG_H6 (OP1488) | *H. sapiens* UBE2W followed by SUMO1-K7TAG with an N-terminal proline followed by a TEV site and a G97 deletion with a C-terminal H6-tag under an arabinose promotor (ampicillin resistance). |
| pBAD_Polycistronic_1_UBE2W_(*H.sapiens*)_2_MP_H8_GST_TEV_NEMO(253-338)_wt (OP1347) | *H. sapiens* UBE2W followed by NEMO-wt (253-338) fused to a GST with an N-terminal proline followed by a H8-tag and a TEV site under an arabinose promotor (ampicillin resistance). |
| pBAD_Polycistronic_1_UBE2W_(*H.sapiens*)_2_MP_H8_GST_TEV_NEMO(253- | *H. sapiens* UBE2W followed by NEMO-K302TAG (253-338) fused to a GST with an N-terminal proline followed by a H8-tag and a TEV site under an arabinose promotor |

| | |
|---|---|
| 338)_K302TAG (OP1402) | (ampicillin resistance). |
| pBAD_Polycistronic_1_UBE2W_(*H.sapiens*)_2_MP_H8_GST_TEV_NEMO(253-338)_K309TAG (OP1349) | *H. sapiens* UBE2W followed by NEMO-K309TAG (253-338) fused to a GST with an N-terminal proline followed by a H8-tag and a TEV site under an arabinose promotor (ampicillin resistance). |
| pBAD_Polycistronic_1_UBE2W_(*H.sapiens*)_2_MP_H6_TEV_Ran_wt (OP1310) | *H. sapiens* UBE2W followed by *H. sapiens* Ran-wt with an N-terminal proline followed by a H6-tag and a TEV site under an arabinose promotor (ampicillin resistance). |
| pBAD_Polycistronic_1_UBE2W_(*H.sapiens*)_2_MP_H6_TEV_Ran_wt (OP1311) | *H. sapiens* UBE2W followed by *H. sapiens* Ran-K71TAG with an N-terminal proline followed by a H6-tag and a TEV site under an arabinose promotor (ampicillin resistance). |

#### 1.2 Amino acid sequences of proteins

Ub

MQIFVKTLTGKTITLEVEPSDTIENVKAKIQDKEGIPPDQQRLIFAGKQLEDGRTLSDY
NIQKESTLHLVLRLRGG

MP-H6-TEV-linear-diUb(G76V)

MPHHHHHHENLYFQGGSMQIFVKTLTGKTITLEVEPSDTIENVKAKIQDKEGIPPDQ
QRLIFAGKQLEDGRTLSDYNIQKESTLHLVLRLRGVMQIFVKTLTGKTITLEVEPSDTI
ENVKAKIQDKEGIPPDQQRLIFAGKQLEDGRTLSDYNIQKESTLHLVLRLRGG

MP-H6-TEV-linear-triUb(G76V)

MPHHHHHHENLYFQGGSMQIFVKTLTGKTITLEVEPSDTIENVKAKIQDKEGIPPDQ
QRLIFAGKQLEDGRTLSDYNIQKESTLHLVLRLRGVMQIFVKTLTGKTITLEVEPSDTI
ENVKAKIQDKEGIPPDQQRLIFAGKQLEDGRTLSDYNIQKESTLHLVLRLRGVMQIFV
KTLTGKTITLEVEPSDTIENVKAKIQDKEGIPPDQQRLIFAGKQLEDGRTLSDYNIQKE
STLHLVLRLRGG

MP-H6-TEV-linear-tetraUb(G76V)

MPHHHHHHENLYFQGGSMQIFVKTLTGKTITLEVEPSDTIENVKAKIQDKEGIPPDQ
QRLIFAGKQLEDGRTLSDYNIQKESTLHLVLRLRGVMQIFVKTLTGKTITLEVEPSDTI
ENVKAKIQDKEGIPPDQQRLIFAGKQLEDGRTLSDYNIQKESTLHLVLRLRGVMQIFV
KTLTGKTITLEVEPSDTIENVKAKIQDKEGIPPDQQRLIFAGKQLEDGRTLSDYNIQKE
STLHLVLRLRGVMQIFVKTLTGKTITLEVEPSDTIENVKAKIQDKEGIPPDQQRLIFAG
KQLEDGRTLSDYNIQKESTLHLVLRLRGG

UBA1

MAKNGSEADIDEGLYSRQLYVLGHEAMKRLQTSSVLVSGLRGLGVEIAKNIILGGVK
AVTLHDQGTAQWADLSSQFYLREEDIGKNRAEVSQPRLAELNSYVPVTAYTGPLVE
DFLSGFQVVVLTNTPLEDQLRVGEFCHNRGIKLVVADTRGLFGQLFCDFGEEMILTD
SNGEQPLSAMVSMVTKDNPGVVTCLDEARHGFESGDFVSFSEVQGMVELNGNQP
MEIKVLGPYTFSICDTSNFSDYIRGGIVSQVKVPKKISFKSLVASLAEPDFVVTDFAKF
SRPAQLHIGFQALHQFCAQHGRPPRPRNEEDAAELVALAQAVNARALPAVQQNNL
DEDLIRKLAYVATGDLAPINAFIGGLAAQEVMKACSGKFMPIMQWLYFDALECLPVD
KEVLTEDKCLQRQNRYDGQVAVFGSDLQEKLGKQKYFLVGAGAIGCELLKNFAMIG
LGCGEGGEIIVTDMDTIEKSNLNRQFLFRPWDVTKLKSDTAAAAVRQMNPHIRVTSH
QNRVGPDTERIYDDDFFQNLDGVANALDNVDARMYMDRRCVYYRKPLLESGTLGT
KGNVQVVIPFLTESYSSSQDPPEKSIPICTLKNFPNAIEHTLQWARDEFEGLFKQPAE
NVNQYLTDPKFVERTLRLAGTQPLEVLEAVQRSLVLQRPQTWADCVTWACHHWHT
QYSNNIRQLLHNFPPDQLTSSGAPFWSGPKRCPHPLTFDVNNPLHLDYVMAAANLF

AQTYGLTGSQDRAAVATFLQSVQVPEFTPKSGVKIHVSDQELQSANASVDDSRLEE
LKATLPSPDKLPGFKMYPIDFEKDDDSNFHMDFIVAASNLRAENYDIPSADRHKSKLI
AGKIIPAIATTTAAVVGLVCLELYKVVQGHRQLDSYKNGFLNLALPFFGFSEPLAAPR
HQYYNQEWTLWDRFEVQGLQPNGEEMTLKQFLDYFKTEHKLEITMLSQGVSMLYS
FFMPAAKLKERLDQPMTEIVSRVSKRKLGRHVRALVLELCCNDESGEDVEVPYVRY
TIR

MP-H6-TEV-UBE2W (*H. sapiens*)

MPGHHHHHHENLYFQGMASMQKRLQKELLALQNDPPPGMTLNEKSVQNSITQWIV
DMEGAPGTLYEGEKFQLLFKFSSRYPFDSPQVMFTGENIPVHPHVYSNGHICLSILT
EDWSPALSVQSVCLSIISMLSSCKEKRRPPDNSFYVRTCNKNPKKTKWWYHDDTC

MP-H6-TEV-SUMO-Ub-UBE2W (*H.sapiens*)

MPGHHHHHHHHHENLYFQGADEKPKEGVKTENNDHINLKVAGQDGSVVQFKIKRHT
PLSKLMKAYCERQGLSMRQIRFRFDGQPINETDTPAQLEMEDEDTIDVFQQQTGGP
MQIFVKTLTGKTITLEVEPSDTIENVKAKIQDKEGIPPDQQRLIFAGKQLEDGRTLSDY
NIQKESTLHLVLRLRGGMASMQKRLQKELLALQNDPPPGMTLNEKSVQNSITQWIV
DMEGAPGTLYEGEKFQLLFKFSSRYPFDSPQVMFTGENIPVHPHVYSNGHICLSILT
EDWSPALSVQSVCLSIISMLSSCKEKRRPPDNSFYVRTCNKNPKKTKWWYHDDTC

MP-Strep-TEV-SUMO-Ub-UBE2W (UBE2W$^{S/U}$) (*H.sapiens*)

MPWSHPQFEKENLYFQGADEKPKEGVKTENNDHINLKVAGQDGSVVQFKIKRHTP
LSKLMKAYCERQGLSMRQIRFRFDGQPINETDTPAQLEMEDEDTIDVFQQQTGGP
MQIFVKTLTGKTITLEVEPSDTIENVKAKIQDKEGIPPDQQRLIFAGKQLEDGRTLSDY
NIQKESTLHLVLRLRGGMASMQKRLQKELLALQNDPPPGMTLNEKSVQNSITQWIV
DMEGAPGTLYEGEKFQLLFKFSSRYPFDSPQVMFTGENIPVHPHVYSNGHICLSILT
EDWSPALSVQSVCLSIISMLSSCKEKRRPPDNSFYVRTCNKNPKKTKWWYHDDTC

H6-USP2

MGSSHHHHHHSSGLVPRGSSSPGRDGMNSKSAQGLAGLRNLGNTCFMNSILQCL
SNTRELRDYCLQRLYMRDLHHGSNAHTALVEEFAKLIQTIWTSSPNDVVSPSEFKT
QIQRYAPRFVGYNQQDAQEFFRFLLDGLHNEVNRVTLRPKSNPENLDHLPDDEKG
RQMWRKYLEREDSRIGDLFVGQLKSSLTCTDCGYCSTVFDPFWDLSLPIAKRGYPE
VTLMDCMRLFTKEDVLDGDAAPTCCRCRGRKRCIKKFSIQRFPKILVLHLKRFSESRI
RTSKLTTFVNFPLRDLDLREFASENTNHAVYNLYAVSNHSGTTMGGHYTAYCRSPG
TGEWHTFNDSSVTPMSSSQVRTSDAYLLFYELASPPSRM

GST-TEV-NTF2

MSPILGYWKIKGLVQPTRLLLEYLEEKYEEHLYERDEGDKWRNKKFELGLEFPNLPY
YIDGDVKLTQSMAIIRYIADKHNMLGGCPKERAEISMLEGAVLDIRYGVSRIAYSKDF

ETLKVDFLSKLPEMLKMFEDRLCHKTYLNGDHVTHPDFMLYDALDVVLYMDPMCLD
AFPKLVCFKKRIEAIPQIDKYLKSSKYIAWPLQGWQATFGGGDHPPKSDLVPRGSEN
LYFQ↓GGSGMGDKPIWEQIGSSFIQHYYQLFDNDRTQLGAIYIDASCLTWEGQQFQ
GKAAIVEKLSSLPFQKIQHSITAQDHQPTPDSCIISMVVGQLKADEDPIMGFHQMFLL
KNINDAWVCTNDMFRLALHNFG

H6-SUMO-RCC1

MHHHHHHGSMDSEVNQEAKPEVKPEVKPETHINLKVSDGSSEIFFKIKKTTPLRRLM
EAFAKRQGKEMDSLRFLYDGIRIQADQTPEDLDMEDNDIIEAHREQIGGGSPKRIAK
RRSPPADAIPKSKKVKVSHRSHSTEPGLVLTLGQGDVGQLGLGENVMERKKPALVS
IPEDVVQAEAGGMHTVCLSKSGQVYSFGCNDEGALGRDTSVEGSEMVPGKVELQE
KVVQVSAGDSHTAALTDDGRVFLWGSFRDNNGVIGLLEPMKKSMVPVQVQLDVPV
VKVASGNDHLVMLTADGDLYTLGCGEQGQLGRVPELFANRGGRQGLERLLVPKCV
MLKSRGSRGHVRFQDAFCGAYFTFAISHEGHVYGFGLSNYHQLGTPGTESCFIPQN
LTSFKNSTKSWVGFSGGQHHTVCMDSEGKAYSLGRAEYGRLGLGEGAEEKSIPTLI
SRLPAVSSVACGASVGYAVTKDGRVFAWGMGTNYQLGTGQDEDAWSPVEMMGK
QLENRVVLSVSSGGQHTVLLVKDKEQS

GST-3C-UBE2N

MSPILGYWKIKGLVQPTRLLLEYLEEKYEEHLYERDEGDKWRNKKFELGLEFPNLPY
YIDGDVKLTQSMAIIRYIADKHNMLGGCPKERAEISMLEGAVLDIRYGVSRIAYSKDF
ETLKVDFLSKLPEMLKMFEDRLCHKTYLNGDHVTHPDFMLYDALDVVLYMDPMCLD
AFPKLVCFKKRIEAIPQIDKYLKSSKYIAWPLQGWQATFGGGDHPPKSDLEVLFQGP
LGSAGLPRRIIKETQRLLAEPVPGIKAEPDESNARYFHVVIAGPQDSPFEGGTFKLEL
FLPEEYPMAAPKVRFMTKIYHPNVDKLGRICLDILKDKWSPALQIRTVLLSIQALLSAP
NPDDPLANDVAEQWKTNEAQAIETARAWTRLYAMNNI*

GST-3C-UBE2V1

MSPILGYWKIKGLVQPTRLLLEYLEEKYEEHLYERDEGDKWRNKKFELGLEFPNLPY
YIDGDVKLTQSMAIIRYIADKHNMLGGCPKERAEISMLEGAVLDIRYGVSRIAYSKDF
ETLKVDFLSKLPEMLKMFEDRLCHKTYLNGDHVTHPDFMLYDALDVVLYMDPMCLD
AFPKLVCFKKRIEAIPQIDKYLKSSKYIAWPLQGWQATFGGGDHPPKSDLEVLFQGP
LGSPGEVQASYLKSQSKLSDEGRLEPRKFHCKGVKVPRNFRLLEELEEGQKGVGD
GTVSWGLEDDEDMTLTRWTGMIIGPPRTIYENRIYSLKIECGPKYPEAPPFVRFVTKI
NMNGVNSSNGVVDPRAISVLAKWQNSYSIKVVLQELRRLMMSKENMKLPQPPEGQ
CYSN*

MP-TEV-SUMO-wt-deltaG-H6

MPENLYFQADEKPKEGVKTENNDHINLKVAGQDGSVVQFKIKRHTPLSKLMKAYCE
RQGLSMRQIRFRFDGQPINETDTPAQLEMEDEDTIDVFQQQTGHHHHHH

MP-TEV-SUMO-K11TAG-deltaG-H6

MPENLYFQADEKPKEGV*TENNDHINLKVAGQDGSVVQFKIKRHTPLSKLMKAYCE
RQGLSMRQIRFRFDGQPINETDTPAQLEMEDEDTIDVFQQQTGHHHHHH*

A-SUMO-H6

MADEKPKEGVKTENNDHINLKVAGQDGSVVQFKIKRHTPLSKLMKAYCERQGLSM
RQIRFRFDGQPINETDTPAQLEMEDEDTIDVFQQQTGGHHHHHH

HA-SUMO-H6

MYPYDVPDYAADEKPKEGVKTENNDHINLKVAGQDGSVVQFKIKRHTPLSKLMKAY
CERQGLSMRQIRFRFDGQPINETDTPAQLEMEDEDTIDVFQQQTGHHHHHH

MP-Ub-TEV-H3-wt-H6

MPGSMQIFVKTLTGKTITLEVEPSDTIENVKAKIQDKEGIPPDQQRLIFAGKQLEDGR
TLSDYNIQKESTLHLVLRLRGGENLYFQARTKQTARKSTGGKAPRKQLATKAARKS
APATGGVKKPHRYRPGTVALREIRRYQKSTELLIRKLPFQRLVREIAQDFKTDLRFQ
SSAVMALQEASEAYLVALFEDTNLCAIHAKRVTIMPKDIQLARRIRGERARSHHHHH
H

MP-Ub-TEV-H3-K27TAG-H6

MPGSMQIFVKTLTGKTITLEVEPSDTIENVKAKIQDKEGIPPDQQRLIFAGKQLEDGR
TLSDYNIQKESTLHLVLRLRGGENLYFQARTKQTARKSTGGKAPRKQLATKAAR*SA
PATGGVKKPHRYRPGTVALREIRRYQKSTELLIRKLPFQRLVREIAQDFKTDLRFQS
SAVMALQEASEAYLVALFEDTNLCAIHAKRVTIMPKDIQLARRIRGERARSHHHHHH

MP-TEV-PCNA-wt-H6

MPENLYFQGFEARLVQGSILKKVLEALKDLINEACWDISSSGVNLQSMDSSHVSLVQ
LTLRSEGFDTYRCDRNLAMGVNLTSMSKILKCAGNEDIITLRAEDNADTLALVFEAPN
QEKVSDYEMKLMDLDVEQLGIPEQEYSCVVKMPSGEFARICRDLSHIGDAVVISCAK
DGVKFSASGELGNGNIKLSQTSNVDKEEEAVTIEMNEPVQLTFALRYLNFFTKATPL
SSTVTLSMSADVPLVVEYKIADMGHLKYYLAPKIEDEEGSHHHHHH

MP-TEV-PCNA-K164TAG-H6

MPENLYFQGFEARLVQGSILKKVLEALKDLINEACWDISSSGVNLQSMDSSHVSLVQ
LTLRSEGFDTYRCDRNLAMGVNLTSMSKILKCAGNEDIITLRAEDNADTLALVFEAPN
QEKVSDYEMKLMDLDVEQLGIPEQEYSCVVKMPSGEFARICRDLSHIGDAVVISCA*
DGVKFSASGELGNGNIKLSQTSNVDKEEEAVTIEMNEPVQLTFALRYLNFFTKATPL
SSTVTLSMSADVPLVVEYKIADMGHLKYYLAPKIEDEEGSHHHHHH

MP-H6-TEV-tau(244-372)-wt

MPHHHHHHGSENLYFQGMQTAPVPMPDLKNVKSKIGSTENLKHQPGGGKVQIINK
KLDLSNVQSKCGSKDNIKHVPGGGSVQIVYKPVDLSKVTSKCGSLGNIHHKPGGGQ
VEVKSEKLDFKDRVQSKIGSLDNITHVPGGGNKKIE

MP-H6-TEV-tau(244-372)-K353TAG

MPHHHHHHGSENLYFQGMQTAPVPMPDLKNVKSKIGSTENLKHQPGGGKVQIINK
KLDLSNVQSKCGSKDNIKHVPGGGSVQIVYKPVDLSKVTSKCGSLGNIHHKPGGGQ
VEVKSEKLDFKDRVQS*IGSLDNITHVPGGGNKKIE

MP-H8-GST-TEV-NEMO(253-338)-wt

MPHHHHHHHHGSMSPILGYWKIKGLVQPTRLLLEYLEEKYEEHLYERDEGDKWRN
KKFELGLEFPNLPYYIDGDVKLTQSMAIIRYIADKHNMLGGCPKERAEISMLEGAVLDI
RYGVSRIAYSKDFETLKVDFLSKLPEMLKMFEDRLCHKTYLNGDHVTHPDFMLYDA
LDVVLYMDPMCLDAFPKLVCFKKRIEAIPQIDKYLKSSKYIAWPLQGWQATFGGGDH
PPKGIEGSENLYFQGERKRGMQLEDLKQQLQQAEEALVAKQEVIDKLKEEAEQHKI
VMETVPVLKAQADIYKADFQAERQAREKLAEKKELLQEQLEQLQR

MP-H8-GST-TEV-NEMO(253-338)-K302TAG

MPHHHHHHHHGSMSPILGYWKIKGLVQPTRLLLEYLEEKYEEHLYERDEGDKWRN
KKFELGLEFPNLPYYIDGDVKLTQSMAIIRYIADKHNMLGGCPKERAEISMLEGAVLDI
RYGVSRIAYSKDFETLKVDFLSKLPEMLKMFEDRLCHKTYLNGDHVTHPDFMLYDA
LDVVLYMDPMCLDAFPKLVCFKKRIEAIPQIDKYLKSSKYIAWPLQGWQATFGGGDH
PPKGIEGSENLYFQGERKRGMQLEDLKQQLQQAEEALVAKQEVIDKLKEEAEQHKI
VMETVPVL*AQADIYKADFQAERQAREKLAEKKELLQEQLEQLQR*

MP-H8-GST-TEV-NEMO(253-338)-K309TAG

MPHHHHHHHHGSMSPILGYWKIKGLVQPTRLLLEYLEEKYEEHLYERDEGDKWRN
KKFELGLEFPNLPYYIDGDVKLTQSMAIIRYIADKHNMLGGCPKERAEISMLEGAVLDI
RYGVSRIAYSKDFETLKVDFLSKLPEMLKMFEDRLCHKTYLNGDHVTHPDFMLYDA
LDVVLYMDPMCLDAFPKLVCFKKRIEAIPQIDKYLKSSKYIAWPLQGWQATFGGGDH
PPKGIEGSENLYFQGERKRGMQLEDLKQQLQQAEEALVAKQEVIDKLKEEAEQHKI
VMETVPVLKAQADIY*ADFQAERQAREKLAEKKELLQEQLEQLQR

MP-H6-TEV-Synuclein-wt

MPHHHHHHGSENLYFQGMDVFMKGLSKAKEGVVAAAEKTKQGVAEAAGKTKEGV
LYVGSKTKEGVVHGVATVAEKTKEQVTNVGGAVVTGVTAVAQKTVEGAGSIAAATG
FVKKDQLGKNEEGAPQEGILEDMPVDPDNEAYEMPSEEGYQDYEPEA

MP-H6-TEV-Synuclein-K10TAG

MPHHHHHHGSENLYFQGMDVFMKGLS*AKEGVVAAAEKTKQGVAEAAGKTKEGVL
YVGSKTKEGVVHGVATVAEKTKEQVTNVGGAVVTGVTAVAQKTVEGAGSIAAATGF
VKKDQLGKNEEGAPQEGILEDMPVDPDNEAYEMPSEEGYQDYEPEA

MP-H6-TEV-Synuclein-K21TAG

MPHHHHHHGSENLYFQGMDVFMKGLSKAKEGVVAAAE*TKQGVAEAAGKTKEGVL
YVGSKTKEGVVHGVATVAEKTKEQVTNVGGAVVTGVTAVAQKTVEGAGSIAAATGF
VKKDQLGKNEEGAPQEGILEDMPVDPDNEAYEMPSEEGYQDYEPEA

MP-TEV-H6-Ran-wt

MPENLYFQGHHHHHHMAAQGEPQVQFKLVLVGDGGTGKTTFVKRHLTGEFEKKY
VATLGVEVHPLVFHTNRGPIKFNVWDTAGQEKFGGLRDGYYIQAQCAIIMFDVTSRV
TYKNVPNWHRDLVRVCENIPIVLCGNKVDIKDRKVKAKSIVFHRKKNLQYYDISAKSN
YNFEKPFLWLARKLIGDPNLEFVAMPALAPPEVVMDPALAAQYEHDLEVAQTTALP
DEDDDL

MP-TEV-H6-Ran-K71TAG

MPENLYFQGHHHHHHMAAQGEPQVQFKLVLVGDGGTGKTTFVKRHLTGEFEKKY
VATLGVEVHPLVFHTNRGPIKFNVWDTAGQE*FGGLRDGYYIQAQCAIIMFDVTSRV
TYKNVPNWHRDLVRVCENIPIVLCGNKVDIKDRKVKAKSIVFHRKKNLQYYDISAKSN
YNFEKPFLWLARKLIGDPNLEFVAMPALAPPEVVMDPALAAQYEHDLEVAQTTALP
DEDDDL

MP-TEV-SUMO1-wt-H6

MPENLYFQSDQEAKPSTEDLGDKKEGEYIKLKVIGQDSSEIHFKVKMTTHLKKLKES
YCQRQGVPMNSLRFLFEGQRIADNHTPKELGMEEEDVIEVYQEQTGHHHHHH

MP-TEV-SUMO1-K7TAG-

MPENLYFQSDQEA*PSTEDLGDKKEGEYIKLKVIGQDSSEIHFKVKMTTHLKKLKESY
CQRQGVPMNSLRFLFEGQRIADNHTPKELGMEEEDVIEVYQEQTGHHHHHH

Nedd8

MLIKVKTLTGKEIEIDIEPTDKVERIKERVEEKEGIPPQQQRLIYSGKQMNDEKTAADY
KILGGSVLHLVLALRGG

Mb-PylRS-wt

MDKKPLDVLISATGLWMSRTGTLHKIKHHEVSRSKIYIEMACGDHLVVNNSRSCRTA
RAFRHHKYRKTCKRCRVSDEDINNFLTRSTESKNSVKVRVVSAPKVKKAMPKSVSR
APKPLENSVSAKASTNTSRSVPSPAKSTPNSSVPASAPAPSLTRSQLDRVEALLSPE
DKISLNMAKPFRELEPELVTRRKNDFQRLYTNDREDYLGKLERDITKFFVDRGFLEIK
SPILIPAEYVERMGINNDTELSKQIFRVDKNLCLRPMLAPTLYNYLRKLDRILPGPIKIF
EVGPCYRKESDGKEHLEEFTMVNFCQMGSGCTRENLEALIKEFLDYLEIDFEIVGDS
CMVYGDTLDIMHGDLELSSAVVGPVSLDREWGIDKPWIGAGFGLERLLKVMHGFKN
IKRASRSESYYNGISTNL

Mb-PylRS(C313V)

MDKKPLDVLISATGLWMSRTGTLHKIKHHEVSRSKIYIEMACGDHLVVNNSRSCRTA
RAFRHHKYRKTCKRCRVSDEDINNFLTRSTESKNSVKVRVVSAPKVKKAMPKSVSR
APKPLENSVSAKASTNTSRSVPSPAKSTPNSSVPASAPAPSLTRSQLDRVEALLSPE
DKISLNMAKPFRELEPELVTRRKNDFQRLYTNDREDYLGKLERDITKFFVDRGFLEIK
SPILIPAEYVERMGINNDTELSKQIFRVDKNLCLRPMLAPTLYNYLRKLDRILPGPIKIF
EVGPCYRKESDGKEHLEEFTMVNFVQMGSGCTRENLEALIKEFLDYLEIDFEIVGDS
CMVYGDTLDIMHGDLELSSAVVGPVSLDREWGIDKPWIGAGFGLERLLKVMHGFKN
IKRASRSESYYNGISTNL

Ma-PylRS(H227I, Y228P)

MTVKYTDAQIQRLREYGNGTYEQKVFEDLASRDAAFSKEMSVASTDNEKKIKGMIA
NPSRHGLTQLMNDIADALVAEGFIEVRTPIFISKDALARMTITEDKPLFKQVFWIDEKR
ALRPMLAPNLYSVMRDLRDHTDGPVKIFEMGSCFRKESHSGMHLEEFTMLNLVDM
GPRGDATEVLKNYISVVMKAAGLPDYDLVQEESDVYKETIDVEINGQEVCSAAVGPI
PLDAAHDVHEPWSGAGFGLERLLTIREKYSTVKKGGASISYLNGAKIN

MP-H6-TEV-UBE2W(*D. rerio*)
MPGHHHHHHENLYFQ↓GMASMQKRLQKELLALQNDPPPGMTLNEKSVQNTITQWI
VDMEGASGTVYEGEKFQLLFKFSSRYPFDSPQVMFTGDNIPVHPHVYSNGHICLSIL
TEDWSPALSVQSVCLSIISMLSSCKEKRRPPDNSFYVRTCNKNPKKTKWWYHDDT
C

MP-H6-TEV-UBE2W(*D. tinctorium*)
MPGHHHHHHENLYFQ↓GMASMQKRLQKELIAIQKEPPPGVSVNPANIGPNLTQWIV
DMEGASGTLYEGEHFQLSFKFSPKYPFDSPEVTFIGSNIPIHPHVYSNGHICLSILSE
DWSPALSVISVCLSIISMLSSCKEKKRPPDNSLYVKTCSKNPKKTKWWYHDDNV

MP-H6-TEV-UBE2W(*D. melongaster*)
MPGHHHHHHENLYFQ↓GMMKLFKKPKDKIPKSEILTEPKEPKTPPVSKCGKPLQLD
NSRWERRLHKELMSLIKEPPPGVTIDTESVQQNLSEWKINIKGFEGTLYEGEDFQLL

FKFNNKYPFDSPEVTFIGTNIPVHPHVYSNGHICLSILTEDWSPALSVQSVCLSIASM
LSSCREKKRPPDNTIYVKTCNKNPKKTKWWYHDDSV

MP-H6-TEV-UBE2W(*C. elegans*)
MPGHHHHHHENLYFQ↓GMSDAATRRLMKELAQLKSEAPEGLLVDNTSTSNDLKQW
KIGVVGAEGTLYAGEVFMLQFTFGPQYPFNSPEVMFVGETIPAHPHIYSNGHICLSIL
SDDWTPALSVQSVCLSILSMLSSSKEKKHPIDDAIYVRTCSKNPSKTRWWFHDDSV

MP-H6-TEV-UBE2W(*C. aegerita*)
MPGHHHHHHENLYFQ↓GMGSVSARRLAKELREIQSEGCPVGITLVDASDFSKWLFT
IEVMGNSQYQGEAYTLQFRFDAQYPISSPAVQFVVTDGKEAPVHPHVYSNGHICASI
LGSEWSPVLSVIAVCVTLQSMLASCKKKERPADNDRYVRTAPDNPKKTLFHYDDDT

MP-H6-TEV-UBE2W(*Z. mays*)
MPGHHHHHHENLYFQ↓GMTSSSSPSRKALSKIACNRLQKELAEWQLSPPAGFKHK
VSDNLQRWVIEVTGAAGTLYAGETYQLQVDFPEHYPMEAPQVIFLNPAPMHPHIYS
NGHICLDILYDSWSPAMTVSSVCISILSMLSSSPAKQRPVDNDRYVRNCRNGRSPK
ETRWWFHDDTV

MP-H6-TEV-UBE2W(*P. falciparum*)
MPGHHHHHHENLYFQ↓GLGNANYRIQKELNNFLKNPPINCTIDVHPSNIRIWIVQYV
GLENTIYANEVYKIKIIFPDNYPLKPPIVYFLQKPPKHTHVYSNGDICLSVLGDDYNPS
LSISGLILSIISMLSSAKEKKLPIDNYTHADAKPGSSQNNFLYHDDKC

### 2. Chemical synthesis

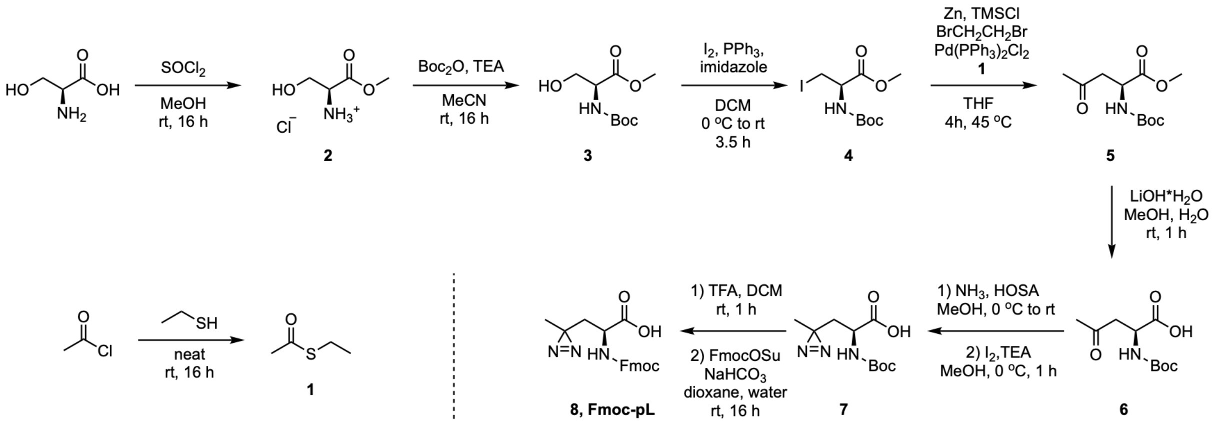

**Scheme S1:** Synthetic route to Fmoc-pL, **8**. *Fmoc: fluorenylmethoxycarbonyl protecting group*

***S*-Ethyl ethanethioate (1).** Acetyl chloride (5.0 mL, 70 mmol) and ethanethiol (5.0 mL, 68 mmol) were combined neat, and the mixture was stirred at rt under nitrogen for 16 h. The reaction was purged with nitrogen for 20 min to remove HCl, affording **1** as a brown liquid (6.3 g, 60 mmol, 86%) that was used in the next step without further purification. $^{1}$H NMR (500 MHz, $CDCl_3$) δ 2.83 (q, *J* = 7.4 Hz, 2H), 2.28 (s, 3H), 1.20 (t, *J* = 7.4 Hz, 3H).

***(S)*-3-Hydroxy-1-methoxy-1-oxopropan-2-aminium chloride (2).** Thionyl chloride (6.9 mL, 95 mmol, 1.0 equiv) was added dropwise to a suspension of L-serine (10 g, 95 mmol) in MeOH (100 mL). The mixture was stirred at rt for 16 h. The reaction was concentrated *in vacuo* to afford **2** as a white solid (14.7 g, 95 mmol, quant.). $^{1}$H NMR (500 MHz, DMSO-$d_6$) δ 8.66 (br s, 3H), 5.62 (s, 1H), 4.04 (t, *J* = 3.5 Hz, 1H), 3.81 (d, *J* = 3.5 Hz, 2H), 3.71 (s, 3H).

**Methyl (*tert*-butoxycarbonyl)-L-serinate (3).** To a mixture of **2** (14.7 g, 95 mmol) in MeCN (225 mL) was added triethylamine (53 mL, 380 mmol, 4.0 equiv). The mixture was stirred at rt for 30 min. Di-*tert*-butyl dicarbonate (20.7 g, 95 mmol, 1.0 equiv) was added portionwise, and the resulting mixture was stirred at rt for 16 h. The reaction was concentrated *in vacuo*, and the resulting oil was dissolved in DCM (250 mL). The organic layer was washed with 1 M aq. HCl (400 mL), dried over anhydrous $Na_2SO_4$, filtered, and concentrated *in vacuo* to afford **3** as a yellow oil (20.8 g, 95 mmol, quant.) that was used in the next step without further purification. $^{1}$H NMR (500 MHz, $CDCl_3$) δ 5.59 (d, *J* = 8.1 Hz, 1H), 4.42-4.24 (m, 1H), 3.91 (dd, *J* = 11.3, 3.9 Hz, 1H), 3.82 (dd, *J* = 11.3, 3.6 Hz, 1H), 3.73 (s, 3H), 3.08 (br s, 1H), 1.40 (s, 9H).

**Methyl (*R*)-2-((*tert*-butoxycarbonyl)amino)-3-iodopropanoate (4).** Triphenylphosphine (14.95 g, 57 mmol, 1.25 equiv) and imidazole (4.04 g, 59 mmol, 1.3 equiv) were dissolved in DCM (100 mL), and the solution was cooled to 0 °C. Iodine (14.47 g, 57 mmol, 1.25 equiv) was added in three portions, and the mixture was stirred at 0 °C for 30 min. A solution of **3** (10 g, 46 mmol) in DCM (100 mL) was added dropwise over 30 min. The resulting mixture was stirred at 0 °C for 1 h, then warmed to rt and stirred for 2 h. The mixture was concentrated *in vacuo*, and the residue was taken up in a minimal volume of DCM and filtered. The filtrate was concentrated *in vacuo*. The crude residue was purified by flash column chromatography (0–15% EtOAc in hexanes) to afford **4** as a light yellow liquid (10.9 g, 33 mmol, 72%). $^1$H NMR (500 MHz, $CDCl_3$) δ 5.36 (d, *J* = 7.8 Hz, 1H), 4.49 (dt, *J* = 8.0, 4.0 Hz, 1H), 3.76 (s, 3H), 3.55 (dd, *J* = 10.3, 3.9 Hz, 1H), 3.52 (dd, *J* = 10.3, 4.0 Hz, 1H), 1.42 (s, 9H).

**Methyl (*S*)-2-((*tert*-butoxycarbonyl)amino)-4-oxopentanoate (5).** A flame-dried flask was charged with zinc dust (2.34 g, 36 mmol, 6.0 equiv) under vacuum and backfilled with $N_2$. THF (8.0 mL) and 1,2-dibromoethane (173 µL, 2.0 mmol, 0.33 equiv) were added, and the mixture was stirred at 45 °C under $N_2$ for 20 min. The suspension was cooled to rt, and TMSCl (51 µL, 0.40 mmol, 0.067 equiv) was added. The mixture was stirred for 30 min at rt and heated to 45 °C. A solution of **4** (1.97 g, 6.0 mmol) in THF (8.0 mL) was added, and the mixture was stirred at 45 °C for 30 min. A solution of **1** (938 mg, 9.0 mmol, 1.5 equiv) in THF (2.0 mL) and $PdCl_2(PPh_3)_2$ (211 mg, 0.30 mmol, 5.0 mol %) in THF (5.0 mL) were added sequentially. The resulting mixture was stirred at 45 °C for 4 h. The reaction was quenched with EtOAc (5.0 mL) and filtered through a pad of Celite. The filtrate was concentrated *in vacuo*, and the residue was dissolved in EtOAc (50 mL). The organic layer was washed with 1 M aq. HCl (50 mL) and brine (50 mL), dried over anhydrous $Na_2SO_4$, filtered, and concentrated *in vacuo*. The crude residue was purified by flash column chromatography (0–45% EtOAc in hexanes) to afford **5** as an inseparable mixture with Boc-Ala-OMe (orange oil, 452 mg, 1.84 mmol, 31%, recalculated from HNMR), which was used in the next step without further purification. $^1$H NMR (400 MHz, $CDCl_3$) δ 5.47 (d, *J* = 8.9 Hz, 1H), 4.49 (dd, *J* = 8.7, 4.4 Hz, 1H), 3.73 (s, 3H), 3.17 (dd, *J* = 18.2, 4.4 Hz, 1H), 2.95 (dd, *J* = 18.2, 4.3 Hz, 1H), 2.16 (s, 3H), 1.44 (s, 9H).

***(S)*-2-((*tert*-Butoxycarbonyl)amino)-4-oxopentanoic acid (6). 5** (1.25 g, 5.1 mmol) was dissolved in MeOH (15 mL), and the solution was cooled to 0 °C. Water (5.0 mL) and 1 M aq. LiOH (10 mL, 10 mmol, 2.0 equiv) were added, and the mixture was stirred at rt for 1 h. The pH of the reaction was adjusted to 3–4 with 1 M aq. HCl, and the MeOH was removed *in vacuo*. The aqueous residue was extracted with EtOAc (2 × 10 mL). The combined organic layers were washed with brine (10 mL), dried over anhydrous $Na_2SO_4$, filtered, and concentrated *in vacuo*. The crude residue was purified by flash column chromatography (30–100% EtOAc in hexanes) to afford **6** as a colorless oil (1.06 g, 4.6 mmol, 90%). $^1$H NMR (500 MHz, $CDCl_3$) δ 10.53 (br s, 1H), 5.58 (d, *J* = 8.7 Hz, 1H), 4.42 (dt, *J* = 8.9, 4.6 Hz, 1H), 3.07 (dd, *J* = 18.3, 4.6 Hz, 1H), 2.89 (dd, *J* = 17.9, 4.5 Hz, 1H), 2.08 (s, 3H), 1.32 (s, 9H).

***(S)-2-((tert-Butoxycarbonyl)amino)-3-(3-methyl-3H*-diazirin-3-yl)propanoic acid (7).** To **6** (1.06 g, 4.6 mmol) at 0 °C was added 7 N ammonia in MeOH (15 mL). The mixture was stirred at 0 °C for 5 h. A solution of hydroxylamine-*O*-sulfonic acid (HOSA, 625 mg, 5.5 mmol, 1.2 equiv) in MeOH (3.0 mL) was added dropwise over 20 min at 0 °C. The mixture was stirred at 0 °C for 3 h, then warmed to rt and stirred for 16 h. The reaction was filtered, and the filtrate was concentrated *in vacuo* to completely remove ammonia. The residue was dissolved in MeOH (5.0 mL), and triethylamine (1.39 mL, 10 mmol, 2.2 equiv) was added. The mixture was cooled to 0 °C, and a saturated solution of iodine in MeOH was added dropwise until an orange color persisted. The mixture was concentrated *in vacuo*, and the residue was dissolved in EtOAc (50 mL). The organic layer was washed with 1 M aq. HCl (50 mL) and brine (50 mL), dried over anhydrous $Na_2SO_4$, filtered, and concentrated *in vacuo*. The crude residue was purified by flash column chromatography (0–2.5% MeOH in DCM with 0.1% AcOH) to afford **7** as a yellow oil (0.35 g, 1.4 mmol, 31%). $^1$H NMR (500 MHz, CDCl3) δ 10.66 (br s, 1H), 5.13 (d, *J* = 8.3 Hz, 1H), 4.37 (td, *J* = 7.9, 4.7 Hz, 1H), 2.04 (dd, *J* = 15.0, 5.0 Hz, 1H), 1.59 (dd, *J* = 13.1, 8.4 Hz, 1H), 1.46 (s, 9H), 1.08 (s, 3H). HRMS (ESI-QTOF) *m/z*: [M + Na]$^+$ calcd for $C_{10}H_{17}N_3O_4Na$ 266.1111; found 266.1107.

**(*S*)-2-((((9*H*-fluoren-9-yl)methoxy)carbonyl)amino)-3-(3-methyl-3*H*-diazirin-3-yl)propanoic acid (8). 7** (0.35 g, 1.4 mmol) was dissolved in TFA/DCM (1:1, v/v) containing 1% $H_2O$ (5.0 mL), and the mixture was stirred at rt for 1 h. The reaction was concentrated *in vacuo*, and the residue was dissolved in 1,4-dioxane/$H_2O$ (1:1, v/v, 10 mL). $NaHCO_3$ (588 mg, 7.0 mmol, 5.0 equiv) and Fmoc-OSu (708 mg, 2.1 mmol, 1.5 equiv) were added sequentially. The mixture was stirred at rt for 16 h. The 1,4-dioxane was removed *in vacuo*, and the aqueous residue was acidified to pH 3–4 with 1 M aq. HCl, followed by extraction with EtOAc (2 × 15 mL). The combined organic layers were washed with brine (15 mL), dried over anhydrous $Na_2SO_4$, filtered, and concentrated *in vacuo*. The crude residue was purified by flash column chromatography (0–3.5% MeOH in DCM with 0.1% AcOH) to afford **8** as a colorless oil (0.41 g, 1.1 mmol, 79%). $^1$H NMR (500 MHz, DMSO-$d_6$) δ 12.81 (br s, 1H), 7.90 (dt, *J* = 7.6, 0.9 Hz, 2H), 7.78-7.72 (m, 3H), 7.45-7.38 (m, 2H), 7.33 (tdd, *J* = 7.5, 2.6, 1.2 Hz, 2H), 4.46-4.13 (m, 3H), 3.84 (ddd, *J* = 10.8, 8.5, 4.2 Hz, 1H), 1.89 (dd, *J* = 14.9, 4.2 Hz, 1H), 1.66 (dd, *J* = 14.9, 10.8 Hz, 1H), 1.02 (s, 3H). HRMS (ESI-QTOF) *m/z*: [M + Na]$^+$ calcd for $C_{20}H_{19}N_3O_4Na$ 388.1268; found 388.1265.

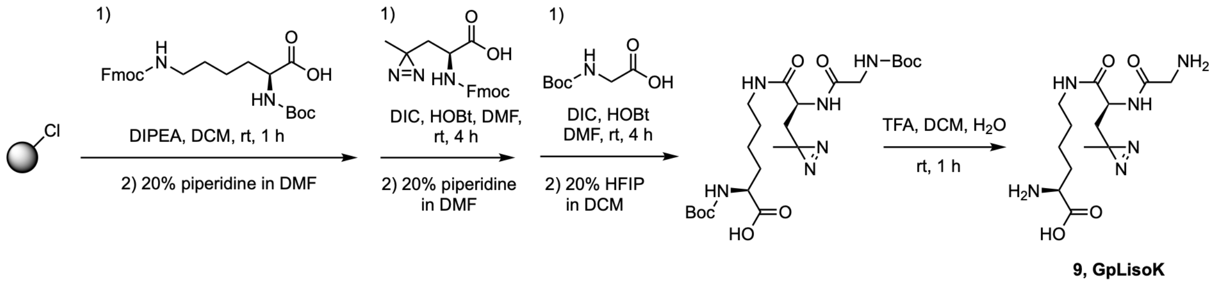

**Scheme S2:** Synthetic route to GpLisoK, **9.**

**GpLisoK (9).** 2-Chlorotrityl chloride (CTC) resin (1.72 g, 2.8 mmol, 1.63 mmol/g) was loaded into a fritted column under $N_2$ and washed with dry DCM (40 mL). A solution of Boc-Lys(Fmoc)-OH (3.0 g, 6.4 mmol, 2.3 equiv) and DIPEA (1.95 mL, 11.2 mmol, 4.0 equiv) in DCM (40 mL) was added, and the mixture was agitated for 1 h. The resin was washed with DCM (5 × 40 mL) and DMF (3 × 40 mL). The Fmoc group was removed by treating the resin with 20% piperidine in DMF (40 mL) twice for 15 min, with DMF washes (2 × 40 mL) between treatments. The resin was then washed with DMF (5 × 40 mL).

Prior to coupling, the resin was washed with dry DMF (2 × 40 mL) under $N_2$ and allowed to swell in dry DMF (20 mL). A solution of **8** (1.53 g, 4.2 mmol, 1.5 equiv) and HOBt (0.814 g, 6.0 mmol, 2.1 equiv) in dry DMF (40 mL) was added, followed by DIC (1.32 mL, 8.4 mmol, 3.0 equiv). The mixture was agitated for 4 h, then washed with DMF (5 × 40 mL). Fmoc deprotection and subsequent washing were performed exactly as described above.

Prior to the final coupling, the resin was washed with dry DMF (2 × 40 mL) under $N_2$ and allowed to swell in dry DMF (20 mL). A solution of Boc-Gly-OH (1.47 g, 8.4 mmol, 3.0 equiv) and HOBt (0.814 g, 6.0 mmol, 2.1 equiv) in DMF (40 mL) was added, followed by DIC (1.55 mL, 10 mmol, 3.6 equiv). The mixture was agitated for 4 h. The resin was washed with DMF (5 × 40 mL) and DCM (5 × 40 mL).

The peptide was cleaved from the resin by treatment with 20% HFIP in DCM (40 mL) three times for 15 min, with DCM washes (40 mL) between treatments. The combined washes were concentrated *in vacuo*. The residue was dissolved in DCM (60 mL) and water (1.0 mL). TFA (40 mL) was added slowly, and the mixture was stirred at rt for 1 h. All solvents were removed *in vacuo*, and the aqueous residue was lyophilized to afford **9** as a colorless solid (650 mg, 2.0 mmol, 71%). $^1$H NMR (400 MHz, DMSO-$d_6$) δ 8.70 (d, $J$ = 8.5 Hz, 1H), 8.31 (s, 3H), 8.24 (t, $J$ = 5.6 Hz, 1H), 8.13 (s, 3H), 4.26 (m, 1H), 3.86 (m, 1H), 3.71–3.44 (m, 2H), 3.17 (q, $J$ = 6.6 Hz, 2H), 1.90–1.67 (m, 3H), 1.60–1.24 (m, 5H), 1.02 (s, 3H). HRMS (ESI-QTOF) *m/z*: $[M + H]^+$ calcd for $C_{13}H_{25}N_6O_4$ 329.1932; found 329.1930.

### 3. Protein expression and purification

#### 3.1 Expression and purification of tagless Ub

Chemically competent *E. coli* Rosetta2 (DE3) were transformed with pET17b-Ub plasmid (see Supplementary Table S1). After recovery with 1 mL of SOC medium for 1 h at 37 °C, the cells were cultured overnight in 50 mL of 2× YT medium containing ampicillin (100 µg/mL), chloramphenicol (50 µg/mL) and 1 % glucose at 37 °C, 200 rpm. The overnight culture was diluted to an $OD_{600}$ (optical density at 600 nm) of 0.05 in 3 L of fresh 2× YT medium supplemented with ampicillin (50 µg/mL) and chloramphenicol (25 µg/mL) and cultured at 37 °C with shaking (200 rpm) until $OD_{600}$ reached 0.8 - 1.0. IPTG (isopropyl β-D-1-thiogalactopyranoside) was added to a final concentration of 1 mM and protein expression was induced for 3 h at 37 °C. The cells were harvested by centrifugation (4000 × *g*, 40 min, 4 °C) and resuspended in lysis buffer (50 mM Tris pH 7.6, 10 mM $MgCl_2$, 1 mM EDTA (ethylenediaminetetraacetic acid)). The cell suspension was lysed by high pressure cell disruption using a (Constant Systems CF1) 2x 30.000 psi followed by centrifugation (14,000 × *g* 40 min, 4°C). The cleared lysate was transferred into a glass beaker in an ice-water bath placed on a magnetic stirrer. Precipitation was performed with 35 % perchloric acid until pH 4.0 - 4.5 was reached. After 5 min incubation at 4 °C while stirring, the milky solution was centrifuged (14,000 × *g*, 40 min, 4 °C) and the supernatant was transferred into a dialysis tubing with a MWCO (molecular weight cut-off) of 2 kDa (Roth). Dialysis was performed overnight at 4 °C with 50 mM ammonium acetate buffer pH 4.5. The dialyzed solution was centrifuged (14,000 × *g*, 40 min, 4 °C), filtered and purified via a HiTrap SP FF 5 mL cation exchange chromatography (GE, gradient 0 - 1 M NaCl). Fractions that showed > 95% purity, as judged by sodium dodecyl sulfate-polyacrylamide gel electrophoresis (SDS-PAGE), were pooled, concentrated, and further purified via size-exclusion chromatography (SEC) using a Superdex Increase 75 10/300 (GE Healthcare) with a buffer containing 25 mM Tris pH 7.5 at 4 °C, 150 mM NaCl. Fractions containing pure Ub were pooled together and concentrated using Amicon® centrifugal filter units with a 3 kDa MWCO (Millipore). Protein concentration was calculated from the measured A280 absorption (extinction coefficients were calculated with ProtParam (https://web.expasy.org/protparam/)). In case of Ub the determination of protein concentration using the absorption at 280 nm is inaccurate (due to the low extinction coefficient (ε)), and therefore bicinchoninic acid assay (BCA) (Thermo Scientific™) and Bradford (Sigma-Aldrich) assays were used for accurate protein concentration determination or the concentration of purified Ub was adjusted densiometrically. Purified Ub was flash frozen using liquid nitrogen and stored at -80 °C until further use.

#### 3.2 Expression and purification of linear di/tri/tetraUb

Chemically competent E. coli K12 were transformed with pBAD-di/tri/tetraUb (encoding for MP-H6-TEV-di/tri/tetraUb with G76V mutation(s) at the distal Ub moieties, see Supplementary Table S1). After recovery with 1 mL of SOC medium for 1 h at 37 °C, the cells were cultured overnight in 50 mL of 2× YT medium containing ampicillin (100 µg/mL) at 37 °C, 200 rpm. The overnight culture was diluted to an OD600 of 0.05 in 200 mL of fresh 2× YT medium supplemented with ampicillin (50 µg/mL) and cultured at 37 °C with shaking (200 rpm) until OD600 = 0.6. Arabinose was added to a final concentration of 0.02 % (w/v) and protein expression was induced for 3 h at 37 °C. The cells were harvested by centrifugation (4000 × g, 10 min, 4 °C) and resuspended in lysis buffer (20 mM Tris pH 8.0, 300 mM NaCl, 30 mM Imidazole, 1 mM PMSF). Afterwards, the cell suspension was incubated on ice for 30 min and sonicated with cooling in an ice-water bath. The lysed cells were centrifuged (14,000 × g, 20 min, 4 °C) and the clear lysate was added to 1 mL Ni-NTA slurry/1 L culture (Cytiva) equilibrated with wash buffer (20 mM Tris pH 8.0, 300 mM NaCl and 30 mM imidazole). Afterwards, the mixture was incubated with agitation for 1 h at 4 °C. After incubation, the mixture was transferred to an empty plastic column and washed with 10 column volumes (CV) of wash buffer. The protein was eluated in 1 mL fractions with wash buffer supplemented with 300 mM imidazole pH 8.0. The fractions containing the di/tri/tetraUb were pooled and concentrated using Amicon centrifugal filter units (Millipore) with a suitable molecular weight cutoff (MWCO) followed by size-exclusion chromatography (SEC) using a Superdex S75 10/300 (Cytiva) with SEC buffer (25 mM Tris pH 7.5, 100 mM NaCl). Fractions containing di/tri/tetraUb variants were pooled and concentrated. Protein concentration was calculated from the measured A280 absorption (extinction coefficients were calculated with ProtParam (https://web.expasy.org/protparam/)). Di/tri/tetraUb were flash frozen using liquid nitrogen and stored at −80 °C until further use.

#### 3.3 Expression and purification of H6-UBE2W and H6-SUMO-Ub-UBE2W

Chemically competent E. coli K12 were transformed with pBAD-H6-UBE2W_Hs (encoding for H. sapiens MP-H6-TEV-UBE2W), pBAD-H6-UBE2W_Dr (encoding for D. rerio MP-H6-TEV-UBE2W) or pBAD-H8-SUMO-Ub-UBE2W_Hs (encoding for H. sapiens MP-H8-TEV-SUMO2-Ub-UBE2W) see Supplementary Table S1. After recovery with 1 mL of SOC medium for 1 h at 37 °C, the cells were cultured overnight in 50 mL of 2× YT medium containing ampicillin (100 µg/mL) at 37 °C, 200 rpm. The overnight culture was diluted to an OD600 of 0.05 in 1.5 L of fresh 2× YT medium supplemented with ampicillin (50 µg/mL) and cultured at 37 °C with shaking (200 rpm) until OD600 = 0.6. Arabinose was added to a final concentration of 0.02 % (w/v) and protein expression was induced for 3 h at 37 °C. The cells were harvested by centrifugation (4000 × g, 40 min, 4 °C) and resuspended in lysis buffer (20 mM Tris pH 8.0, 300 mM NaCl, 30 mM Imidazole, 0.5 mM DTT and 1 mM PMSF). Afterwards, the cell suspension was incubated on ice for 30 min and sonicated with cooling in an ice-water bath. The lysed cells were centrifuged (14,000 × g, 20 min, 4 °C) and the clear lysate was added to 1 mL Ni-NTA slurry/1 L culture (Cytiva) equilibrated with wash buffer (20 mM Tris pH 8.0, 300 mM NaCl, 30 mM imidazole and 0.5 mM DTT). Afterwards, the mixture was incubated with agitation for 1 h at 4 °C. After incubation, the mixture was transferred to an empty plastic column and washed with 10 column volumes (CV) of wash buffer. The protein was eluated in 1 mL fractions with wash buffer supplemented with 300 mM imidazole pH 8.0. The fractions containing the UBE2W variants were pooled and concentrated using Amicon centrifugal filter units (Millipore) with a suitable molecular weight cutoff (MWCO) followed by size-exclusion chromatography (SEC) using a Superdex S75 16/600 (Cytiva) with SEC buffer (25 mM Tris pH 7.5, 100 mM NaCl, 1 mM DTT). Fractions containing UBE2W variants were pooled and concentrated. Protein concentration was calculated from the measured A280 absorption (extinction coefficients were calculated with ProtParam (https://web.expasy.org/protparam/)). Di/tri/tetraUb were flash frozen using liquid nitrogen and stored at −80 °C until further use.

#### 3.4 Expression and purification of Strep-SUMO-Ub-UBE2W

Chemically competent E. coli K12 were transformed with pBAD-Strep-SUMO-Ub-UBE2W_Hs (encoding for H. sapiens MP-Strep-TEV-SUMO2-Ub-UBE2W, see Supplementary Table S1). After recovery with 1 mL of SOC medium for 1 h at 37 °C, the cells were cultured overnight in 50 mL of 2× YT medium containing ampicillin (100 µg/mL) at 37 °C, 200 rpm. The overnight culture was diluted to an OD600 of 0.05 in 200 mL of fresh 2× YT medium supplemented with ampicillin (50 µg/mL) and cultured at 37 °C with shaking (200 rpm) until OD600 = 0.6. Arabinose was added to a final concentration of 0.02 % (w/v) and protein expression was induced for 3 h at 37 °C. The cells were harvested by centrifugation (4000 × g, 10 min, 4 °C) and resuspended in lysis buffer (20 mM Tris pH 8.0, 100 mM NaCl, 1 mM PMSF). Afterwards, the cell suspension was incubated on ice for 30 min, sonicated with cooling in an ice-water bath and the lysate was cleared by centrifugation (14,000 × g, 20 min, 4 °C). The cleared lysate was purified using a 1 mL StrepTrap XT Column (Cytiva) using a gradient (0-100%) of Buffer A (20 mM Tris pH 7.5, 100 mM NaCl) and Buffer B (Buffer A supplemented with 50 mM Biotin). The fractions containing the Strep-SUMO-Ub-UBE2W were pooled and concentrated using Amicon centrifugal filter units (Millipore) with a suitable molecular weight cutoff (MWCO) followed by size-exclusion chromatography (SEC) using a Superdex S75 16/600 (Cytiva) with SEC buffer (10 mM Hepes pH 7.4, 100 mM NaCl,). Fractions containing Strep-SUMO-Ub-UBE2W were pooled and concentrated. Protein concentration was calculated from the measured A280 absorption (extinction coefficients were calculated with ProtParam (https://web.expasy.org/protparam/)). Strep-SUMO-Ub-UBE2W was flash frozen using liquid nitrogen and stored at −80 °C until further use.

#### 3.5 Expression and purification of GST-tagged proteins (GST-UBE2N, GST-UBE2V1, and GST-NTF2)

Chemically competent E. coli Rosetta2 (DE3) were transformed with pGEX-6P-1-UBE2N (encoding for GST-UBE2N), pGEX6P1-UBE2V1 (encoding for GST-UBE2V1), or pGEX-NTF2 (encoding for GST-NTF2) see Supplementary Table S1. After recovery

with 1 mL of SOC medium for 1 h at 37 °C, the cells were cultured overnight in 50 mL of 2× YT medium containing ampicillin (100 µg/mL) and chloramphenicol (50 µg/mL) at 37 °C, 200 rpm. The overnight culture was diluted to an OD600 of 0.05 in 1 L of fresh 2× YT medium supplemented with ampicillin (50 µg/mL) and chloramphenicol (25 µg/mL) and cultured at 37 °C with shaking (200 rpm) until OD600 = 0.8. IPTG was added to a final concentration of 0.25 mM and protein expression was induced for 18 h at 20 °C. The cells were harvested by centrifugation (4000 × g, 40 min, 4 °C) and resuspended in lysis buffer (50 mM Tris pH 8.0, 300 mM sucrose, 50 mM NaF, 2 mM DTT, 0.1 mg/mL DNase I, and one cOmpleteTM protease inhibitor tablet (Roche)). The cell suspension was lysed by high pressure cell disruption using a (Constant Systems CF1) 2x 30.000 psi followed by centrifugation (14,000 × g 40 min, 4°C). The lysed cells were centrifuged (15,000 × g, 40 min, 4 °C), the cleared lysate added to Glutathione Sepharose 4B (GE Healthcare) (0.1 mL of slurry per 100 mL of culture) and the mixture was incubated with agitation for 1 h at 4 °C. After incubation, the mixture was transferred to an empty plastic column and washed with 10 CV of wash buffer (25 mM Tris pH 8.5, 400 mM NaCl, 5 mM DTT). The protein was eluated in 1 mL fractions with elution buffer (wash buffer supplemented with 10 mM GSH pH 8.0). The fractions containing GST-tagged proteins were pooled and concentrated with Amicon® centrifugal filter units with a 10 kDa MWCO (Millipore) followed by SEC using a Superdex S75 16/600 (GE Healthcare) with SEC buffer (50 mM Tris pH 7.5, 150 mM NaCl, 1 mM DTT). Fractions containing the GST-tagged proteins were pooled and concentrated. Protein concentration was calculated from the measured A280 absorption (extinction coefficients were calculated with ProtParam (https://web.expasy.org/protparam/)). GST-tagged proteins were flash frozen using liquid nitrogen and stored at −80 °C until further use.
Facultative GST-cleavage was performed by diluting GST-tagged E2-enzymes to 50 µM in SEC buffer (50 mM Tris pH 7.5, 150 mM NaCl, 1 mM DTT) followed by addition of PreScission Protease (Sigma–Aldrich, Cat. No. GE27-0843-01) or TEV Protease (Sigma–Aldrich, Cat. No. T4455). After incubation at 4 °C for 1 h preequilibrated Glutathione Sepharose 4B (GE Healthcare) was added to remove GST and PreScission protease. The flow through was collected, concentrated and applied to SEC using a Superdex S75 16/600 (GE Healthcare) with SEC buffer. Fractions containing the untagged proteins were pooled and concentrated. Protein concentration was calculated from the measured A280 absorption (extinction coefficients were calculated with ProtParam (https://web.expasy.org/protparam/)). E2-enzymes and NTF2 were flash frozen using liquid nitrogen and stored at −80 °C until further use.

#### 3.6 Expression and purification of RCC1

Chemically competent *E. coli* BL21 (DE3) were transformed with pET13M-H6-SUMO-RCC1 plasmid (see Supplementary Table S1). After recovery with 1 mL of SOC medium for 1 h at 37 °C, the cells were cultured overnight in 50 mL of 2× YT medium containing kanamycin (50 µg/mL) and 1 % glucose at 37 °C, 200 rpm. The overnight culture was diluted to an $OD_{600}$ (optical density at 600 nm) of 0.05 in 200 mL of fresh 2× YT medium supplemented with kanamycin (25 µg/mL) and cultured at 37 °C with

shaking (200 rpm) until $OD_{600}$ reached 0.6 IPTG (isopropyl β-D-1-thiogalactopyranoside) was added to a final concentration of 0.5 mM and protein expression was induced for 4 h at 37 °C. The cells were harvested by centrifugation (4000 × g, 10 min, 4 °C) and resuspended in lysis buffer (20 mM Tris pH 8.0, 300 mM KCl, 30 mM Imidazole, 2 mM $MgCl_2$, 1 mM TCEP, 10% Glycerol). Afterwards, the cell suspension was incubated on ice for 30 min and sonicated with cooling in an ice-water bath. The lysed cells were centrifuged (14,000 × g, 20 min, 4 °C) and the clear lysate was added to 1 mL Ni-NTA slurry/1 L culture (Cytiva) equilibrated with lysis buffer. Afterwards, the mixture was incubated with agitation for 1 h at 4 °C. After incubation, the mixture was transferred to an empty plastic column and washed with 10 column volumes (CV) of lysis buffer. The protein was eluated in 1 mL fractions with lysis buffer supplemented with 300 mM imidazole pH 8.0. The fractions containing H6-SUMO-RCC1 were pooled, and SUMO protease was added followed by dialysis into lysis buffer at 4 °C overnight. Reverse Ni-NTA purification was performed as described above and the flow through was collected and
concentrated using Amicon centrifugal filter units (Millipore) with a suitable molecular weight cutoff (MWCO) followed by size-exclusion chromatography (SEC) using a Superdex S75 10/300 (Cytiva) with SEC buffer (25 mM Tris pH 7.5, 150 mM KCl, 2 mM $MgCl_2$, 10% Glycerol). Fractions containing RCC1 were pooled and concentrated. Protein concentration was calculated from the measured A280 absorption (extinction coefficients were calculated with ProtParam (https://web.expasy.org/protparam/)). was flash frozen using liquid nitrogen and stored at −80 °C until further use.

#### 3.7 Expression and purification of H6-tagged XisoK-bearing proteins (SUMO2, PCNA, tau, synuclein, Ub-H3, and Ran)

Chemically competent E. coli K12 cells were co-transformed with pBAD_POI (which encodes H6-tagged POI with a TAG codon at the denoted position, see Supplementary Table S1) and pEVOL_aaRS (which encodes either the Ma "IP" aaRS/tRNA pair for incorporation of LisoK/pLisoK or Mb wt aaRS/tRNA pair for incorporation of all other XisoKs, see Supplementary Table S1) plasmids.
After recovery with 1 mL of SOC medium for 1 h at 37 °C, the cells were cultured overnight in 5 mL of 2× YT medium containing ampicillin (100 µg/mL) and chloramphenicol (50 µg/mL) at 37 °C, 200 rpm. The overnight culture was diluted to an OD600 of approx. 0.05 in 100 mL autoinduction medium[10] supplemented with ampicillin (50 µg/mL), chloramphenicol (25 µg/mL) and the corresponding ncAA (0.5 mM) and cultured at 37 °C with shaking (200 rpm) overnight. The cells were harvested by centrifugation (4000 × g, 10 min, 4 °C) and resuspended in lysis buffer (20 mM Tris pH 8.0, 300 mM NaCl, 30 mM Imidazole and 1 mM PMSF). Afterwards, the cell suspension was incubated on ice for 30 min and sonicated with cooling in an ice-water bath. The lysed cells were centrifuged (14,000 × g, 20 min, 4 °C) and the clear lysate was added to 1 mL Ni-NTA slurry/1 L culture (Cytiva) equilibrated with wash buffer (20 mM Tris pH 8.0, 300 mM NaCl, 30 mM imidazole). Afterwards, the mixture was incubated with agitation for 1 h at 4 °C. After incubation, the mixture was transferred to an empty plastic column and washed with 10 column volumes (CV) of wash buffer.

The protein was eluated in 1 mL fractions with wash buffer supplemented with 300 mM imidazole pH 8.0. The fractions containing the POI were pooled and concentrated using Amicon centrifugal filter units (Millipore) with a suitable molecular weight cutoff (MWCO) followed by size-exclusion chromatography (SEC) using a Superdex S75 10/300 (Cytiva) with SEC buffer (20 mM Tris pH 7.5, 150 mM NaCl, 1 mM DTT). Fractions containing POI variants were pooled and concentrated. Protein concentration was calculated from the measured A280 absorption (extinction coefficients were calculated with ProtParam (https://web.expasy.org/protparam/)). POIs were flash frozen using liquid nitrogen and stored at −80 °C until further use.

This protocol was used for all SUMO2-K11XisoK variants and PCNA-K164XisoK.

For purification of synuclein variants buffers were adapted accordingly (lysis buffer: 20 mM Tris pH 7.5 and 20 mM Imidazole; SEC Buffer: 20 mM Tris pH 7.5).

For purification of tau variants a heat precipitation step was included by incubating the clarified lysate in an 85 °C water bath for 10 min followed by centrifugation (14,000 × g, 20 min, 4 °C) prior to addition to Ni-NTA slurry.

Ub-H3 was purified from inclusion bodies and refolded. Therefore, the insoluble pellet after sonication and lysate clearance was resuspended in Buffer A (50 mM Tris pH 7.5, 500 mM NaCl, 30 mM Imidazole, 6 M Urea) followed by sonication and centrifugation (14,000 × g, 20 min, 4 °C) and applied to Ni-NTA slurry. After a 10 CV wash step with Buffer A the protein was eluated with Buffer A supplemented with 300 mM Imidazole followed by refolding via dialysis against 20 mM Tris pH 7.5, 100 mM NaCl (six steps).

For purification of Ran buffers were adapted accordingly (lysis buffer: 20 mM Tris pH 7.5, 300 mM NaCl, 20 mM Imidazole, 10 % Glycerol, 5 mM MgCl2, 1 mM TCEP and 1 mM PMSF; SEC Buffer: 20 mM Tris pH 7.5, 100 mM NaCl, 5 mM MgCl2, 1 mM TCEP, 10 % Glycerol). Cleavage of N-terminal MP-TEV was performed after SEC using TEV protease in SEC buffer (12 h, 4 °C) followed by SEC purification.

#### 3.8 Expression and purification of H6-tagged wt proteins (SUMO2, PCNA, tau, synuclein, Ub-H3, and Ran)

Expression and purification of wt proteins was performed identical to their ncAA-bearing counterparts described in 3.7. with the exception that no ncAAs were added to the expression medium.

#### 3.9 Expression and purification of USP2

USP2 was expressed and purified as previously described.[11]

#### 3.10 *In vitro* UBE2W-mediated ubiquitylation of POIs

In vitro UBE2W-mediated ubiquitylation of POIs was performed in UBE2W reaction buffer (50 mM HEPES pH 7.1, 100 mM NaCl, 10 mM ATP, 10 mM $MgCl_2$, 1 mM DTT) at 37°C using optimized reaction conditions: e.g. 40 µM POI, 40 µM UBE2W variant, 100 nM UBA1, 100 – 400 µM Ub. Exact conditions for each POI and Ub donor (mono/di/tri/tetraUb or K63-linked diUb) are given in the according figure legends. Reaction mixtures were incubated at 37 °C and samples were taken at the denoted time points and quenched by the addition of 4× SDS loading buffer. After boiling at 95 °C for 5 min and centrifugation (21,000 × g, 5 min) samples were loaded on 4 - 12% SDS-PAGE gels and visualized by Coomassie staining.

#### 3.11 Preparative in vitro ubiquitylation of SUMO2-K11LisoK

Preparative in vitro UBE2W-mediated ubiquitylation of SUMO2-K11LisoK was performed as described above (40 µM SUMO2-K11LisoK, 40 µM Strep-SUMO-Ub-UBE2W, 50 nM UBA1, 150 µM Ub in UBE2W reaction buffer at 37 °C). After overnight incubation 400 µM phenylvinyl-sulfon was added to the reaction followed by incubation for 30 min at 37 °C. Afterwards the reaction was diluted 10-fold into Ni-NTA wash buffer (20 mM Tris pH 7.5, 300 mM NaCl, 30 mM Imidazole) and 100 uL of washed Ni-NTA slurry was added followed by incubation with agitation for 1 h at 4 °C. After incubation, the mixture was transferred to an empty plastic column and washed with 10 column volumes (CV) of wash buffer. The protein was eluated in 0.2 mL fractions with wash buffer supplemented with 300 mM imidazole pH 8.0. The fractions containing the Ub-SUMO2 / SUMO2 mixture were pooled, concentrated and buffer exchanged into SEC Buffer (20 mM Tris pH 7.5, 100 mM NaCl) using Amicon centrifugal filter units (Millipore) with a 10 kDa MWCO. TEV protease was added to the concentrated and buffer exchanged Ub-SUMO/SUMO mixture for 1 h at room temperature. Finally, size-exclusion chromatography was performed with a Superdex S75 10/300 (Cytiva) in SEC buffer to separate Ub-SUMO2 from SUMO2. Fractions containing Ub-SUMO2 were pooled and concentrated. Protein concentration was calculated from the measured A280 absorption (extinction coefficients were calculated with ProtParam (https://web.expasy.org/protparam/)). Ub-SUMO2 was flash frozen using liquid nitrogen and stored at −80 °C until further use. Isolated yield calculated from SUMO2-K11LisoK starting material was 25.4 %. SDS-PAGE analysis of reaction progress and purification can be found in SI Figure 1.

#### 3.12 *In cellulo* UbyW cascade for ubiquitylation and neddylation of POIs

Chemically competent *E. coli* K12 cells were co-transformed with pBAD-POI-UBE2W (which encodes H6-tagged POI with a TAG codon at the denoted position and a UBE2W variant (see Supplementary Table S1) and pEVOL-Ubl-UBA1-aaRS (which encodes for ubiquitin (or linear di/tri/tetraUb or Nedd8), E1-activating enzyme and the Ma “IP” aaRS/tRNA pair for incorporation of LisoK/pLisoK, see Supplementary Table S1) plasmids.
After recovery with 1 mL of SOC medium for 1 h at 37 °C, the cells were cultured overnight in 5 mL of 2× YT medium containing ampicillin (100 µg/mL) and chloramphenicol (50 µg/mL) at 37 °C, 200 rpm. The overnight culture was diluted to an OD600 of approx. 0.05 in 100 mL autoinduction medium[10] supplemented with ampicillin (50 µg/mL), chloramphenicol (25 µg/mL) and GLisoK or GpLisoK (0.5 mM) and cultured at 37 °C with shaking (200 rpm) overnight. The cells were harvested by centrifugation (4000 × g, 10 min, 4 °C) and resuspended in lysis buffer (20 mM Tris pH 8.0, 300 mM NaCl, 30 mM Imidazole and 1 mM PMSF). Afterwards, the cell suspension was incubated on ice for 30 min and sonicated with cooling in an ice-water bath. The lysed cells were centrifuged (14,000 × g, 20 min, 4 °C) and the clear lysate was added to 1 mL Ni-NTA slurry/1 L culture (Cytiva) equilibrated with wash buffer (20 mM Tris pH 8.0, 300 mM NaCl, 30 mM imidazole). Afterwards, the mixture was incubated with agitation for 1 h at 4 °C. After incubation, the mixture was transferred to an empty plastic column and washed with 10 column volumes (CV) of wash buffer. The protein was eluated in 0.2 mL fractions with wash buffer supplemented with 300 mM imidazole pH 8.0. The fractions containing the Ub-POI/POI mixture were pooled and concentrated using Amicon centrifugal filter units (Millipore) with a suitable molecular weight cutoff (MWCO) followed by size-exclusion chromatography (SEC) using a Superdex S75 10/300 (Cytiva) with SEC buffer (20 mM Tris pH 7.5, 150 mM NaCl, 1 mM DTT). Fractions containing Ub-POI variants were pooled and concentrated. Protein concentration was calculated from the measured A280 absorption (extinction coefficients were calculated with ProtParam (https://web.expasy.org/protparam/)). Ubiquitylated POIs were flash frozen using liquid nitrogen and stored at −80 °C until further use. Facultative TEV cleavage was performed by adding TEV protease to the Ub-POI conjugate for 3 h at 4 °C followed by reverse Ni-NTA purification and SEC as described above.

This protocol was used for purification of Ub/diUb/triUb/tetraUb/N8/Ub(L73P)-SUMO2(K11LisoK) conjugates as well as for Ub/N8-PCNA. For Ub-SUMO1(K7LisoK) and GST-NEMO(K302/309LisoK) SEC was not performed.

For purification of Ub-Ran(K71LisoK) and Ub-Ran(K71pLisoK) purification buffers were adjusted accordingly (Lysis Buffer: 20 mM Tris pH 7.5, 300 mM NaCl, 20 mM Imidazole, 2 mM $MgCl_2$, 1 mM TCEP, 10% Glycerol; SEC Buffer: 20 mM Tris pH 7.5, 100 mM KCl, 2 mM $MgCl_2$, 1 mM TCEP, 10% Glycerol)

The following yields were obtained for SEC purified conjugates:

| | |
|---|---|
| Ub-SUMO2(K11pLisoK) | Yield: 3.1 mg/L |
| diUb-SUMO2(K11pLisoK) | Yield: 4.8 mg/L |
| triUb-SUMO2(K11LisoK) | Yield: 2.3 mg/L |
| tetraUb-SUMO2(K11LisoK) | Yield: 1.4 mg/L |
| Ub(L73P)-SUMO2(K11LisoK) | Yield: 6.8 mg/L |
| Nedd8-SUMO2(K11LisoK) | Yield: 1.6 mg/L |
| Ub-PCNA(K164LisoK) | Yield: 35.0 mg/L of 1:1 Ub-PCNA:PCNA |
| Ub-PCNA(K164pLisoK) | Yield: 21.3 mg/L of 1:1 Ub-PCNA:PCNA |
| N8-PCNA(K164pLisoK) | Yield: 11.5 mg/L of 1:3 N8-PCNA:PCNA |
| Ub-Ran(K71LisoK) | Yield: 1.0 mg/L |
| Ub-Ran(K71pLisoK) | Yield: 0.7 mg/L |

#### 3.13 Assembly of K63-linked diUbs

Assembly of K63-linked diUbs was carried out as previously described with slight adjustments.[11] In short, the assembly reactions contained 100 nM H6-UBE1 (Boston Biochem, Cat. No. E-304-050), 2.5 µM UBE2N, 2,5 µM UBE2V1, 1 mM MP-TEV-Ub wt. Reactions were incubated at 37 °C for typically 16 h in diUb reaction buffer (50 mM Tris pH 7.5, 10 mM MgCl2, 0.6 mM DTT, 10 mM ATP). After incubation, pH of the reaction was adjusted to 4.5 using 1 M ammonium acetate followed by centrifugation (21,000 × g, 5 min, 4 °C). Finally the reaction mixture was subjected to SEC using a Superdex S75 16/600 (Cytiva) with SEC Buffer (20 mM Hepes pH 7.8, 100 mM NaCl). Fractions containing the desired K63-linked diUb were pooled and concentrated using Amicon® centrifugal filter units with a 10 kDa MWCO (Millipore). Protein concentration was determined using BCA (Thermo Scientific™) and Bradford (Sigma–Aldrich) assays. K63-diUb was flash frozen using liquid nitrogen and stored at −80 °C until further use.

#### 3.14 Nucleotide loading of Ran

Nucleotide loading of Ran was performed as previously described.[12] Ub-Ran or Ran was diluted to 50 µM into exchange buffer (20 mM Tris pH 7.5, 100 mM KCl, 1 mM TCEP, 10 mM EDTA) followed by the addition of 1.5 mM of target nucleotide (e.g. GDP or Mant-GDP). After incubation for 3 h at RT 20 mM of $MgCl_2$ were added followed by SEC using a Superdex S75 16/600 (Cytiva) with SEC Buffer (20 mM Tris pH 7.5, 100 mM KCl, 2 mM $MgCl_2$, 1 mM TCEP, 10 % Glycerol). Nucleotide loaded Rans were flash frozen using liquid nitrogen and stored at −80 °C until further use.

### 4. Biochemical and *In Vitro* Assays

#### 4.1 *In vitro* USP2 cleavage assay

*In vitro* deubiquitylation assays were performed in DUB buffer (50 mM Tris, pH 7.5, 100 mM NaCl, 1 mM DTT). Ub-SUMO2(K11LisoK) or Ub(L73P)-SUMO2(K11LisoK) (10 μM) were incubated with the catalytic domain of USP2 (1 μM) at 37 °C. Reactions were initiated by the addition of the USP2 enzyme. At indicated time points samples were taken and immediately quenched with 4 × SDS sample buffer and analyzed by Coomassie stained SDS-PAGE.

#### 4.2 Conjugate stability in HEK293T cell lysate

HEK293T cells were lysed in buffer containing 1% (v/v) NP-40, 50 mM Tris-HCl (pH 8.0), 150 mM NaCl, and 1 mM TCEP. Lysis was assisted by sonication at 4 °C using a Bioruptor Plus (Diagenode) for 10 cycles (30 s ON / 30 s OFF). The crude lysate was cleared by centrifugation at 21,000 × *g* for 20 min at 4 °C. Purified Ub-SUMO2(K11LisoK) or Ub(L73P)-SUMO2(K11LisoK) conjugates were added to the freshly prepared, cleared lysate to a final concentration of 15 μM. Reactions were incubated on ice for 4 hours, either in the presence or absence of an inhibitor cocktail consisting of 1× cOmplete protease inhibitor (Roche) and 5 mM *N*-ethylmaleimide. Following incubation, the probes were isolated from the lysate via Ni-NTA affinity chromatography, SDS sample buffer was added, and samples were boiled at 95 °C for 10 min. Samples were subsequently resolved by SDS-PAGE and visualized by Coomassie staining.

#### 4.3 NTF2 pull-down assay

NTF2 pull down assays were performed by charging 2 μg of Ran-GDP, Ran-K71LisoK-GDP or Ub-Ran(K71LisoK)-GDP in 100 μL pull-down buffer (PDB: 20 mM Tris pH 7.5, 150 mM KCl, 2 mM MgCl2, 1 mM TCEP, 0.01% NP40) onto 1 μL washed magnetic Ni-NTA slurry (CubeBiotech) for 15 min at 4 °C. Afterwards excess of Ran was removed by washing the magnetic beads three times using a magnetic rack. NTF2 was diluted to 1 μM in PDB and added to the bead-bound Ran (50 μL/pull-down) for 30 min at 4 °C followed by three washing steps with PDB supplemented with 30 mM Imidazole. Elution was performed by adding 20 μL of PDB supplemented with 300 mM Imidazole followed by SDS-PAGE analysis.

#### 4.4 RCC1-mediated nucleotide exchange assays

RCC1-mediated nucleotide exchange assays were performed as previously described.[12] Mant-GDP loaded Ran and Ub-Ran(K71LisoK) were diluted to 700 nM in reaction buffer (20 mM Tris pH 7.5, 100 mM KCl, 5 mM MgCl2, 0.5 mM TCEP). Measurements were carried out in quintuplicates in 384-well plates (Greiner 781906) using a Tecan spark platereader monitoring fluorescence (ex: 355 nm, em: 448 nm). A 10 μL aliquot of 700 nM Mant-GDP Ran was added to each well, followed by injection of 10 μL of a mixture containing 10 nM RCC1 and 100 μM GDP using the injector. Fluorescence was monitored every 10 s for 15 min. All Data analysis was performed using GraphPad Prism v11.0.

### 5. Cell Lysate and Protein Mass Spectrometry Methods

#### 5.1 Cell Lysate Preparation, Photocrosslinking, and Affinity Purification

**Cell harvest and lysis** HEK293T cells were grown to 90% confluency in five 10 cm culture dishes. Cells were washed once with warm DPBS and harvested into 10 mL of cold DPBS. Following centrifugation (700 × g, 7 min, 4 °C), the supernatant was aspirated and the cell pellets were stored on ice. Each pellet was resuspended in 300 μL of cold lysis buffer (20 mM Tris pH 8.0, 150 mM NaCl, 2 mM $MgCl_2$, 0.1% NP-40, 1 mM TCEP, 10% glycerol) supplemented with 1× cOmplete protease inhibitor cocktail (Roche) and 5 mM iodoacetamide (IAA) to irreversibly inhibit deubiquitinase activity. To ensure comprehensive protein solubilization while maintaining weak protein-protein interactions, the resuspension was subjected to ultrasonication using a Bioruptor Plus (Diagenode) at 4 °C for 10 cycles (30 s ON / 30 s OFF). The crude lysate was cleared by centrifugation (21,000 × g, 20 min, 4 °C) and the supernatants were combined.

**In-lysate photocrosslinking** For photocrosslinking, the diazirine-containing probes Ran(K71pLisoK) or Ub-Ran(K71pLisoK) were added to 100 μL aliquots of the cleared HEK293T lysate to a final probe concentration of 500 nM. As a negative control, lysate without probe was processed in parallel. The mixtures were incubated for 15 min on ice to allow for interactor binding. Samples were subsequently transferred to a 24-well plate (100 μL per well) and irradiated on ice with UV light (365nm, 2000 mJ/cm²) using a UVP crosslinker (Analytik Jena).

**Affinity purification** Magnetic His-affinity beads (Cube Biotech, 1 μL slurry per sample) were equilibrated by washing once with 500 μL of wash buffer (50 mM Tris pH 8.0, 150 mM NaCl, 20 mM imidazole, 0.01% NP-40) supplemented with 300 mM imidazole, followed by a second wash with 500 μL of wash buffer. The UV-irradiated lysates were added to the equilibrated beads and incubated overnight at 4 °C on a rotating mixer (IKA Trayster basic).

Following incubation, the supernatant was removed using a magnetic rack. The beads were subjected to three washes. For each wash step, 500 μL of wash buffer was added, the samples were incubated for 4 min on the rotating mixer, and the beads were isolated for 2 min on the magnetic rack. The beads were then washed once with 500 μL of submission buffer (50 mM ammonium bicarbonate, 0.01% NP-40) under the same rotation/magnet conditions. Finally, the beads were resuspended in 50 μL of 50 mM ammonium bicarbonate, pH 7.8. for downstream proteomic sample preparation. To validate pulldown efficiency, parallel control samples (1 replicate per condition) were boiled in SDS sample buffer and evaluated by SDS-PAGE and Coomassie staining (see Fig. S19a).

*Note on lysis optimization:* Initial pulldown optimization utilizing recombinantly expressed RCC1 spike-ins indicated that high detergent concentrations disrupted the physiological interaction between the Ran probes and known interactors. Consequently, lysis and washing buffers were formulated with minimized detergent concentrations. To compensate for reduced detergent and prevent aggregation, strict mechanical solubilization via the Bioruptor was employed. Downstream proteomic analysis confirmed that this low-detergent, high-shear approach successfully maintained broad proteomic solubilization and coverage.

#### 5.2 LFQ-AP-MS Sample Preparation and Measurement

Captured proteins were digested directly on the affinity beads. Beads were washed twice with digestion buffer (10 mM Tris, 2 mM $CaCl_2$, pH 8.2). For reduction and alkylation, beads were resuspended in digestion buffer supplemented with Tris(2-carboxyethyl)phosphine (5 mM) and 2-chloroacetamide (15 mM). Following a 30-minute incubation in the dark at 30 °C, 500 ng of sequencing-grade modified trypsin (Promega, V5111) was added. Digestion proceeded overnight at 37 °C. Supernatants were collected, and residual peptides were extracted from the beads using 0.1% (v/v) trifluoroacetic acid (TFA) in 50% (v/v) acetonitrile (ACN). The respective supernatants and extracts were combined and dried under vacuum. Prior to analysis, peptides were resuspended in 20 µL of LC-MS loading buffer (3% ACN, 0.1% formic acid [FA]) containing iRT standard peptides.

Peptides were analyzed by LC-MS/MS on an Orbitrap Exploris 480 mass spectrometer (Thermo Fisher Scientific) coupled to a Waters ACQUITY UPLC M-Class system configured for 75-µm single-pump trapping. Chromatographic separation was performed on a nanoEase HSS C18 T3 column (75 µm × 250 mm, 100 Å; Waters, 186008818) maintained at 50 °C. The mobile phase consisted of 0.1% (v/v) formic acid in water (Solvent A) and 0.1% (v/v) formic acid in acetonitrile (Solvent B). Peptides were eluted at a flow rate of 300 nL/min using a piecewise linear gradient from 5% to 33% Solvent B over 45 min

Data-independent acquisition (DIA) was performed using 70 non-overlapping 10 *m/z* windows spanning an *m/z* range of 350 to 1050. Full MS1 scans were acquired at a resolution of 120,000. Precursors were quadrupole-isolated and fragmented via higher-energy collisional dissociation (HCD) at a normalized collision energy (NCE) of 28, assuming a default charge state of 3. MS2 spectra were acquired at a resolution of 30,000, with a normalized AGC target of 3000% and the maximum injection time set to automatic.

### 5.3 Tandem Mass Spectrometry for Modification-site Determination

Recombinant proteins were resuspended in 50 µL 6 M urea in 50 mM ammonium bicarbonate and incubated at room temperature for 30 min. Reduction of disulfide bonds was performed by addition 0.25 µL of 500 mM TCEP (final concentration 2.5 mM) and incubating for 30 min at 37 °C, 800 rpm. Cysteine alkylation was carried out by addition of 0.5 µL freshly prepared 500 mM iodoacetamide (IAA, final concentration 5 mM) and incubating for 30 min at room temperature in the dark. For enzymatic digestion, the sample was diluted by addition of 23 µL of 150 mM $NH_4HCO_3$ (pH 7.8) and then incubated with 200 ng of Lys-C (Promega) for 2 h at 37 °C, 600 rpm. 330 µL of 50 mM $NH_4HCO_3$ (pH 7.8) was added and the sample was subsequently incubated with 400 ng of trypsin (Promega) overnight at 37 °C, 600 rpm. Digested peptide samples were purified, desalted, and concentrated using C18 spin columns (Thermo Fisher Scientific) according to the manufacturers protocol. The desalted eluate was dried using a speed vacuum concentrator before its resuspension in 25 µL of 0.1% formic acid. High-resolution Tandem MS spectra were obtained at the Molecular and Biomolecular Analysis Service (MoBiAS) of ETH Zürich Department of Chemistry and Applied Biosciences. An ESI-TIMS-QTOF-MS system (timsTOF Pro, Bruker Daltonics, Germany) equipped with collision- induced dissociation (CID) using $N_2$ as the collision gas was used. Tryptic peptides were pressure-loaded onto a 25 cm × 75 µm i.d. C18, 1.6 µm column (Ionoptics Ltd., Australia) with a 5 mm × 0.3 mm i.d. C18, 5 µm column (Thermo Scientific, Lithuania) serving as a guard column, maintained at 40 °C. The mobile phase consisted of (A) water with 0.1% formic acid and (B) acetonitrile with 0.1% formic acid. The gradient was initiated at 2% B, linearly increasing to 35% B over 120 min, then further increasing to 95% B in 2 min, all at a flow rate of 300 nL $min^{-1}$. This was followed by isocratic conditions of 95% B for 8 min. The total run time, including column conditioning, was 163 min.

### 5.4 Intact protein Mass Spectrometry

**ESI-LC-MS** LC-MS analysis of full-length proteins was performed on an Agilent 1260 Infinity Series LC system with an Agilent 6210 ESI Single Quadrupole mass spectrometer using a Jupiter C4 column (2 mm, 150 mm, 300 Å, 5 µm) capillary column (Phenomenex, Torrence, USA). The analysis was performed at RT with a flow rate of 900 µL/min and a gradient of 10% - 55% solvent B in 1.65 min, followed by a gradient of 55 %-90 % solvent B in 0.85 min (solvent A: 0.1 % FA in water, solvent B 0.1 % FA in ACN). Data generated on the Agilent LC-MS was analyzed with OpenLab ChemStation (Agilent).

### 6. Computational and Bioinformatics Analysis

#### 6.1 Data Processing for LFQ-AP-MS

Raw LC-MS/MS data were processed using DIA-NN (v1.9.2) in library-free mode with the match-between-runs feature enabled. A predicted spectral library was generated from the UniProt human reference proteome (UP000005640). To prevent ambiguous peptide assignments and redundant protein group identifications, the sequences of multiple ubiquitin-encoding genes (*UBB*, *UBC*, *UBA52*, and *RPS27A*) were consolidated into a single ubiquitin consensus entry. The *in silico* digest was configured for fully tryptic peptides ranging from 6 to 30 amino acids, allowing zero missed cleavages. Carbamidomethylation of cysteine was set as a fixed modification. Methionine oxidation (UniMod:35) was set as a variable modification. Precursor charge states were restricted to 2-3, with precursor and fragment m/z ranges set to 350-1050 and 200-1800, respectively. Protein and peptide identifications were filtered at a 1% false discovery rate (FDR).

Statistical analysis and differential expression analysis (DEA) were performed in R (v4.5.1) using the *prolfqua* package (v1.4.0) and prolfquapp (v2.0.10).[7] Protein abundance estimates were extracted from the DIA-NN main report, filtering for proteins quantified by a minimum of two unique peptides. Robust scaling (robscale) was applied to the log2-transformed intensities. Normalization was validated by visual inspection of kernel density estimates of protein log2-intensities across samples, confirming that robust scaling appropriately aligned the distributions without dampening biological variance typical in AP-MS datasets, (Supplementary Fig. S18b).

Two samples (Ub-Ran_4 and bead_1) were identified as outliers based on unbiased quality control metrics, specifically missingness analysis and protein abundance heatmaps and were subsequently excluded. The removal of these outliers decreased the overall coefficient of variation (CV), the primary interactors, USP15 and RCC1, remain the highest-ranked statistically significant hits regardless of outlier inclusion.

Data exploration and visualization were conducted using a *exploreDE* (v7) based application. Volcano and MA plots were generated without missing value imputation, applying a log2 fold-change threshold of ±0.5 and an FDR-adjusted p-value threshold of <0.05. The PCA pairs plot (Supplementary Fig. S18e) was generated using the normalized data of the top 2000 features ranked by standard deviation. The sample correlation heatmap (Supplementary Fig. S19a) was rendered with a diverging scale limit of 4.

Manual inspection of specific protein hits and validation of peptide-level evidence were performed using Scaffold DIA (v3.2.1, Proteome Software). This software was specifically employed to verify the identification of USP15 and to determine the number of unique peptides associated with this hit (Supplementary Fig. S19d).

#### 6.2 Data Processing for Modification-site validation

Raw MS/MS data were processed and searched using the Mascot search engine (Matrix Science, v3.1).[3] Spectra were searched against a consolidated database comprising the UniProt *Homo sapiens* reference proteome and a custom database containing the recombinant target protein sequences. Search parameters were configured as follows: trypsin was specified as the protease, allowing for up to two missed cleavages; precursor and fragment mass tolerances were both set to ± 20 ppm. Carbamidomethylation of cysteine was set as a fixed modification. Variable modifications included oxidation of methionine, deamidation of asparagine and glutamine, and the site-specific footprint of the isopeptide-linked ubiquitin remnant on lysine residues (GGLisoK, Δmass = +228.1348 Da). Peptide Identifications were filtered at a 1% FDR. Modification sites were confirmed by manual inspection of the high-resolution MS/MS spectra to ensure unambiguous *b*- and *y*-ion coverage surrounding the modified residue, (Fig. 1e).

#### 6.3 Ubi-score Analysis

Analysis and visualizations were performed using R (v4.3.1) in VSCode (v1.99.3). The ubiquitylation site positional importance score for all human ubiquitylation sites was extracted from Table S6, *van Gerwen et al.*.[13] The score of RAN K71 was plotted compared to all other ubiquitylation sites using the ggplot2 package (v3.4.2). Quantitative fold change values of RAN K71 in di-glycine proteomics data were extracted from *van Gerwen et al.*.[13] Fold change values from conditions comprising translation inhibition, protein folding stress, UV treatment, and proteasome inhibition were extracted and plotted using the ggplot2 package.

**Supplementary Table S2. Regulation of Ran(K71).** Summarized Log2 fold change values for RAN K71 site in selected di-glycine proteomics experiments, as reported by *van Gerwen et al.*[13] Columns: *uniprot_site*: Site identifier for RAN K71, *log2FC*: log2 fold change value comparing an experimental condition to a control condition, *Condition group*: Type of experimental condition, *Description*: Description of the experimental condition and control condition, Treatment: Description of the experimental condition, *Time (min)*: Duration of the experimental condition, *Control*: Description of the control condition *Pubmed*: Pubmed identifier for datasets published before *van Gerwen et al.*[13] *Dataset ID*: Identifier for the individual dataset the condition came from. The identifier matches Table S3 from *van Gerwen et al.*[13]

| uniprot_ site | log2FC | Condition group | Description | Treatment | Time (min) | Control | Pubmed | Dataset ID |
|---|---|---|---|---|---|---|---|---|
| P62826_ K71 | 1.21753 | DNA damage | UV+Recovery_vs_Control_time(min)_120 | UV+Recovery | 120 | Control | | unpublished_Homo sapiens_8 |
| P62826_ K71 | 0.994981 | Protein folding stress | Canavanine_vs_Control_time(min)_120 | Canavanine | 120 | Control | | unpublished_Homo sapiens_1 |
| P62826_ K71 | 0.975975 | Translation inhibition | Bortezomib 1um + cycloheximide_vs_Bortezomib 1um_time(min)_480 | Bortezomib 1um + cycloheximide | 480 | Bortezomib 1um | 21906983 | published_Homo sapiens_2 |
| P62826_ K71 | 0.851191 | Proteasome inhibition | Bortezomib 1um_vs_DMSO_time(min)_480 | Bortezomib 1um | 480 | DMSO | 21906983 | published_Homo sapiens_2 |
| P62826_ K71 | 0.823195 | Translation inhibition | Cycloheximide_vs_DMSO_time(min)_480 | Cycloheximide | 480 | DMSO | 21906983 | published_Homo sapiens_2 |
| P62826_ K71 | 0.778068 | Protein folding stress | AZC_vs_Control_time(min)_240 | AZC | 240 | Control | | unpublished_Homo sapiens_2 |
| P62826_ K71 | 0.738641 | Proteasome inhibition | Epoxomicin 5nm_vs_Control_time(min)_480 | Epoxomicin 5nm | 480 | Control | 27185884 | published_Homo sapiens_14 |
| P62826_ K71 | 0.616008 | DNA damage | UV+Recovery_vs_Control_time(min)_15 | UV+Recovery | 15 | Control | | unpublished_Homo sapiens_8 |
| P62826_ K71 | 0.317813 | Proteasome inhibition | Epoxomicin 1um_vs_DMSO_time(min)_480 | Epoxomicin 1um | 480 | DMSO | 21906983 | published_Homo sapiens_2 |
| P62826_ K71 | 0.314586 | DNA damage | UV radiation_vs_Control_time(min)_NA | UV radiation | | Control | 26051181 | published_Homo sapiens_11 |
| P62826_ K71 | 0.304022 | DNA damage | UV+Recovery_vs_Control_time(min)_60 | UV+Recovery | 60 | Control | | unpublished_Homo sapiens_8 |
| P62826_ K71 | 0.279504 | DNA damage | UV+Recovery_vs_Control_time(min)_480 | UV+Recovery | 480 | Control | | unpublished_Homo sapiens_8 |
| P62826_ K71 | 0.031958 | DNA damage | UV radiation_vs_Control_time(min)_NA | UV radiation | | Control | 26051181 | published_Homo sapiens_11 |
| P62826_ K71 | 0.010307 | Protein folding stress | AZC_vs_Control_time(min)_120 | AZC | 120 | Control | | unpublished_Homo sapiens_2 |
| P62826_ K71 | -0.08795 | Proteasome inhibition | Epoxomicin 10nm_vs_Control_time(min)_480 | Epoxomicin 10nm | 480 | Control | 27185884 | published_Homo sapiens_14 |
| P62826_ K71 | -0.25587 | Proteasome inhibition | Epoxomicin 500nm_vs_Control_time(min)_480 | Epoxomicin 500nm | 480 | Control | 27185884 | published_Homo sapiens_14 |
| P62826_ K71 | -0.36664 | Proteasome inhibition | Epoxomicin 20um_vs_Control_time(min)_480 | Epoxomicin 20um | 480 | Control | 26051182 | published_Homo sapiens_12 |
| P62826_ K71 | -0.52144 | Proteasome inhibition | Epoxomicin 500nm_vs_Control_time(min)_480 | Epoxomicin 500nm | 480 | Control | | unpublished_Homo sapiens_5 |
| P62826_ K71 | -0.5891 | Proteasome inhibition | Epoxomicin 25nm_vs_Control_time(min)_480 | Epoxomicin 25nm | 480 | Control | 27185884 | published_Homo sapiens_14 |
| P62826_ K71 | -0.89145 | Proteasome inhibition | Bortezomib 1um_vs_DMSO_time(min)_480 | Bortezomib 1um | 480 | DMSO | 21906983 | published_Homo sapiens_2 |

| | | | | | | | | |
|---|---|---|---|---|---|---|---|---|
| P62826_K71 | -1.68675 | Proteasome inhibition | Bortezomib 1um_vs_DMSO_time(min)_480 | Bortezomib 1um | 480 | DMSO | 21906983 | published_Homo sapiens_2 |

### 6.4 UBE2W Homology Alignment

**Structural Homology Analysis** Structural similarity searches were conducted using the Foldseek web server (search.foldseek.com).[6] The structure of human UBE2W (PDB: 2MT6) was used as the query against the AlphaFold Protein Structure Database and the Protein Data Bank (PDB100). Alignments were computed using the server's default 3Di/AA algorithm with an E-value cutoff of 0.001. Foldseek scores are reported in Fig. S15a.

**Multiple Sequence Alignment** Amino acid sequences of the identified UBE2W homologs were retrieved from UniProt. Multiple sequence alignment was performed using the T-Coffee web server (v13.46.0) via the EMBL-EBI Job Dispatcher sequence analysis framework. The percent identity matrix comparing the homologs was calculated using the Clustal 2.1 algorithm integrated within the T-Coffee pipeline. The resulting MSA and identity scores are provided in Supplementary Fig. S15.

### 7. Small molecule characterization

### 7.1 LC-MS

LC-MS analysis of small molecules was performed on an Agilent 1260 Infinity Series LC system with an Agilent 6210 ESI Single Quadrupole mass spectrometer using a Luna® C18(2)-HST (2 mm, 100 mm, 100 Å, 2.5 µm) capillary column (Phenomenex, Torrence, USA). The analysis was performed at RT with a flow rate of 360 µL/min and a gradient of 15-95 % solvent B in 3.5 min (solvent A: 0.1 % formic acid (FA) in water, solvent B 0.1 % FA in acetonitrile (ACN)).

### 7.2 HR-MS

For HR-MS analysis samples were submitted to the Molecular and Biomolecular Analysis Service MoBiAS (ETH Zurich) and analyzed on a Bruker Daltonics maXis ESI-QTOF.

#### 7.3 NMR

$^1$H spectra were measured on Bruker Avance III HD 300 MHz (300 MHz for $^1$H-NMR), Bruker Avance III 400 UltraShield (400 MHz for $^1$H-NMR), or Bruker Avance-I AV500 UltraShield (500 MHz for $^1$H-NMR). The spectra were analyzed using MestReNova, Version 14.3.1-31739 (Mestrelab Research S.L.). The spectra were referenced to the respective solvent to determine the chemical shift (δ) in ppm (DMSO$d_6$: 2.54, $CD_3OD$: 3.34, $CDCl_3$: 7.26547. The coupling constants are given in Hertz (Hz) and signal multiplicity is characterized as follows: s (singlet), d (doublet), t (triplet), q (quartet), qint (quintet), m (multiplet), br (broad), and combinations thereof.

### Data Availability

All mass spectrometry proteomics data generated in this study will be deposited to the ProteomeXchange Consortium via the PRIDE partner repository upon publication.

### Author Contributions

M.F and K.L. conceived the research plan and experimental strategy. M.F. synthesized ncAAs, purified all proteins and conjugates, performed in vitro UbyW assays, set up and optimized the in cellulo cascade, conducted Ran nucleotide exchange assays as well as NTF2 pull-downs. P.S. performed MS-MS analysis of Ub-SUMO conjugate, conducted Ran proteomics and data evaluation and performed the USP15 DUB assay. J.v.G. calculated positional importance score for Ran ubiquitylation sites. D.K. synthesized GpLisoK. F.W. helped with initial in vitro experiments and cloning. All authors analysed the data, and P.S. wrote the paper with input from the other authors.

### Acknowledgements

This work was supported by funding from ETH Zurich and the European Research Council (ERC under the European Union’s Horizon 2020 research and innovation program, grant agreement no. 101003289—Ubl-tool to K.L.). The authors gratefully acknowledge the Functional Genomics Center Zurich (FGCZ) of University of Zurich and ETH Zurich, and particularly Dr. Tobias Kockmann, for the support on proteomics analyses. The authors acknowledge the staff of the MoBiAS MS service (ETH Zürich) for assistance with the tandem mass spectrometry for modification site determination. We also thank Lang group members for useful discussions and input.
